# S-Palmitoylation stabilizes OGT and the OGT-PPP1CC complex

**DOI:** 10.64898/2026.08.29.747956

**Authors:** Xiaoxuan Lu, Ting Xu, Jie Li, Yihan Liu, Wen Zhou, Kai Wang, Chang Niu, Ni Tang, Leiliang Zhang, Jing Li

**Author notes:** To whom correspondence should be addressed: **Jing Li,** Beijing Key Laboratory of DNA Damage Response, College of Life Sciences, Capital Normal University, Beijing 100048, China, **Leiliang Zhang,** Department of Pathogen Biology, School of Clinical and Basic Medical Sciences, Shandong First Medical University & Shandong Academy of Medical Sciences, Jinan, Shandong 250117, China, **Ni Tang,** Key Laboratory of Molecular Biology for Infectious Diseases (Ministry of Education), Institute for Viral Hepatitis, Department of Infectious Diseases, The Second Affiliated Hospital, Chongqing Medical University, Chongqing 400010, China, **Chang Niu,** Beijing Key Laboratory of DNA Damage Response, College of Life Sciences, Capital Normal University, Beijing 100048, China. These authors contributed equally to this work (Xiaoxuan Lu, Ting Xu, Jie Li).

## Abstract

O-linked β-N-acetylglucosamine (O-GlcNAc) transferase (OGT) is the sole writer for intracellular O-GlcNAcylation. It catalyzes O-GlcNAcylation of thousands of protein substrates, but relatively less is known about the post-translational modifications that occur on OGT itself. Herein, we demonstrate that OGT is S-palmitoylated at Cys-472 and Cys-477, which is mediated by the S-acyltransferase Zinc Finger DHHC-Type Palmitoyl transferase 14 (zDHHC14) and removed by acyl protein thioesterase 2 (APT2). S-Palmitoylation stabilizes OGT by shunting it away from the lysosomal chaperone-mediated autophagy (CMA) pathway, as S-palmitoylation decreases the interaction between OGT and heat shock cognate 70 kDa protein (HSC70), the CMA chaperone. *Via* label-free quantitative mass spectrometry, we find that S-palmitoylation elevates the affinity between OGT and protein phosphatase 1 catalytic subunit gamma (PPP1CC), but not PPP1CB. We further demonstrate that S-palmitoylation of OGT augments binding with Yes-associated protein-1 (YAP), a protein that associates with PPP1CC, and subsequently enhances YAP O-GlcNAcylation. Our work unearths S-palmitoylation of OGT and CMA-mediated degradation of lysosomal OGT, the orchestration of which finetunes the activity of key OGT complexes, such as OGT-PPP1CC, and contributes to OGT substrate selectivity.

## INTRODUCTION

*O*-linked *N*-acetylglucosamine (*O*-GlcNAc) transferase (OGT) catalyzes the intracellular O-GlcNAcylation reactions that modulate cellular signaling (1,2), including cell division (3), transcription (4), immunology (5) and stress response (6). Its impairment is closely associated with human diseases, such as diabetes (7), Alzheimer’s disease and cancer (8). The recent profiling of O-GlcNAc substrates has revealed more than 7 000 proteins, but they do not share any conserved motif, suggesting that regulation of the O-GlcNAcome may be more complicated than we originally thought. Post-translational modifications (PTMs) on OGT itself may provide an inroad to monitor and finetune its substrates.

Work from the last 4 decades has shown that OGT is regulated by many PTMs. OGT itself is O-GlcNAcylated at S389 to regulate its localization (9) and is ubiquitinated at K352 to prime for ubiquitin-mediated mitotic degradation (10). In terms of phosphorylation, OGT is phosphorylated by Glycogen synthase kinase 3β (GSK3β) (11), by calcium/calmodulin-dependent kinase II (CaMKII) and Checkpoint kinase 1 (Chk1) and at S20 (12) (13), by AMP-activated protein kinase (AMPK) at T444 for chromatin association (14,15), by phosphatidylinositol 3-kinase β (PI3Kβ) at T985 to enhance OGT activity (16) and by ULK1 at S576 for stabilization (17). Its phosphorylation is also stimulated by Epidermal growth factor (EGF) at Y976 (18). These PTMs regulate OGT stability, protein-protein interaction, localization and stability. As a result, OGT is shown to localize to the nucleus, cytosol, midbody during cytokinesis (12), plasma membrane upon serum stimulation (19), mitochondria (20) and recently the lysosome (21).

The lysosome not only senses cellular nutrimental status, but also mediates decay of cellular wastes to maintain cellular homeostasis (22). Autophagy mediates the disposal process and it comes in three forms: macroautophagy, microautophagy and chaperone-mediated autophagy (CMA) (23). CMA substrates first bind to heat shock cognate 70 kDa protein (HSC70), a cytosolic chaperone, then they are translocated into the lysosome by lysosomal-associated membrane protein 2A (LAMP2A) on the lysosomal membrane (24). Failures in CMA lead to not only protein quality control defects, but also dysfunction of DNA damage response, metabolic reprogramming and stress response (24). Recently, O-GlcNAcase (OGA), the eraser for O-GlcNAcylation, is shown to be degraded to CMA (25).

It is currently unknown whether OGT is subject to any lipidation modifications. As one form of lipidation, S-palmitoylation involves covalently linking the fatty acid palmitate to the Cys residues of protein substrates *via* a thioester bond (26) (27). Due to its lability, S-palmitoylation is highly dynamic: it is catalyzed by S-acyltransferase Zinc Finger DHHC-Type Palmitoyl transferases (zDHHCs) and reversed by de-palmitoylases (28). By altering the stability and localization of the substrates, S-palmitoylation thus allows steering the protein function in a hydrophobic setting (28).

In this work, we present evidence that OGT is S-palmitoylated. Through bioinformatic prediction and biochemical validation, we found two major S-palmitoylated residues, C472/C477. We further identify zDHHC14 as one of the writers and acyl protein thioesterase 2 (APT2) as the eraser. Moreover, we show that OGT is subject to CMA, and S-palmitoylation blocks the CMA process. Downstream of OGT S-palmitoylation, we identify that it promotes the stability of the OGT-protein phosphatase 1 catalytic subunit gamma (PPP1CC) complex and Yes-associated protein-1 (YAP) O-GlcNAcylation. Our work thus reveals how lipidation regulates OGT stability and offers new insights into the crosstalk between S-palmitoylation and OGT.

## RESULTS

### OGT is palmitoylated at C472 and C477

As S-palmitoylation is known to respond to lipid metabolism, we speculate that OGT could respond to lipid and be modified by S-palmitoylation. To test this possibility, co-immunoprecipitation (coIP) coupled with acyl-biotinyl exchange (ABE) assay was carried out, and we found that ectopic OGT was palmitoylated, but OGA was not (Fig. 1A). To map the potential S-palmitoylation sites, we used the following websites: http://biocuckoo.org (for Cys-472) and http://proteininformatics.org/mkumar/palmpred/index.html (for Cys-477) (29). We generated the single and double mutants (C472S/C477S, 2CS) accordingly, which were then subject to ABE assays (Fig.1B-C). The OGT-2CS mutant attenuated S-palmitoylation significantly (Fig.1B-C), but not the single mutants (Supplementary Figure S1). Thus, OGT is palmitoylated at two major conserved Cys residues, which are quite conserved among species (Fig. 1D). As the 2CS mutant still displays S-palmitoylation to some extent, it is highly probable that there are still other S-palmitoylation sites.

**Figure 1.**
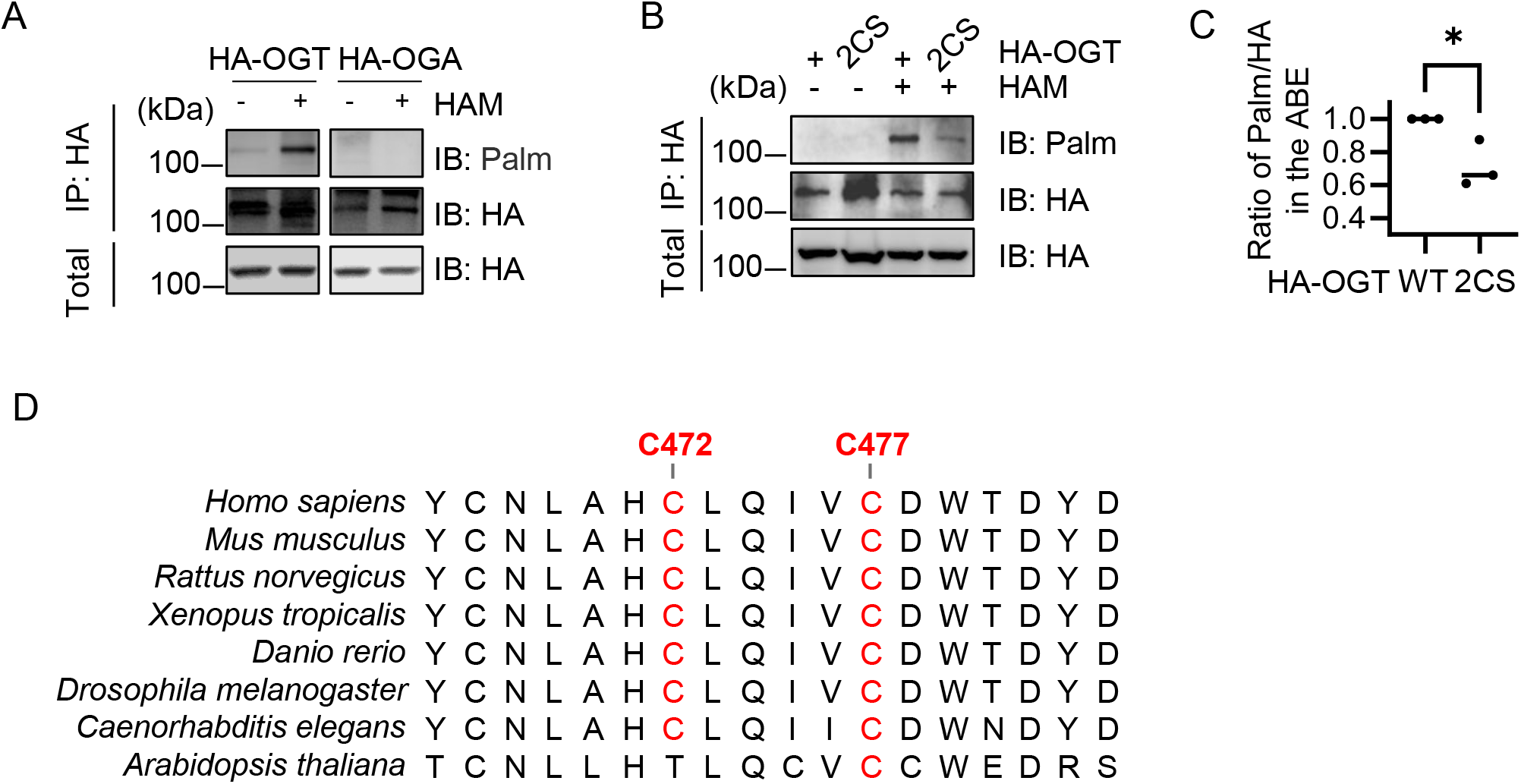
OGT is S-palmitoylation at C472/C477. A. 293T cells were transfected with HA-OGT and HA-OGA plasmids, and the cell lysates were subject to immunoprecipitation and acyl-biotin exchange (ABE) experiments using the Streptavidin-HRP antibodies and then immunoprecipitated with anti-HA antibodies, as described in experimental procedures. B. 293T cells were transfected with HA-OGT and HA-OGT-C472S/C477S (2CS) plasmids, and the cell lysates were subject to immunoprecipitation and ABE experiments. C. Quantitation of (B). The statistical analysis in (C) was presented as mean ± SD from n = 3 biologically independent experiments. Statistical significance was determined by two-tailed unpaired Student’s t-test, *P < 0.01. All Western blots were performed for at least three times. D. Sequence alignments of OGT palmitoylated cysteine residues across different species.

### OGT associates with zDHHC14

As protein S-palmitoylation is catalyzed by the 23 zinc finger DHHC-type containing zDHHC) family members (26), we screened for potential zDHHC members that interact with OGT (Supplementary Figure S2), and zDHHC14 shows robust binding affinity with OGT reciprocally (Fig. 2A-B). When endogenous affinity was examined, zDHHC14 was shown to coIP with OGT (Figure 2C). These findings suggest that OGT binds with zDHHC14.

**Figure 2.**
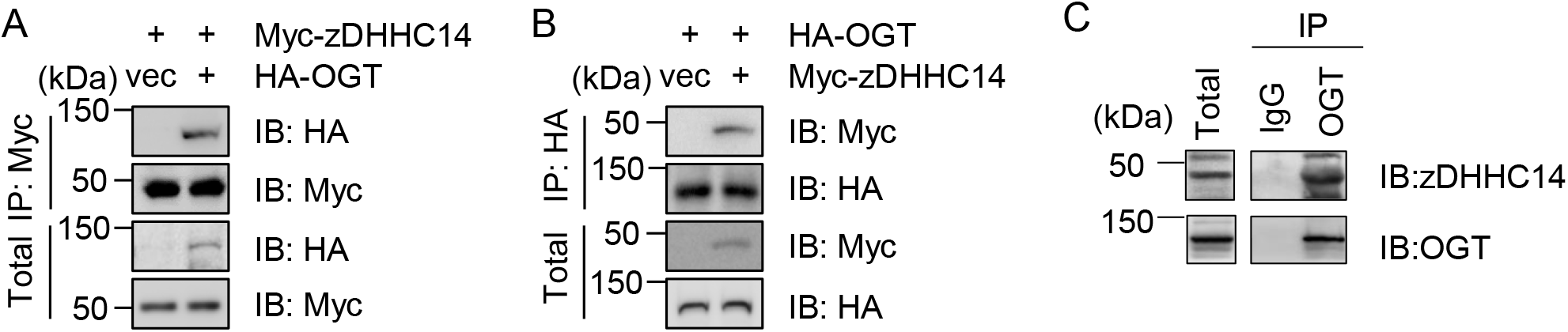
zDHHC14 associates with OGT. A-B, Reciprocal co-immunoprecipitation between exogenous zDHHC14-Flag and HA-OGT proteins. 293T cells were transfected with zDHHC14-Flag and HA-OGT plasmids, and the cell lysates were subject to immunoprecipitation and immunoblotting experiments using the antibodies indicated. C. Cellular lysates were immunoprecipitated with anti-OGT antibodies and the immunoprecipitates were immunoblotted with the antibodies indicated.

### APT2 modulates OGT de-S**-**palmitoylation

It is known that the de-S-palmitoylation reaction is mediated by APT1-2, and APT1 mainly localizes in the mitochondria (30). We thus tested the interaction between OGT and APT2 (Fig. 3A-B). APT2 shows a robust association with OGT. We then analyzed the endogenous interaction between APT2 and OGT (Fig. 3C), and found the two proteins associate with each other.

**Figure 3.**
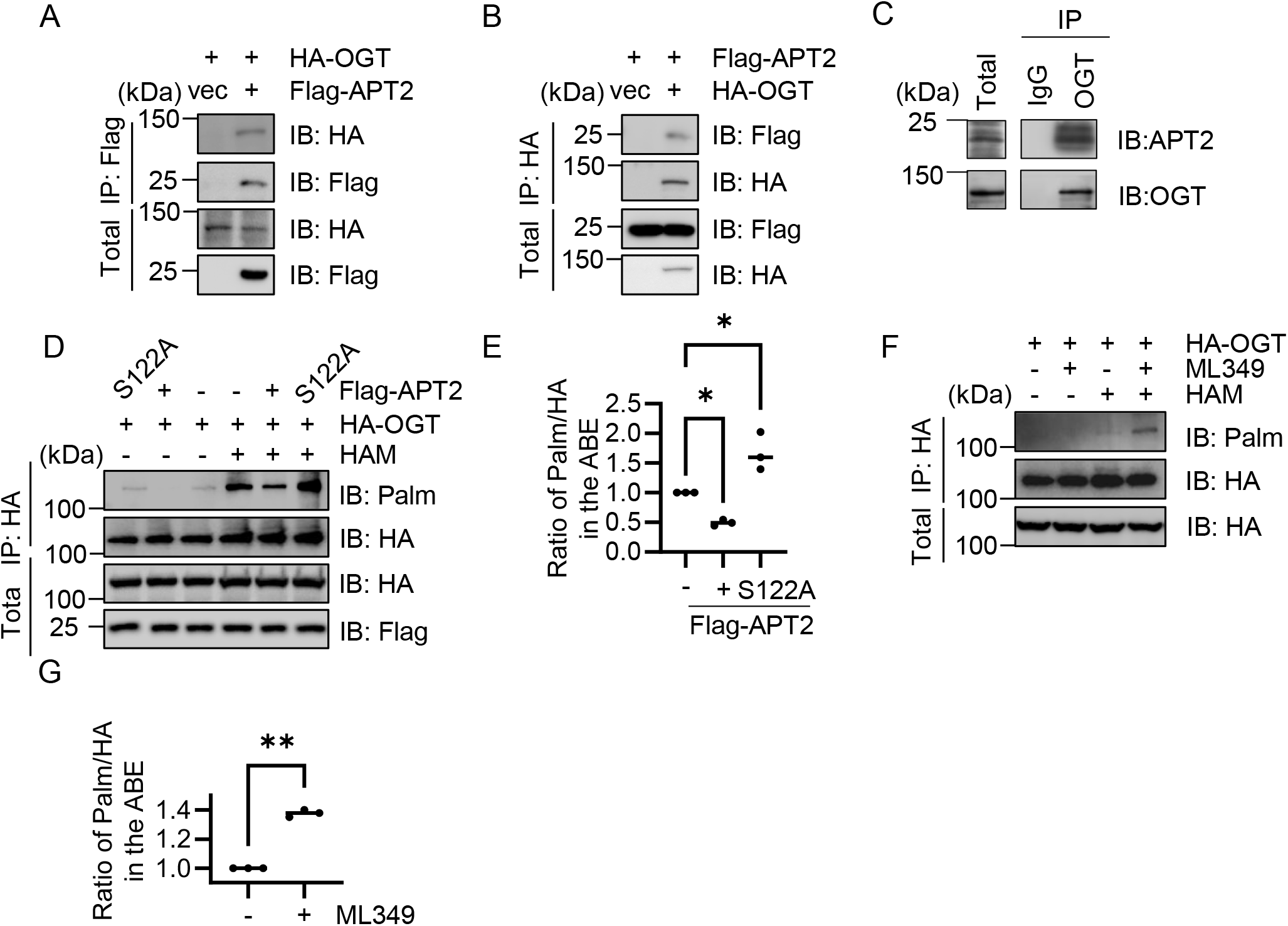
APT2 de-S-palmitoylates OGT. A-B Co-immunoprecipitation between exogenous Flag-APT2 and HA-OGT proteins. 293T cells were transfected with Flag-APT2 and HA-OGT plasmids, and the cell lysates were subject to immunoprecipitation and immunoblotting experiments using the antibodies indicated. C. Cellular lysates were immunoprecipitated with anti-OGT antibodies and the immunoprecipitates were immunoblotted with the antibodies indicated. D. 293 cells were transfected with Flag-APT2-WT or APT2-S122A plasmids and together with HA-OGT, and the cell lysates were subject to immunoprecipitation and ABE assays. E. Quantitation of D. The statistical analysis in (E) was presented as mean ± SD from n = 3 biologically independent experiments. Statistical significance was determined by one-way ANOVA followed by Sidak’s multiple comparisons test. (\**P* < 0.01). F. Cells were transfected with HA-OGT plasmids, then treated with ML349 or mock treated, and the cell lysates were subject to immunoprecipitation and ABE experiments. G. Quantitation of F. The statistical analysis in (G)was performed as mean ± SD from n = 3 biologically independent experiments. Statistical significance was determined by two-tailed unpaired Student’s t-test. (** *P* < 0.001).

We further used Flag-APT2 in the ABE assay, and found that APT2 significantly attenuated OGT S-palmitoylation, but the enzymatic-dead APT2-S122A mutant did not (Fig. 3D-E). We also used an APT2-specific inhibitor, ML349. ML349 significantly increased the S-palmitoylation levels of OGT in ABE assays (Fig. 3F-G). Taken together, these results suggest that APT2 modulates the de-S-palmitoylation of OGT.

### OGT S**-**palmitoylation shunts OGT away from chaperone-mediated autophagy

As S-palmitoylation has been shown to crosstalk with the CMA pathway (31) and OGT has been demonstrated to localize to the lysosome (21), we wondered whether S-palmitoylation also exerts the same effect on OGT. To test this, we set out to examine whether OGT is regulated by CMA. Interestingly, the interaction between OGT and HSC70 has been detected in several proteomic studies (32–34) and we also identified endogenous coIP between HSC70 and OGT (Fig 4A). Two siRNAs targeting *HSC70* were used, and OGT levels were discernably higher compared to the control (Fig. 4B-C). Then we tested the affinity between OGT-2CS and HSC70 and found that 2CS elevated HSC70 binding robustly (Fig. 4D-G).

**Figure 4.**
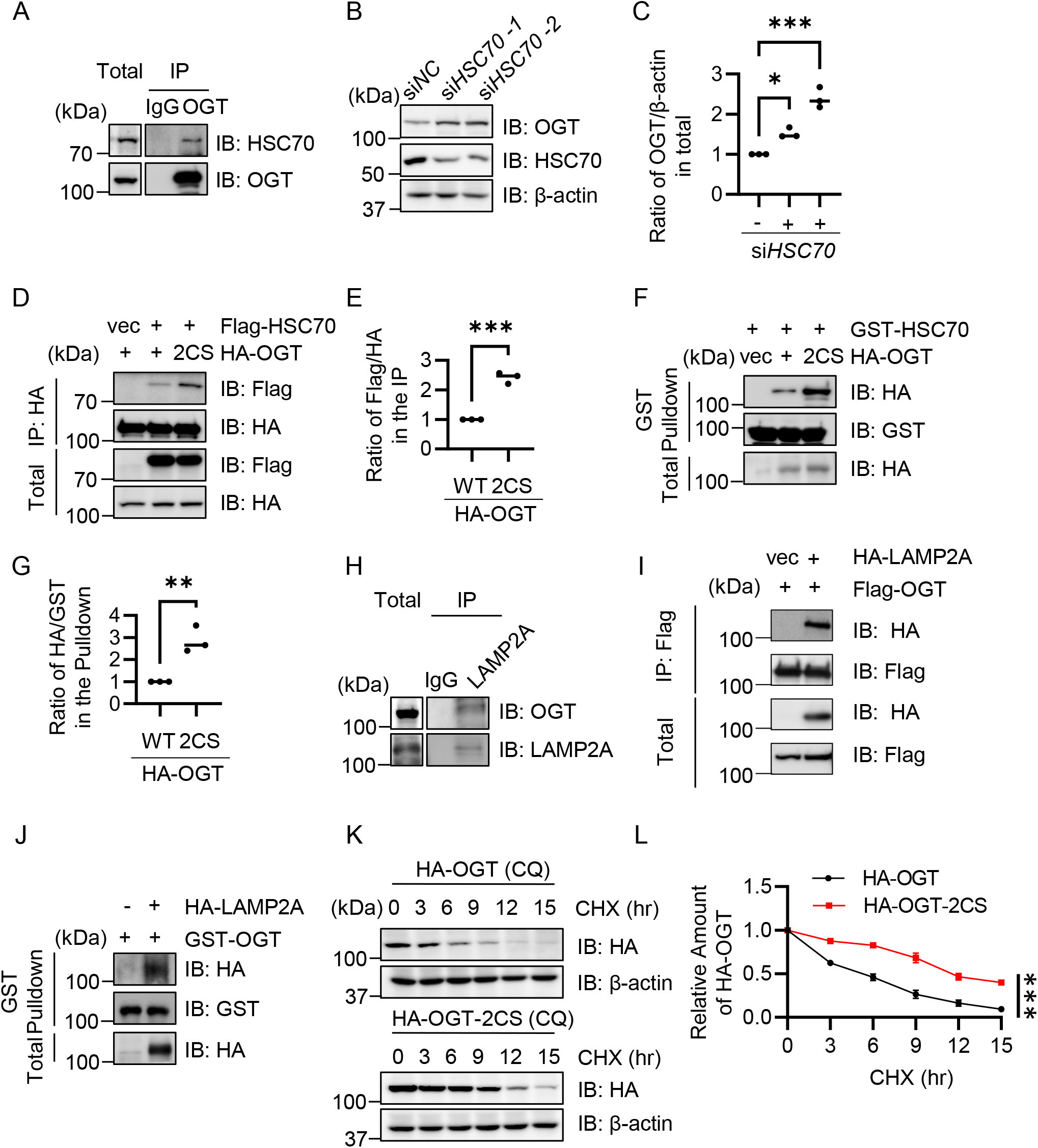
OGT S-palmitoylation inhibits degradation by chaperone-mediated autophagy (CMA). A. Cellular lysates were subject to immunoprecipitation by anti-OGT antibodies and immunoblotted with the antibodies indicated. B. Cells were treated with two independent siRNAs targeting *HSC70,* then the cell lysates were collected and immunoblotted with the antibodies indicated. C. Quantitation of (B). The statistical analysis in (C) was presented as mean ± SD from n = 3 biologically independent experiments. Statistical significance was determined by one-way ANOVA followed by Sidak’s multiple comparisons test. (\**P* < 0.05, *** *P* < 0.001). D. Cells were transfected with Flag-HSC70, together with HA-OGT-WT, or −2CS plasmids, and then the lysates were immunoprecipitated and immunoblotted with the antibodies indicated. E. Quantitation of (D).The statistical analysis in (E) was presented as mean ± SD from n = 3 biologically independent experiments. Statistical significance was determined by two-tailed unpaired Student’s t-test. (*** *P* < 0.001).F. Cells were transfected with HA-OGT-WT, −2CS plasmids, and the lysates were incubated with recombinant GST-HSC70 proteins. Then GST-pulldown experiments were carried out. G. Quantitation of (F). The statistical analysis in (G) was presented as mean ± SD from n = 3 biologically independent experiments. Statistical significance was determined by two-tailed unpaired Student’s t-test. (** *P* < 0.01). H. Cellular lysates were subject to immunoprecipitation by anti-LAMP2A antibodies and immunoblotted with the antibodies indicated. I. Cells were transfected with Flag-LAMP2A and HA-OGT plasmids. J. Cells were transfected with HA-LAMP2A plasmids, and the cellular lysates were incubated with recombinant GST-OGT proteins and GST-pulldown experiments were carried out. K. Cells were transfected with HA-OGT-WT or −2CS plasmids, and then treated with chloroquine (CQ) and Cycloheximide (CHX). L. Quantitation of (K). The statistical analysis in (L) was presented as mean ± SD from n = 3 biologically independent experiments. Statistical significance was determined by two-way ANOVA with Sidak’s multiple comparisons test.

We further validated the CMA regulation of OGT by assessing the interaction between OGT and Lamp2a (Fig. 4H-I), and found that both endogenous and exogenous OGT interact with Lamp2a. GST-OGT also could pulldown Lamp2a, further implicating OGT in the CMA pathway (Fig. 4J). Last, we used chloroquine (CQ) and cycloheximide (CHX) to examine OGT half-life (Fig. 4K-L) and found that OGT-2CS is stabler than WT, suggesting that dampening the CMA pathway increases OGT abundance. Taken together, OGT degradation is partly regulated by CMA, and S-palmitoylation shunts OGT away from CMA.

### OGT S**-**palmitoylation increases OGT-PPP1CC binding

We wondered what is downstream of OGT S-palmitoylation. To this end, a label-free quantitative mass spectrometry assay was carried out. When the interactome was compared between OGT and OGT-2CS, we found that significantly less PPP1CC binds with OGT-2CS (Supplementary Table S1-2). Endogenous OGT binds with PPP1CC (Fig. 5A), consistent with previous findings that OGT interacts with PPP1CB and PPP1CC but not PPP1CA (35). In an coIP assay, OGT-2CS displayed less binding with PPP1CC (Fig. 5B-C). Also, in a GST pulldown assay, OGT-WT showed more affinity with PPP1CC (Fig. 5D-E), which are consistent with the mass spectrometry results. When PPP1CB was examined, OGT-2CS has no discernable effects (Supplementary Figure S3.), suggesting that OGT S-palmitoylation specifically modulates OGT-PPP1CC stability.

**Figure 5.**
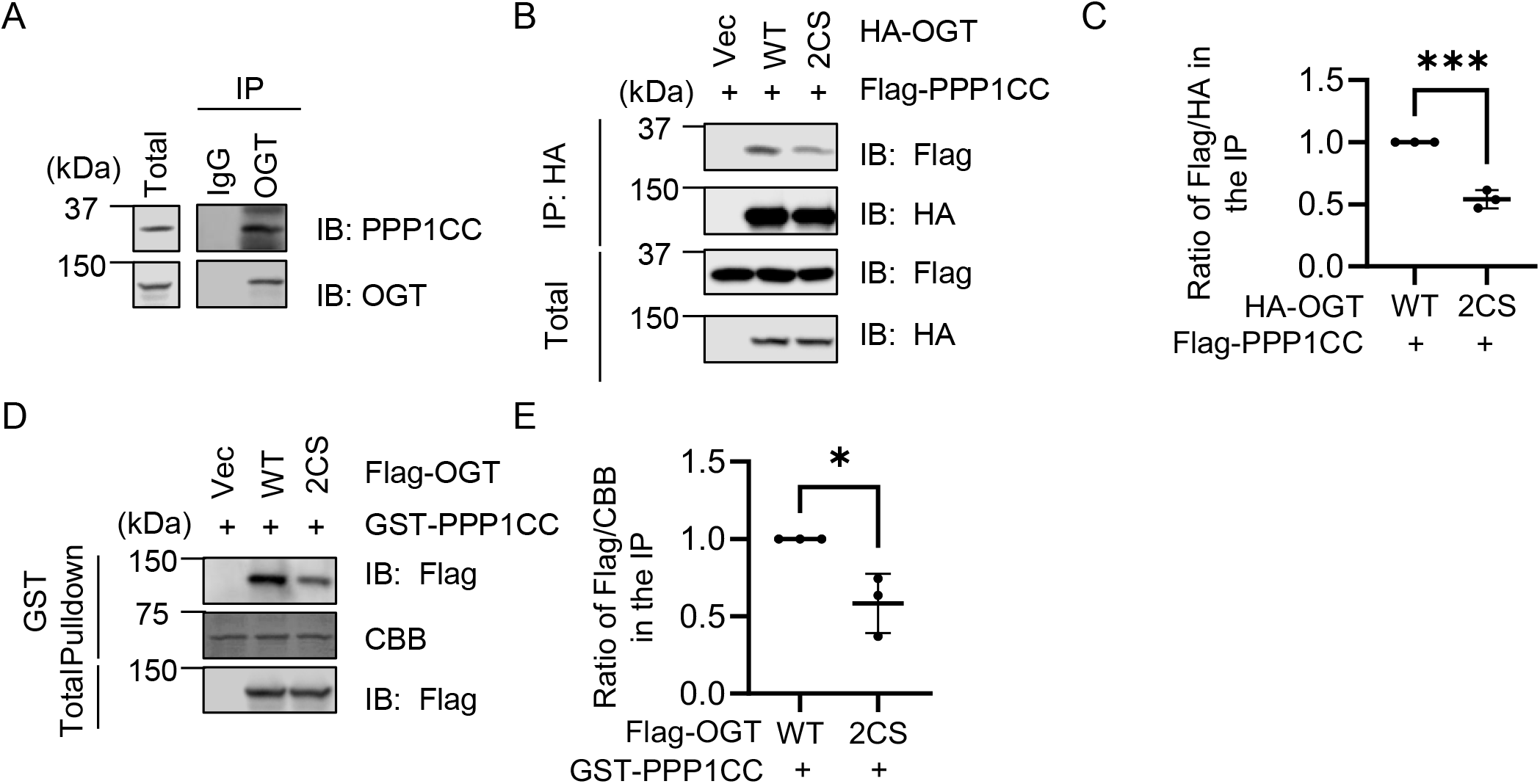
S-Palmitoylation stabilizes the OGT-PPP1CC complex. A, Cellular lysates were immunoprecipitated with anti-OGT antibodies and the immunoprecipitates were immunoblotted with the antibodies indicated. B, Co-immunoprecipitation between exogenous Flag-PPP1CC and HA-OGT proteins. HEK293T cells were transfected with HA-Vec, HA-OGT and HA-OGT-2CS together with Flag-PPP1CC plasmids and the cell lysates were subject to immunoprecipitation and immunoblotting experiments using the antibodies indicated. C, Quantitation of (B). D, Cells were transfected with Flag-Vec, Flag-OGT-WT, Flag-OGT-2CS plasmids, and the lysates were incubated with recombinant GST-PPP1CC proteins. And then GST-pulldown experiments were carried out. E, Quantitation of (D). The statistical analysis in (C, E) was performed as mean ± SD from n = 3 biologically independent experiments. Statistical significance was determined by two-tailed unpaired Student’s t-test. (\**P* < 0.01, *** *P* < 0.0001).

### OGT S**-**palmitoylation upregulates YAP O-GlcNAcylation

Since YAP has been shown to associate with PPP1CC (36) and OGT (37), we tested whether PPP1CC upregulates the interaction between OGT and YAP. As shown in Fig. 6A-B, PPP1CC overproduction significantly elevated the association between YAP and OGT. When OGT-WT and −2CS were co-transfected with YAP plasmids, OGT-2CS displayed significantly less binding with YAP (Fig. 6C-D). Consequently, upon OGT-2CS transfection, YAP reduced much O-GlcNAcylation (Fig. 6E-F) and increased phosphorylation (Fig. 6G-H), consistent with previous findings that YAP O-GlcNAcylation antagonizes phosphorylation (37). Thus, we found that O-GlcNAcylation of one target of PPP1CC, YAP, is regulated by OGT S-palmitoylation. It is possible that other PPP1CC partners are regulated in the same manner.

**Figure 6.**
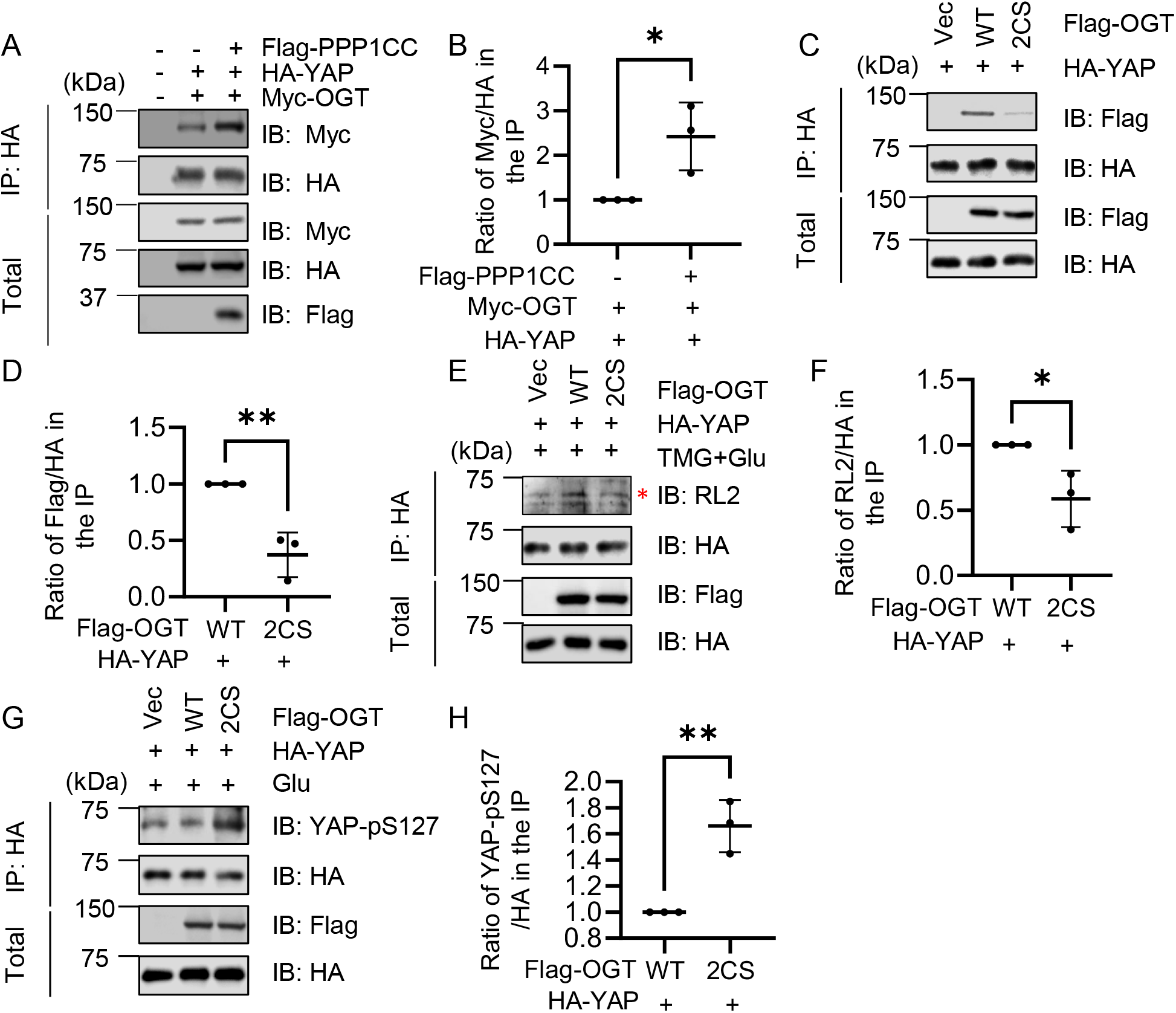
OGT S-palmitoylation increases YAP O-GlcNAcylation. A, Cells were transfected with HA-YAP, Myc-OGT and Flag-PPP1CC plasmids. B, quantitation of (A). C, HEK293T cells were transfected with vector, Flag-OGT-WT and −2CS together with HA-YAP plasmids and then the lysates were immunoprecipitated and immunoblotted with the antibodies indicated. D, Quantitation of (C). E, HEK293T cells were transfected with Flag-Vec, Flag-OGT-WT and Flag-OGT-2CS together with HA-YAP plasmids and then treated with 0.25 mM Thiamet-G (TMG) for 24 h, and 30 mM Glucose (Glu) for 3 h. Cells were then subjected to immunoprecipitation and immunoblotting with indicated antibodies. F, Quantitation of (E). G, HEK293T cells were transfected with vector, Flag-OGT-WT and −2CS together with HA-YAP plasmids and then treated with Glucose (Glu) for 3 h, and then the lysates were immunoprecipitated and immunoblotted with the antibodies indicated. H, Quantitation of (G). The statistical analysis was performed using Student’s t-test The statistical analysis in (B, D and F) was performed as mean ± SD from n = 3 biologically independent experiments. Statistical significance was determined by two-tailed unpaired Student’s t-test. (\**P* < 0.01, ** *P* < 0.001).

## DISCUSSION

In this work, we demonstrated that OGT is S-palmitoylated at Cys-472 and Cys-477, which shunts it away from CMA in the lysosome. S-palmitoylation promotes the affinity between OGT and PPP1CC and may regulate the O-GlcNAcylation of some PPP1CC downstream targets, such as YAP (Fig. 7). We note that most of our experiments were performed in HEK293T/293 cells. Validation of the molecular mechanism in physiologically relevant cell types or disease models will be further needed to establish general relevance.

**Figure 7.**
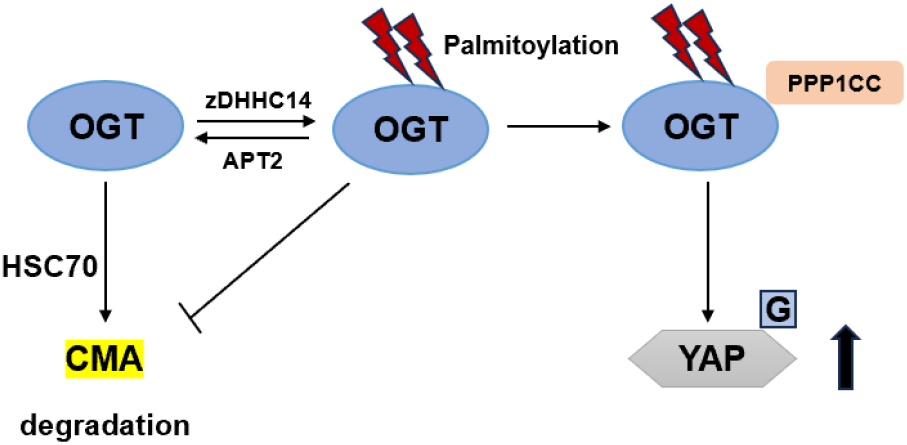
A model showing the role of OGT S-palmitoylation. We propose that OGT is palmitoylated, which stabilizes OGT by inhibiting its HSC70-mediated CMA degradation, enhances the affinity between OGT and PPP1CC, and subsequent O-GlcNAcylation of target proteins, such as YAP.

It is not known whether OGT is subject to other lipidation processes, such as N-myristoylation or S-prenylation. Other lipid modifications, such as glycosylphosphatidylinositol (GPI) anchor and cholesterylation, occur in the lumen of secretory organelles (27). But OGT has been reported to glycosylate GRASP55, the Golgi stacking protein, to promote autophagosome-lysosome fusion (38), suggesting that OGT could potentially localize to the secretory organelles. Perhaps OGT is potentially subject to other lipidation forms. It is also worth mentioning that ABE assay detects reversible S-acylation, so OGT may be modified by other forms of S-acylation.

We found that Cys-472 and Cys-477 are S-palmitoylated. Our results do not exclude the possibility that OGT harbors other S-palmitoylation sites. First, in our ABE assay, S-palmitoylation is still discernable in the 2CS mutant. Second, S-palmitoylation could occur on other sites if different cell lines were used or distinct biological stimuli were applied. Third, many proteins are known to have multiple S-palmitoylation sites, such as NOD-like receptor pyrin domain-containing protein 3 (NLRP3). NLRP3 is palmitoylated at Cys-898 by Toll-like receptor (TLR) ligation for translocation to dispersed trans-Golgi network (39). It is also modified at Cys-126, which is inhibited by disulfiram, an FDA-approved drug (40). Therefore, there could be other S-palmitoylation sites on OGT.

OGT has been shown to localize to the lysosome, where it O-GlcNAcylates lysosomal proteins, such as Cathepsin B (21). We show in this work that OGT is subject to lysosomal degradation *via* the CMA pathway, and S-palmitoylation inhibits CMA. On the contrary, the NLRP3 inflammasome is recently shown to be S-palmitoylated, and subsequently shunted to CMA for degradation (31). It will be worthwhile to explore the relationship between S-palmitoylation and CMA on other proteins.

The relationship between OGT and protein phosphatases are largely unexplored. Previously, OGT has been shown to bind PPP1CB and PPP1CC (35). Later, OGT and OGA were shown to co-purify together with PPP1C during M phase of the cell cycle (41)(PPP1CA/B/C were not differentiated in this study). Our work demonstrates that OGT S-palmitoylation specifically regulates the binding between OGT and PPP1CC, and subsequently O-GlcNAcylation of PPP1CC targets, such as YAP. The S-palmitoylation sites we identified are located in the intrinsically disordered region of OGT. Currently, it is difficult to predict its effects on the OGT structure. But it may alter OGT localization, such as the case of NLRP3 (39). It is a possible scenario that other OGT PTMs regulate OGT-PPP1CB affinity and exert effects on PPP1CB substrates. The variety of OGT PTMs thus confers specificity on OGT interactome and OGT substrates.

## EXPERIMENTAL PROCEDURES

### Cell culture, antibodies and plasmids

All cells were purchased from COBIOER (CBP60232, Nanjing, China). The cell lines were validated using STR profiling and free from mycoplasma contamination for all experiments. The antibodies were: anti-Flag (XHY021L, Beijing, China), anti-HA (Proteintech,51064-2-AP, Chicago, USA), HRP-conjugated Affinipure Goat anti Rabbit IgG(H+L) (Proteintech, SA00001-2, Chicago, USA), Streptavidin-HRP (Genscript, M00091, USA), goat anti-rabbit Alexa Fluor 594 (Invitrogen, A-11037, USA), goat anti-mouse Alexa Fluor 647 (Invitrogen, A-21235, USA), HSC70 (Abcam, # ab19136); β-actin (Immunoway, #YM3028), OGT (Abcam, # ab96718), LAMP2A (Abcam, # ab18528), RL2 (Abcam, # ab2739); zDHHC4 (Invitrogen, #PA5-98601), APT2 (Abcam, #AB151578), PPP1CC (ABclonal, #A21818), Phospho-YAP (Ser127) (CST, #13008). siRNA sequences targeting *HSC70* are: GAACAAGAGAGCTGTAAGA, GTGCCATGACAAAGGATAA.

### IP and immunoblotting

The immunoprecipitation and immunoblotting assays were described before (42). The following primary antibodies were used for IB: anti-HA (1:3000), anti-Flag (1:1000), anti-Hsc70 (1:4000), anti-OGT (1:2000), anti-LAMP2A (1:1000), anti-RL2 (1:1000), anti-ZDHHC14 (1:2000), anti-APT2 (1:1000), anti-PPP1CC (1:1000), anti-Phospho-YAP (Ser127) (1:1000). The ECL detection system (Beyotiome, P0018FM-1, P0018FM-1, China) was used for immunoblotting. Bio-Rad ChemiDoc Toch was employed to detect signals, and the signals were quantitated by the Image J software. All western blots were repeated for at least three times.

### Chemical treatment

Chloroquine (CQ) was used at 10 µM for 2 h. ML349 (MCE, New Jersey, USA) was used at 8 µM for 18 h after transfected with indicated plasmids for 6 h. Thiamet-G (TMG) was used at 0.25 mM for 24 h. Glucose was used at 30 mM for 3 h.

### Acyl-Biotin Exchange (ABE) assay

After immunoprecipitation (IP) with the anti-HA antibodies, we carried out the Acyl-Biotin Exchange (ABE) assay by irreversibly blocking unmodified cysteine thiol groups using N-ethylmaliemide (NEM) for 30 min. Then the beads were washed and incubated with hydroxylamine (HAM) for 1 h at room temperature for specific cleavage and unmasking of the palmitoylated cysteine’s thiol group. Each group was divided into two parts, one with HAM (+ HAM), and the other without HAM (-HAM). After washing the beads were selectively labeled with a thiol-reactive biotinylation reagent, biotin-BMCC (ProteoChem, OC3930, USA) for 1 h at room temperature. Then Western Blotting was carried out to directly measure S-palmitoylation levels.

### Label-free quantitative mass spectrometry

#### Sample Preparation

The precipitated protein pellets were fully dissolved in 8 M urea, incubated in 10 mM DTT in 50 mM ammonium bicarbonate at 37°C for 45 min, then incubated in 10 mM iodoacetamide in 50 mM ammonium bicarbonate at ambient temperature for 1 hr in the dark. The solution was then diluted to a urea concentration of 2 M using 50 mM ammonium bicarbonate, followed by trypsin digestion with an enzyme:protein ratio of 1:40 at 37 °C overnight. Formic acid was added to the solution to a final concentration of 0.1% to quench the digestion. All samples were vacuum-centrifuged to dryness, and resuspended in 0.1% formic acid in water prior to LC-MS/MS analysis.

#### LC-MS/MS Parameters

For LC-MS/MS analysis, the samples were reconstituted in 0.2% formic acid, loaded onto a 100μm x 2 cm pre-column and separated on a 75μm x 15 cm capillary column with laser-pulled sprayer. Both columns were packed in-house with Luna 3 μm C18(2) bulk packing material (Phenomenex, USA). An Easy nLC 1000 system (Thermo Scientific, USA) was used to deliver the following HPLC gradient: 5-35% B in 60 min, 35-75% B in 4 min, then held at 75% B for 10 min (A = 0.1% formic acid in water, B = 0.1% formic acid in acetonitrile) at a flow rate of 300 nL/min. The eluted peptides were sprayed into a Velos Pro Orbitrap Elite mass spectrometer (Thermo Scientific, USA) equipped with a nano-ESI source. The mass spectrometer was operated in data-dependent mode with a full MS scan (375–1600 M/z) in FT mode at a resolution of 120000 followed by CID (Collision Induced Dissociation) MS/MS scans on the 10 most abundant ions in the initial MS scan. Automatic gain control (AGC) targets were 1e6 ions for Orbitrap scans and 5e4 for MS/MS scans. For dynamic exclusion, the following parameters were used: isolation window, 2 m/z; repeat count, 1; repeat duration, 25 s; and exclusion duration, 25 s.

#### Data Analysis

Data processing was carried out using Thermo Proteome Discoverer 2.4 using a Swissprot Human Database (version 2017-10-25). Carbamidomethyl (Cys) were chosen as static modification, and oxidation (Met) was chosen as variable modification. Mass tolerance was 10 ppm for precursor ions and 0.6 Da for fragment ions. Maximum missed cleavages was set as 2. Peptide spectral matches (PSM) were validated using the Percolator algorithm, based on q-values at a 1% FDR. The protein abundances were calculated by summing sample abundances of the corresponding peptides after normalization.

## Data Availability

The mass spectrometry proteomics data have been deposited to the ProteomeXchange Consortium via the PRIDE (43) partner repository with the dataset identifier PXD079515 and 10.6019/PXD079515.

## Abbreviations

IP: Immunoprecipitation
IB: Immunoblotting
TMG: Thiamet-G
MS: Mass spectrometry
O-GlcNAc: O-linked β-N-acetylglucosamine
OGT: O-GlcNAc transferase
PTM: post-translational modification
OGA: O-GlcNAcase
HSC70: heat shock cognate 70 kDa protein
CMA: chaperone-mediated autophagy
CQ: chloroquine
CHX: cycloheximide
ERK: Extracellular signal-regulated kinase
zDHHC: zinc finger DHHC-type containing
APT: Acyl protein thioesterase
ABE: acyl-biotinyl exchange
HAM: hydroxylamine

## Acknowledgements

We thank Dr. Xing Chen (Peking Univ.) for support. Jing L. is supported by the National Natural Science Foundation of China (NSFC) fund (92478113 and 32271285) and the Yunnan Province Academician Expert Workstation Project (202605AF350042). L.Z. is supported by NSFC (82272306 and 82072270), and Taishan Scholars Program (TSTP20221142)

