## Supplementary Figures and Tables for "S-Palmitoylation stabilizes OGT and the OGT-PPP1CC complex"

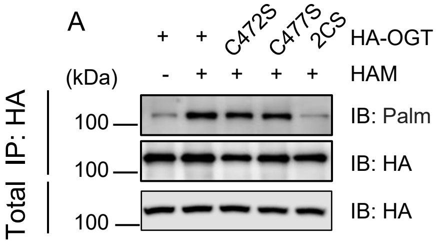

**Supplementary Figure S1. OGT is S-palmitoylated at C472 and C477.** 293T cells were transfected with HA-OGT, HA-OGT-C472S, HA-OGT-C477S or HA-OGT-2CS plasmids.The cell lysates were subject to immunoprecipitation and acyl-biotin exchange (ABE) experiments.

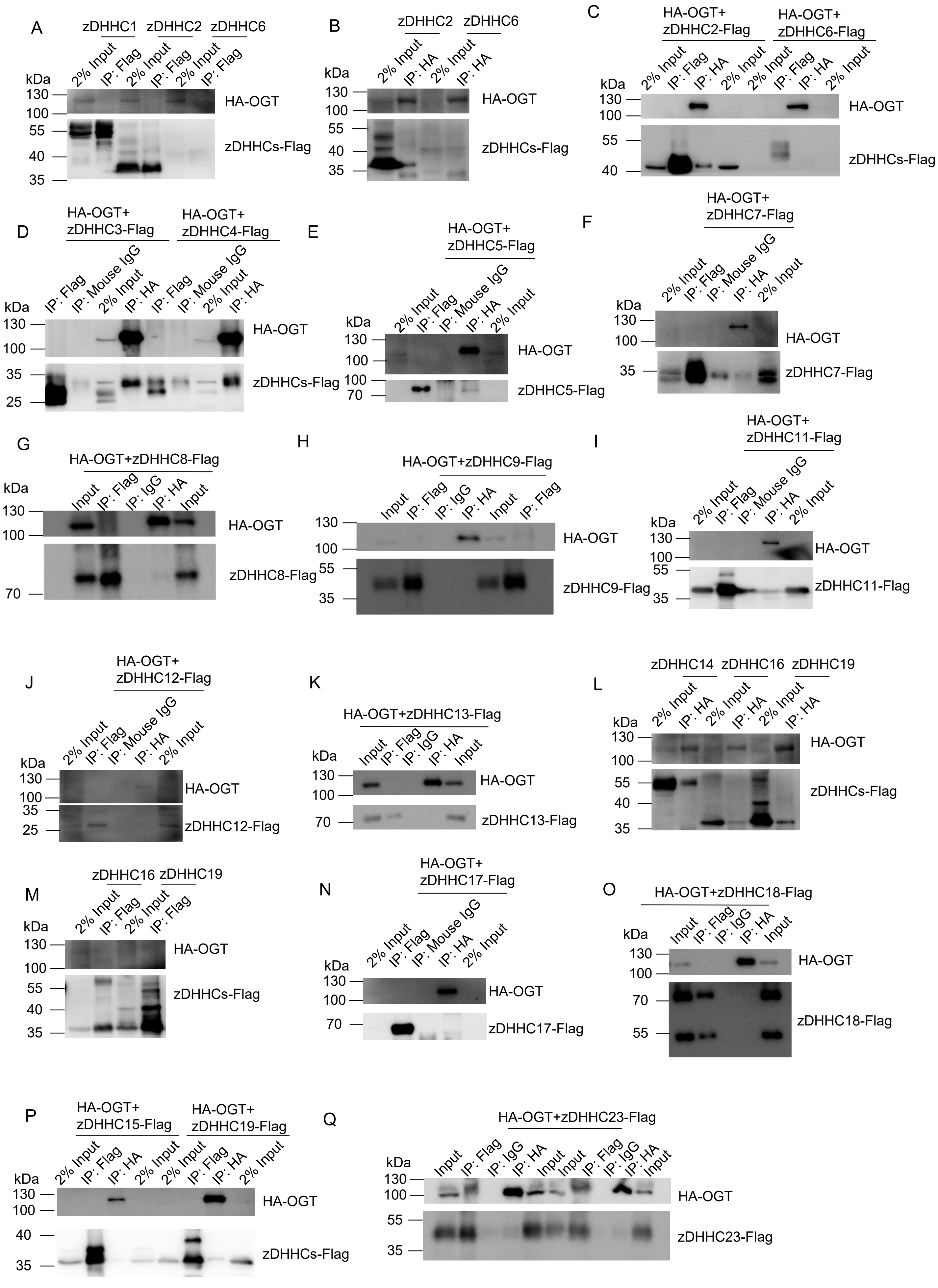

A
zDHHC1
zDHHC2
zDHHC6
2% Input
IP: Flag
kDa
130
100
55
40
35
HA-OGT
zDHHCs-Flag
B
zDHHC2
zDHHC6
2% Input
IP: HA
HA-OGT
zDHHCs-Flag
C
HA-OGT+
zDHHC2-Flag
HA-OGT+
zDHHC6-Flag
2% Input
IP: Flag
IP: HA
HA-OGT
zDHHCs-Flag
D
HA-OGT+
zDHHC3-Flag
HA-OGT+
zDHHC4-Flag
IP: Flag
IP: Mouse IgG
2% Input
IP: HA
HA-OGT
zDHHCs-Flag
E
HA-OGT+
zDHHC5-Flag
HA-OGT
zDHHC5-Flag
F
HA-OGT+
zDHHC7-Flag
HA-OGT
zDHHC7-Flag
G
HA-OGT+zDHHC8-Flag
Input
IP: Flag
IP: IgG
IP: HA
HA-OGT
zDHHC8-Flag
H
HA-OGT+zDHHC9-Flag
HA-OGT
zDHHC9-Flag
I
HA-OGT+
zDHHC11-Flag
HA-OGT
zDHHC11-Flag
J
HA-OGT+
zDHHC12-Flag
HA-OGT
zDHHC12-Flag
K
HA-OGT+zDHHC13-Flag
HA-OGT
zDHHC13-Flag
L
zDHHC14
zDHHC16
zDHHC19
HA-OGT
zDHHCs-Flag
M
zDHHC16
zDHHC19
HA-OGT
zDHHCs-Flag
N
HA-OGT+
zDHHC17-Flag
HA-OGT
zDHHC17-Flag
O
HA-OGT+zDHHC18-Flag
HA-OGT
zDHHC18-Flag
P
HA-OGT+
zDHHC15-Flag
HA-OGT+
zDHHC19-Flag
HA-OGT
zDHHCs-Flag
Q
HA-OGT+zDHHC23-Flag
HA-OGT
zDHHC23-Flag

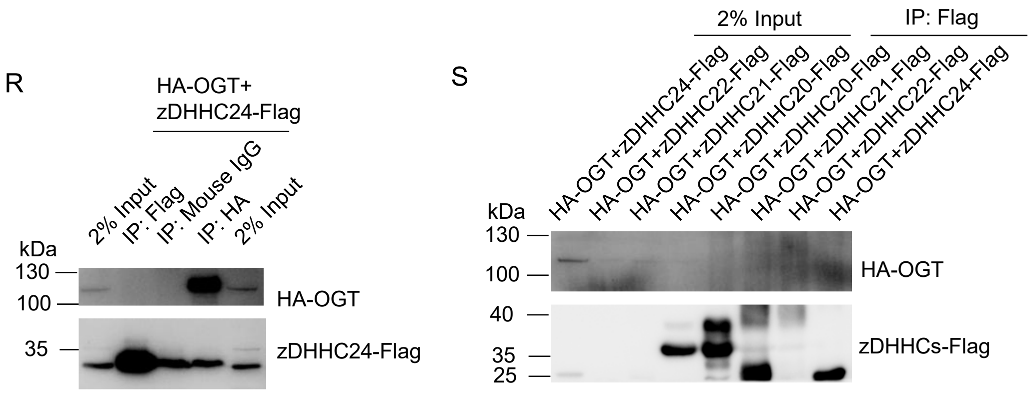

**Supplementary Figure S2. zDHHC14 interacts with OGT.** Protein palmitoylation is catalyzed by members of the zDHHC family. Here, we screened for potential zDHHC interactors of OGT and found that zDHHC14 strongly interacts with OGT.

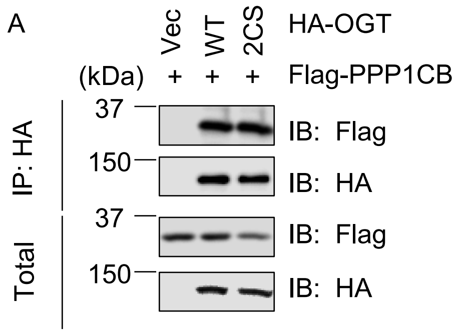

**Supplementary Figure S3.** S-Palmitoylation does not promote the interaction between OGT and PPP1CB. HEK293T cells were transfected with HA-Vec, HA -OGT-WT and HA -OGT-2CS together with Flag-PPP1CC plasmids and then the lysates were immunoprecipitated and immunoblotted with the antibodies indicated.

Table S1 Label-free quantitative mass spectrometry reveals an OGT interactome

| # Protein Pathway Groups | Accession | Description | Coverage [%] | # Peptides | # PSMs | # Unique Peptides | # AAs | MW [kDa] | Score Sequest HT: Sequest HT | Abundance: F2: Sample |
|---|---|---|---|---|---|---|---|---|---|---|
| 0 | O15294 | UDP-N-acetylglucosamine--peptide N-acetylglucosaminyltransferase 110 kDa subunit OS=Homo sapiens OX=9606 GN=OGT PE=1 SV=3 | 80 | 103 | 306 | 103 | 1046 | 116.9 | 869 | 115363466.2 |
| 0 | P35579 | Myosin-9 OS=Homo sapiens OX=9606 GN=MYH9 PE=1 SV=4 | 62 | 134 | 193 | 122 | 1960 | 226.4 | 622.51 | 14201016.81 |
| 0 | P14174 | Macrophage migration inhibitory factor OS=Homo sapiens OX=9606 GN=MIF PE=1 SV=4 | 52 | 8 | 130 | 8 | 115 | 12.5 | 321.37 | 32729618.51 |
| 0 | P0DMV9 | Heat shock 70 kDa protein 1B OS=Homo sapiens OX=9606 GN=HSPA1B PE=1 SV=1 | 69 | 53 | 96 | 40 | 641 | 70 | 282.44 | 16170467.32 |
| 0 | P51610 | Host cell factor 1 OS=Homo sapiens OX=9606 GN=HCFC1 PE=1 SV=2 | 38 | 62 | 97 | 62 | 2035 | 208.6 | 274.98 | 8888667.695 |
| 0 | P08670 | Vimentin OS=Homo sapiens OX=9606 GN=VIM PE=1 SV=4 | 84 | 59 | 97 | 57 | 466 | 53.6 | 269.82 | 16060614.29 |
| 0 | Q13813 | Spectrin alpha chain, non-erythrocytic 1 OS=Homo sapiens OX=9606 GN=SPTAN1 PE=1 SV=3 | 34 | 66 | 81 | 66 | 2472 | 284.4 | 256.87 | 2814855.394 |
| 0 | P35580 | Myosin-10 OS=Homo sapiens OX=9606 GN=MYH10 PE=1 SV=3 | 39 | 66 | 87 | 54 | 1976 | 228.9 | 256.6 | 2625020.935 |
| 0 | P16403 | Histone H1.2 OS=Homo sapiens OX=9606 GN=H1-2 PE=1 SV=2 | 67 | 31 | 80 | 11 | 213 | 21.4 | 226.44 | 33909238.26 |
| 0 | P11142 | Heat shock cognate 71 kDa protein OS=Homo sapiens OX=9606 GN=HSPA8 PE=1 SV=1 | 56 | 45 | 71 | 36 | 646 | 70.9 | 220.84 | 10178115.54 |
| 0 | P63261 | Actin, cytoplasmic 2 OS=Homo sapiens OX=9606 GN=ACTG1 PE=1 SV=1 | 75 | 32 | 61 | 2 | 375 | 41.8 | 219.66 | 332312.2871 |
| 0 | P04264 | Keratin, type II cytoskeletal 1 OS=Homo sapiens OX=9606 GN=KRT1 PE=1 SV=6 | 62 | 47 | 77 | 42 | 644 | 66 | 218.93 | 10486910.1 |
| 0 | P60709 | Actin, cytoplasmic 1 OS=Homo sapiens OX=9606 GN=ACTB PE=1 SV=1 | 75 | 32 | 59 | 2 | 375 | 41.7 | 209.13 | 15760657.6 |
| 0 | P35527 | Keratin, type I cytoskeletal 9 OS=Homo sapiens OX=9606 GN=KRT9 PE=1 SV=3 | 61 | 27 | 50 | 27 | 623 | 62 | 184.65 | 5819552.76 |
| 0 | P10412 | Histone H1.4 OS=Homo sapiens OX=9606 GN=H1-4 PE=1 SV=2 | 56 | 26 | 61 | 5 | 219 | 21.9 | 169.07 | 6328702.288 |
| 0 | P16402 | Histone H1.3 OS=Homo sapiens OX=9606 GN=H1-3 PE=1 SV=2 | 57 | 26 | 59 | 5 | 221 | 22.3 | 166.35 | 530288.2627 |
| 0 | P19338 | Nucleolin OS=Homo sapiens OX=9606 GN=NCL PE=1 SV=3 | 47 | 36 | 51 | 36 | 710 | 76.6 | 158.63 | 3111914.635 |
| 0 | P13645 | Keratin, type I cytoskeletal 10 OS=Homo sapiens OX=9606 GN=KRT10 PE=1 SV=6 | 53 | 26 | 49 | 23 | 584 | 58.8 | 152.76 | 5372389.764 |
| 0 | P07910 | Heterogeneous nuclear ribonucleoproteins C1/C2 OS=Homo sapiens OX=9606 GN=HNRNPC PE=1 SV=4 | 58 | 27 | 54 | 27 | 306 | 33.7 | 152.19 | 14716524.76 |
| 0 | P12956 | X-ray repair cross-complementing protein 6 OS=Homo sapiens OX=9606 GN=XRCC6 PE=1 SV=2 | 52 | 32 | 54 | 32 | 609 | 69.8 | 145.14 | 3934828.592 |
| 0 | P11940 | Polyadenylate-binding protein 1 OS=Homo sapiens OX=9606 GN=PABPC1 PE=1 SV=2 | 59 | 38 | 53 | 13 | 636 | 70.6 | 142.85 | 3869164.55 |
| 0 | O60814 | Histone H2B type 1-K OS=Homo sapiens OX=9606 GN=H2BC12 PE=1 SV=3 | 84 | 24 | 47 | 2 | 126 | 13.9 | 128.59 | 22780317.07 |
| 0 | P35908 | Keratin, type II cytoskeletal 2 epidermal OS=Homo sapiens OX=9606 GN=KRT2 PE=1 SV=2 | 75 | 37 | 40 | 29 | 639 | 65.4 | 127.61 | 3106194.48 |
| 0 | Q5QNW6 | Histone H2B type 2-F OS=Homo sapiens OX=9606 GN=H2BC18 PE=1 SV=3 | 84 | 23 | 45 | 1 | 126 | 13.9 | 127.35 | 229500.7813 |
| 0 | P58876 | Histone H2B type 1-D OS=Homo sapiens OX=9606 GN=H2BC5 PE=1 SV=2 | 84 | 23 | 45 | 1 | 126 | 13.9 | 126.19 | 214872.8594 |
| 0 | Q16778 | Histone H2B type 2-E OS=Homo sapiens OX=9606 GN=H2BC21 PE=1 SV=3 | 84 | 23 | 45 | 0 | 126 | 13.9 | 122.23 | |
| 0 | P33778 | Histone H2B type 1-B OS=Homo sapiens OX=9606 GN=H2BC3 PE=1 SV=2 | 84 | 23 | 44 | 1 | 126 | 13.9 | 119.27 | 1399920.914 |
| 0 | P07197 | Neurofilament medium polypeptide OS=Homo sapiens OX=9606 GN=NEFM PE=1 SV=3 | 38 | 27 | 37 | 25 | 916 | 102.4 | 115.76 | 1593298.258 |
| 0 | P17028 | Zinc finger protein 24 OS=Homo sapiens OX=9606 GN=ZNF24 PE=1 SV=4 | 68 | 21 | 42 | 21 | 368 | 42.1 | 113.66 | 5527581.227 |
| 0 | P68133 | Actin, alpha skeletal muscle OS=Homo sapiens OX=9606 GN=ACTA1 PE=1 SV=1 | 36 | 19 | 37 | 3 | 377 | 42 | 110.5 | 274180.6816 |
| 0 | P11021 | Endoplasmic reticulum chaperone BiP OS=Homo sapiens OX=9606 GN=HSPA5 PE=1 SV=2 | 47 | 26 | 34 | 24 | 654 | 72.3 | 106.88 | 1740696.206 |
| 0 | P46109 | Crk-like protein OS=Homo sapiens OX=9606 GN=CRKL PE=1 SV=1 | 84 | 23 | 34 | 23 | 303 | 33.8 | 103.36 | 3546011.324 |
| 0 | Q9BY77 | Polymerase delta-interacting protein 3 OS=Homo sapiens OX=9606 GN=POLDIP3 PE=1 SV=2 | 66 | 24 | 36 | 24 | 421 | 46.1 | 99.19 | 3181139.182 |
| 0 | P0DJI8 | Serum amyloid A-1 protein OS=Homo sapiens OX=9606 GN=SAA1 PE=1 SV=1 | 75 | 10 | 33 | 6 | 122 | 13.5 | 99.17 | 12412151.31 |
| 0 | P62269 | 40S ribosomal protein S18 OS=Homo sapiens OX=9606 GN=RPS18 PE=1 SV=3 | 76 | 24 | 38 | 24 | 152 | 17.7 | 98.37 | 12018971.41 |
| 0 | P62805 | Histone H4 OS=Homo sapiens OX=9606 GN=H4C1 PE=1 SV=2 | 66 | 21 | 40 | 21 | 103 | 11.4 | 97.17 | 14332033.99 |
| 0 | Q9Y3Y2 | Chromatin target of PRMT1 protein OS=Homo sapiens OX=9606 GN=CHTOP PE=1 SV=2 | 46 | 16 | 28 | 16 | 248 | 26.4 | 95.78 | 11860650.43 |
| 0 | Q12906 | Interleukin enhancer-binding factor 3 OS=Homo sapiens OX=9606 GN=ILF3 PE=1 SV=3 | 34 | 22 | 31 | 22 | 894 | 95.3 | 92.82 | 2076425.77 |
| 0 | Q13310 | Polyadenylate-binding protein 4 OS=Homo sapiens OX=9606 GN=PABPC4 PE=1 SV=1 | 37 | 23 | 34 | 9 | 644 | 70.7 | 92.75 | 464703.751 |
| 0 | P10809 | 60 kDa heat shock protein, mitochondrial OS=Homo sapiens OX=9606 GN=HSPD1 PE=1 SV=2 | 47 | 24 | 30 | 24 | 573 | 61 | 90.15 | 1381821.111 |
| 0 | Q8N257 | Histone H2B type 3-B OS=Homo sapiens OX=9606 GN=H2BU1 PE=1 SV=3 | 59 | 17 | 33 | 1 | 126 | 13.9 | 87.83 | 35872.5625 |
| 0 | P38919 | Eukaryotic initiation factor 4A-III OS=Homo sapiens OX=9606 GN=EIF4A3 PE=1 SV=4 | 51 | 20 | 29 | 20 | 411 | 46.8 | 85.13 | 2997744.945 |
| 0 | Q03111 | Protein ENL OS=Homo sapiens OX=9606 GN=MLLT1 PE=1 SV=2 | 18 | 6 | 24 | 6 | 559 | 62 | 83.81 | 445162.3096 |
| 0 | P60660 | Myosin light polypeptide 6 OS=Homo sapiens OX=9606 GN=MYL6 PE=1 SV=2 | 68 | 17 | 25 | 17 | 151 | 16.9 | 83.18 | 4714127.83 |
| 0 | P67809 | Y-box-binding protein 1 OS=Homo sapiens OX=9606 GN=YBX1 PE=1 SV=3 | 56 | 16 | 31 | 12 | 324 | 35.9 | 81.88 | 2042860.936 |
| 0 | Q86V81 | THO complex subunit 4 OS=Homo sapiens OX=9606 GN=ALYREF PE=1 SV=3 | 66 | 16 | 26 | 16 | 257 | 26.9 | 80.1 | 7189321.639 |
| 0 | Q99878 | Histone H2A type 1-J OS=Homo sapiens OX=9606 GN=H2AC14 PE=1 SV=3 | 52 | 11 | 41 | 3 | 128 | 13.9 | 77.61 | 12752785.17 |
| 0 | Q96FV9 | THO complex subunit 1 OS=Homo sapiens OX=9606 GN=THOC1 PE=1 SV=1 | 44 | 21 | 26 | 21 | 657 | 75.6 | 75.18 | 1106642.917 |
| 0 | P07437 | Tubulin beta chain OS=Homo sapiens OX=9606 GN=TUBB PE=1 SV=2 | 59 | 19 | 22 | 5 | 444 | 49.6 | 74.47 | 257766.4316 |
| 0 | P68371 | Tubulin beta-4B chain OS=Homo sapiens OX=9606 GN=TUBB4B PE=1 SV=1 | 55 | 19 | 23 | 5 | 445 | 49.8 | 72.66 | 868323.4395 |
| 0 | Q12905 | Interleukin enhancer-binding factor 2 OS=Homo sapiens OX=9606 GN=ILF2 PE=1 SV=2 | 51 | 14 | 24 | 14 | 390 | 43 | 72.05 | 860305.4951 |
| 0 | P38159 | RNA-binding motif protein, X chromosome OS=Homo sapiens OX=9606 GN=RBMX PE=1 SV=3 | 51 | 17 | 27 | 17 | 391 | 42.3 | 70.22 | 805134.6484 |
| 0 | P06748 | Nucleophosmin OS=Homo sapiens OX=9606 GN=NPM1 PE=1 SV=2 | 63 | 14 | 21 | 14 | 294 | 32.6 | 69.76 | 1024356.447 |
| 0 | P02768 | Albumin OS=Homo sapiens OX=9606 GN=ALB PE=1 SV=2 | 29 | 18 | 23 | 18 | 609 | 69.3 | 65.5 | 2533714.838 |
| 0 | Q9Y3I0 | RNA-splicing ligase RtcB homolog OS=Homo sapiens OX=9606 GN=RTCB PE=1 SV=1 | 49 | 17 | 24 | 17 | 505 | 55.2 | 65.1 | 948850.1064 |
| 0 | P22626 | Heterogeneous nuclear ribonucleoproteins A2/B1 OS=Homo sapiens OX=9606 GN=HNRNPA2B1 PE=1 SV=2 | 56 | 17 | 25 | 17 | 353 | 37.4 | 64.52 | 1647273.769 |
| 0 | P26373 | 60S ribosomal protein L13 OS=Homo sapiens OX=9606 GN=RPL13 PE=1 SV=4 | 57 | 15 | 21 | 15 | 211 | 24.2 | 64.46 | 1631840.779 |
| 0 | P16989 | Y-box-binding protein 3 OS=Homo sapiens OX=9606 GN=YBX3 PE=1 SV=4 | 51 | 12 | 19 | 8 | 372 | 40.1 | 64.37 | 334523.4863 |
| 0 | P61247 | 40S ribosomal protein S3a OS=Homo sapiens OX=9606 GN=RPS3A PE=1 SV=2 | 72 | 24 | 28 | 24 | 264 | 29.9 | 64.22 | 1900321.723 |
| 0 | Q01082 | Spectrin beta chain, non-erythrocytic 1 OS=Homo sapiens OX=9606 GN=SPTBN1 PE=1 SV=2 | 12 | 19 | 21 | 18 | 2364 | 274.4 | 63.56 | 287734.6533 |

| | | | | | | | | | | |
|---|---|---|---|---|---|---|---|---|---|---|
| 0 | P13010 | X-ray repair cross-complementing protein 5 OS=Homo sapiens OX=9606 GN=XRCC5 PE=1 SV=3 | 27 | 14 | 20 | 14 | 732 | 82.7 | 62.46 | 785378.0947 |
| 0 | Q8NEU8 | DCC-interacting protein 13-beta OS=Homo sapiens OX=9606 GN=APPL2 PE=1 SV=3 | 34 | 14 | 16 | 14 | 664 | 74.4 | 61.74 | 443119.2461 |
| 0 | Q9H361 | Polyadenylate-binding protein 3 OS=Homo sapiens OX=9606 GN=PABPC3 PE=1 SV=2 | 28 | 16 | 23 | 1 | 631 | 70 | 61.15 | 47612.55469 |
| 0 | P62424 | 60S ribosomal protein L7a OS=Homo sapiens OX=9606 GN=RPL7A PE=1 SV=2 | 55 | 18 | 25 | 18 | 266 | 30 | 61.14 | 1509126.969 |
| 0 | P62750 | 60S ribosomal protein L23a OS=Homo sapiens OX=9606 GN=RPL23A PE=1 SV=1 | 56 | 16 | 23 | 16 | 156 | 17.7 | 60.92 | 4055540.229 |
| 0 | Q8NI27 | THO complex subunit 2 OS=Homo sapiens OX=9606 GN=THOC2 PE=1 SV=2 | 16 | 18 | 21 | 18 | 1593 | 182.7 | 60.37 | 733380.8491 |
| 0 | Q9BRP1 | Programmed cell death protein 2-like OS=Homo sapiens OX=9606 GN=PDCD2L PE=1 SV=1 | 43 | 12 | 22 | 12 | 358 | 39.4 | 59.99 | 1698031.396 |
| 0 | P68871 | Hemoglobin subunit beta OS=Homo sapiens OX=9606 GN=HBB PE=1 SV=2 | 76 | 12 | 21 | 6 | 147 | 16 | 59.64 | 5541428.291 |
| 0 | P09651 | Heterogeneous nuclear ribonucleoprotein A1 OS=Homo sapiens OX=9606 GN=HNRNPA1 PE=1 SV=5 | 44 | 15 | 23 | 15 | 372 | 38.7 | 58.8 | 1762427.615 |
| 0 | Q13838 | Spliceosome RNA helicase DDX39B OS=Homo sapiens OX=9606 GN=DDX39B PE=1 SV=1 | 45 | 14 | 20 | 14 | 428 | 49 | 57.99 | 2831581.44 |
| 0 | Q9Y5S9 | RNA-binding protein 8A OS=Homo sapiens OX=9606 GN=RBM8A PE=1 SV=1 | 40 | 9 | 17 | 9 | 174 | 19.9 | 57.45 | 1395541.804 |
| 0 | Q15436 | Protein transport protein Sec23A OS=Homo sapiens OX=9606 GN=SEC23A PE=1 SV=2 | 31 | 15 | 19 | 15 | 765 | 86.1 | 57.2 | 592388.4453 |
| 0 | Q93077 | Histone H2A type 1-C OS=Homo sapiens OX=9606 GN=H2AC6 PE=1 SV=3 | 51 | 9 | 24 | 1 | 130 | 14.1 | 57.09 | 211948.8867 |
| 0 | P62081 | 40S ribosomal protein S7 OS=Homo sapiens OX=9606 GN=RPS7 PE=1 SV=1 | 56 | 11 | 16 | 11 | 194 | 22.1 | 56.41 | 944112.7109 |
| 0 | P84090 | Enhancer of rudimentary homolog OS=Homo sapiens OX=9606 GN=ERH PE=1 SV=1 | 63 | 12 | 26 | 12 | 104 | 12.3 | 55.78 | 19260097.16 |
| 0 | Q06830 | Peroxiredoxin-1 OS=Homo sapiens OX=9606 GN=PRDX1 PE=1 SV=1 | 73 | 12 | 20 | 10 | 199 | 22.1 | 55.07 | 2049873.301 |
| 0 | Q9UKM9 | RNA-binding protein Raly OS=Homo sapiens OX=9606 GN=RALY PE=1 SV=1 | 58 | 16 | 19 | 16 | 306 | 32.4 | 54.98 | 839734.1855 |
| 0 | P62829 | 60S ribosomal protein L23 OS=Homo sapiens OX=9606 GN=RPL23 PE=1 SV=1 | 49 | 11 | 21 | 11 | 140 | 14.9 | 54.36 | 943688.2656 |
| 0 | Q9NZI8 | Insulin-like growth factor 2 mRNA-binding protein 1 OS=Homo sapiens OX=9606 GN=IGF2BP1 PE=1 SV=2 | 35 | 18 | 19 | 17 | 577 | 63.4 | 51.87 | 833259.1357 |
| 0 | P23396 | 40S ribosomal protein S3 OS=Homo sapiens OX=9606 GN=RPS3 PE=1 SV=2 | 53 | 13 | 18 | 13 | 243 | 26.7 | 51.25 | 920446.2832 |
| 0 | P17066 | Heat shock 70 kDa protein 6 OS=Homo sapiens OX=9606 GN=HSPA6 PE=1 SV=2 | 17 | 13 | 18 | 1 | 643 | 71 | 51.16 | 532630.375 |
| 0 | P08708 | 40S ribosomal protein S17 OS=Homo sapiens OX=9606 GN=RPS17 PE=1 SV=2 | 70 | 11 | 17 | 11 | 135 | 15.5 | 50.64 | 1892901.105 |
| 0 | Q9Y2W1 | Thyroid hormone receptor-associated protein 3 OS=Homo sapiens OX=9606 GN=THRAP3 PE=1 SV=2 | 19 | 12 | 16 | 12 | 955 | 108.6 | 49.44 | 724072.8828 |
| 0 | P69905 | Hemoglobin subunit alpha OS=Homo sapiens OX=9606 GN=HBA1 PE=1 SV=2 | 68 | 8 | 16 | 8 | 142 | 15.2 | 48.77 | 4937558.379 |
| 0 | Q08211 | ATP-dependent RNA helicase A OS=Homo sapiens OX=9606 GN=DHX9 PE=1 SV=4 | 19 | 15 | 17 | 15 | 1270 | 140.9 | 46.88 | 402087.4873 |
| 0 | Q9UJV9 | Probable ATP-dependent RNA helicase DDX41 OS=Homo sapiens OX=9606 GN=DDX41 PE=1 SV=2 | 36 | 16 | 16 | 16 | 622 | 69.8 | 46.59 | 687266.8711 |
| 0 | Q86U42 | Polyadenylate-binding protein 2 OS=Homo sapiens OX=9606 GN=PABPN1 PE=1 SV=3 | 50 | 11 | 16 | 11 | 306 | 32.7 | 46.53 | 2945409.15 |
| 0 | Q13769 | THO complex subunit 5 homolog OS=Homo sapiens OX=9606 GN=THOC5 PE=1 SV=2 | 23 | 15 | 18 | 15 | 683 | 78.5 | 45.56 | 600378.5508 |
| 0 | O75533 | Splicing factor 3B subunit 1 OS=Homo sapiens OX=9606 GN=SF3B1 PE=1 SV=3 | 20 | 14 | 18 | 14 | 1304 | 145.7 | 45.05 | 260427.6094 |
| 0 | Q8NC51 | Plasminogen activator inhibitor 1 RNA-binding protein OS=Homo sapiens OX=9606 GN=SERBP1 PE=1 SV=2 | 34 | 13 | 15 | 13 | 408 | 44.9 | 44.51 | 1297109.035 |
| 0 | P61978 | Heterogeneous nuclear ribonucleoprotein K OS=Homo sapiens OX=9606 GN=HNRNPK PE=1 SV=1 | 43 | 11 | 15 | 11 | 463 | 50.9 | 44.36 | 292591.4551 |
| 0 | P68363 | Tubulin alpha-1B chain OS=Homo sapiens OX=9606 GN=TUBA1B PE=1 SV=1 | 43 | 13 | 16 | 13 | 451 | 50.1 | 44 | 702253.4707 |
| 0 | P52272 | Heterogeneous nuclear ribonucleoprotein M OS=Homo sapiens OX=9606 GN=HNRNPM PE=1 SV=3 | 28 | 13 | 15 | 13 | 730 | 77.5 | 42.1 | 394633.1953 |
| 0 | P14618 | Pyruvate kinase PKM OS=Homo sapiens OX=9606 GN=PKM PE=1 SV=4 | 40 | 11 | 12 | 11 | 531 | 57.9 | 41.8 | 540628.0254 |
| 0 | P07196 | Neurofilament light polypeptide OS=Homo sapiens OX=9606 GN=NEFL PE=1 SV=3 | 36 | 15 | 18 | 14 | 543 | 61.5 | 41.19 | 317776.6328 |
| 0 | P23528 | Cofilin-1 OS=Homo sapiens OX=9606 GN=CFL1 PE=1 SV=3 | 86 | 10 | 13 | 10 | 166 | 18.5 | 41.09 | 507319.2168 |
| 0 | O14950 | Myosin regulatory light chain 12B OS=Homo sapiens OX=9606 GN=MYL12B PE=1 SV=2 | 66 | 11 | 13 | 11 | 172 | 19.8 | 41 | 1245289.888 |
| 0 | P06576 | ATP synthase subunit beta, mitochondrial OS=Homo sapiens OX=9606 GN=ATP5F1B PE=1 SV=3 | 32 | 10 | 13 | 10 | 529 | 56.5 | 40.38 | 436472.5303 |
| 0 | Q15084 | Protein disulfide-isomerase A6 OS=Homo sapiens OX=9606 GN=PDIA6 PE=1 SV=1 | 30 | 10 | 12 | 10 | 440 | 48.1 | 39.75 | 449266.5215 |
| 0 | P68400 | Casein kinase II subunit alpha OS=Homo sapiens OX=9606 GN=CSNK2A1 PE=1 SV=1 | 34 | 9 | 12 | 9 | 391 | 45.1 | 39.72 | 355062.9629 |
| 0 | Q13263 | Transcription intermediary factor 1-beta OS=Homo sapiens OX=9606 GN=TRIM28 PE=1 SV=5 | 22 | 11 | 13 | 11 | 835 | 88.5 | 39.7 | 495218.3398 |
| 0 | P62847 | 40S ribosomal protein S24 OS=Homo sapiens OX=9606 GN=RPS24 PE=1 SV=1 | 63 | 11 | 14 | 11 | 133 | 15.4 | 39.59 | 649166.2139 |
| 0 | Q9NYF8 | Bcl-2-associated transcription factor 1 OS=Homo sapiens OX=9606 GN=BCLAF1 PE=1 SV=2 | 18 | 12 | 14 | 12 | 920 | 106.1 | 39.08 | 436049.0518 |
| 0 | P39019 | 40S ribosomal protein S19 OS=Homo sapiens OX=9606 GN=RPS19 PE=1 SV=2 | 50 | 11 | 14 | 11 | 145 | 16.1 | 38.99 | 799116.1504 |
| 0 | P09493 | Tropomyosin alpha-1 chain OS=Homo sapiens OX=9606 GN=TPM1 PE=1 SV=2 | 37 | 9 | 13 | 5 | 284 | 32.7 | 38.24 | 615810.0752 |
| 0 | P62906 | 60S ribosomal protein L10a OS=Homo sapiens OX=9606 GN=RPL10A PE=1 SV=2 | 45 | 10 | 15 | 10 | 217 | 24.8 | 38.2 | 830078.9395 |
| 0 | Q96QD9 | UAP56-interacting factor OS=Homo sapiens OX=9606 GN=FYTTD1 PE=1 SV=3 | 47 | 13 | 16 | 13 | 318 | 35.8 | 38.18 | 578458.9707 |
| 0 | P46779 | 60S ribosomal protein L28 OS=Homo sapiens OX=9606 GN=RPL28 PE=1 SV=3 | 60 | 12 | 15 | 12 | 137 | 15.7 | 38.07 | 1719827.297 |
| 0 | P08865 | 40S ribosomal protein SA OS=Homo sapiens OX=9606 GN=RPSA PE=1 SV=4 | 56 | 11 | 13 | 11 | 295 | 32.8 | 37.68 | 892038.1797 |
| 0 | P0DJI9 | Serum amyloid A-2 protein OS=Homo sapiens OX=9606 GN=SAA2 PE=1 SV=1 | 42 | 5 | 12 | 1 | 122 | 13.5 | 37.29 | |
| 0 | P0DP25 | Calmodulin-3 OS=Homo sapiens OX=9606 GN=CALM3 PE=1 SV=1 | 42 | 6 | 11 | 6 | 149 | 16.8 | 36.73 | 957099.8789 |
| 0 | P61326 | Protein mago nashi homolog OS=Homo sapiens OX=9606 GN=MAGOH PE=1 SV=1 | 47 | 6 | 12 | 1 | 146 | 17.2 | 36.26 | 1237682.18 |
| 0 | P22234 | Multifunctional protein ADE2 OS=Homo sapiens OX=9606 GN=PAICS PE=1 SV=3 | 34 | 10 | 11 | 10 | 425 | 47 | 36.19 | 581907.3428 |
| 0 | O00422 | Histone deacetylase complex subunit SAP18 OS=Homo sapiens OX=9606 GN=SAP18 PE=1 SV=1 | 59 | 11 | 14 | 11 | 153 | 17.6 | 35.95 | 1856665.896 |
| 0 | P62899 | 60S ribosomal protein L31 OS=Homo sapiens OX=9606 GN=RPL31 PE=1 SV=1 | 54 | 10 | 12 | 10 | 125 | 14.5 | 35.92 | 1716133.801 |
| 0 | P68431 | Histone H3.1 OS=Homo sapiens OX=9606 GN=H3C1 PE=1 SV=2 | 59 | 12 | 18 | 1 | 136 | 15.4 | 35.86 | 7473377.496 |
| 0 | Q15459 | Splicing factor 3A subunit 1 OS=Homo sapiens OX=9606 GN=SF3A1 PE=1 SV=1 | 11 | 7 | 11 | 7 | 793 | 88.8 | 35.71 | 177284.2363 |
| 0 | P62263 | 40S ribosomal protein S14 OS=Homo sapiens OX=9606 GN=RPS14 PE=1 SV=3 | 43 | 9 | 14 | 9 | 151 | 16.3 | 35.63 | 1063529.4 |
| 0 | Q02878 | 60S ribosomal protein L6 OS=Homo sapiens OX=9606 GN=RPL6 PE=1 SV=3 | 43 | 13 | 16 | 13 | 288 | 32.7 | 35.22 | 723558.9746 |
| 0 | P68104 | Elongation factor 1-alpha 1 OS=Homo sapiens OX=9606 GN=EEF1A1 PE=1 SV=1 | 32 | 7 | 12 | 7 | 462 | 50.1 | 35.2 | 472275.8896 |
| 0 | Q71DI3 | Histone H3.2 OS=Homo sapiens OX=9606 GN=H3C15 PE=1 SV=3 | 59 | 12 | 18 | 1 | 136 | 15.4 | 35.18 | 6429.286133 |
| 0 | P08779 | Keratin, type I cytoskeletal 16 OS=Homo sapiens OX=9606 GN=KRT16 PE=1 SV=4 | 23 | 11 | 13 | 2 | 473 | 51.2 | 34.89 | |
| 0 | P12268 | Inosine-5'-monophosphate dehydrogenase 2 OS=Homo sapiens OX=9606 GN=IMPDH2 PE=1 SV=2 | 27 | 11 | 12 | 11 | 514 | 55.8 | 34.85 | 486017.5088 |

| | | | | | | | | | | |
|---|---|---|---|---|---|---|---|---|---|---|
| 0 | Q86W42 | THO complex subunit 6 homolog OS=Homo sapiens OX=9606 GN=THOC6 PE=1 SV=1 | 39 | 10 | 13 | 10 | 341 | 37.5 | 34.61 | 415020.6191 |
| 0 | Q13435 | Splicing factor 3B subunit 2 OS=Homo sapiens OX=9606 GN=SF3B2 PE=1 SV=2 | 24 | 11 | 13 | 11 | 895 | 100.2 | 34.59 | 211422.6699 |
| 0 | P02533 | Keratin, type I cytoskeletal 14 OS=Homo sapiens OX=9606 GN=KRT14 PE=1 SV=4 | 21 | 11 | 13 | 2 | 472 | 51.5 | 34.35 | 284962.3555 |
| 0 | P30050 | 60S ribosomal protein L12 OS=Homo sapiens OX=9606 GN=RPL12 PE=1 SV=1 | 64 | 9 | 11 | 9 | 165 | 17.8 | 34.2 | 386313.249 |
| 0 | Q96A72 | Protein mago nashi homolog 2 OS=Homo sapiens OX=9606 GN=MAGOHB PE=1 SV=1 | 47 | 6 | 11 | 1 | 148 | 17.3 | 34.04 | 45861.44922 |
| 0 | P61254 | 60S ribosomal protein L26 OS=Homo sapiens OX=9606 GN=RPL26 PE=1 SV=1 | 58 | 11 | 14 | 3 | 145 | 17.2 | 33.74 | 3802340.008 |
| 0 | P38646 | Stress-70 protein, mitochondrial OS=Homo sapiens OX=9606 GN=HSPA9 PE=1 SV=2 | 24 | 14 | 16 | 14 | 679 | 73.6 | 33.68 | 1120014.814 |
| 0 | Q99497 | Parkinson disease protein 7 OS=Homo sapiens OX=9606 GN=PARK7 PE=1 SV=2 | 66 | 9 | 11 | 9 | 189 | 19.9 | 33.64 | 663927.916 |
| 0 | P62249 | 40S ribosomal protein S16 OS=Homo sapiens OX=9606 GN=RPS16 PE=1 SV=2 | 42 | 7 | 12 | 7 | 146 | 16.4 | 33.52 | 612189.4102 |
| 0 | P84243 | Histone H3.3 OS=Homo sapiens OX=9606 GN=H3-3A PE=1 SV=2 | 59 | 12 | 18 | 1 | 136 | 15.3 | 33.24 | 16310.11523 |
| 0 | O75152 | Zinc finger CCCH domain-containing protein 11A OS=Homo sapiens OX=9606 GN=ZC3H11A PE=1 SV=3 | 20 | 10 | 12 | 10 | 810 | 89.1 | 33.19 | 161872.1914 |
| 0 | P05387 | 60S acidic ribosomal protein P2 OS=Homo sapiens OX=9606 GN=RPLP2 PE=1 SV=1 | 82 | 6 | 8 | 6 | 115 | 11.7 | 33.08 | 318462.582 |
| 0 | P35232 | Prohibitin OS=Homo sapiens OX=9606 GN=PHB PE=1 SV=1 | 39 | 7 | 10 | 7 | 272 | 29.8 | 32.94 | 127498.6914 |
| 0 | P48634 | Protein PRRC2A OS=Homo sapiens OX=9606 GN=PRRC2A PE=1 SV=3 | 12 | 15 | 15 | 15 | 2157 | 228.7 | 32.86 | 532458.4844 |
| 0 | Q9H307 | Pinin OS=Homo sapiens OX=9606 GN=PNN PE=1 SV=5 | 17 | 11 | 14 | 11 | 717 | 81.6 | 32.7 | 349996.084 |
| 0 | O60884 | DnaJ homolog subfamily A member 2 OS=Homo sapiens OX=9606 GN=DNAJA2 PE=1 SV=1 | 40 | 13 | 14 | 13 | 412 | 45.7 | 32.62 | 794193.4141 |
| 0 | Q9NYU2 | UDP-glucose:glycoprotein glucosyltransferase 1 OS=Homo sapiens OX=9606 GN=UGGT1 PE=1 SV=3 | 17 | 13 | 14 | 13 | 1555 | 177.1 | 32.29 | 281784.623 |
| 0 | P05455 | Lupus La protein OS=Homo sapiens OX=9606 GN=SSB PE=1 SV=2 | 28 | 10 | 10 | 10 | 408 | 46.8 | 32.29 | 256979.9338 |
| 0 | P35637 | RNA-binding protein FUS OS=Homo sapiens OX=9606 GN=FUS PE=1 SV=1 | 26 | 8 | 11 | 7 | 526 | 53.4 | 32.29 | 230489.9141 |
| 0 | Q9Y265 | RuvB-like 1 OS=Homo sapiens OX=9606 GN=RUVBL1 PE=1 SV=1 | 32 | 8 | 10 | 8 | 456 | 50.2 | 31.99 | 241253.4063 |
| 0 | Q9Y3C6 | Peptidyl-prolyl cis-trans isomerase-like 1 OS=Homo sapiens OX=9606 GN=PPIL1 PE=1 SV=1 | 43 | 7 | 10 | 7 | 166 | 18.2 | 31.83 | 989821.2598 |
| 0 | P25398 | 40S ribosomal protein S12 OS=Homo sapiens OX=9606 GN=RPS12 PE=1 SV=3 | 74 | 8 | 10 | 8 | 132 | 14.5 | 31.75 | 418705.8613 |
| 0 | P62937 | Peptidyl-prolyl cis-trans isomerase A OS=Homo sapiens OX=9606 GN=PPIA PE=1 SV=2 | 82 | 10 | 11 | 10 | 165 | 18 | 31.57 | 702559.458 |
| 0 | Q96PK6 | RNA-binding protein 14 OS=Homo sapiens OX=9606 GN=RBM14 PE=1 SV=2 | 22 | 12 | 14 | 12 | 669 | 69.4 | 31.55 | 892249.6309 |
| 0 | P67936 | Tropomyosin alpha-4 chain OS=Homo sapiens OX=9606 GN=TPM4 PE=1 SV=3 | 31 | 6 | 10 | 1 | 248 | 28.5 | 31.32 | 11808.50977 |
| 0 | Q9UNX3 | 60S ribosomal protein L26-like 1 OS=Homo sapiens OX=9606 GN=RPL26L1 PE=1 SV=1 | 58 | 10 | 12 | 2 | 145 | 17.2 | 31.14 | 113529.0625 |
| 0 | Q9Y224 | RNA transcription, translation and transport factor protein OS=Homo sapiens OX=9606 GN=RTRAF PE=1 SV=1 | 42 | 7 | 11 | 7 | 244 | 28.1 | 31.06 | 334064.9961 |
| 0 | P47914 | 60S ribosomal protein L29 OS=Homo sapiens OX=9606 GN=RPL29 PE=1 SV=2 | 33 | 8 | 11 | 8 | 159 | 17.7 | 30.81 | 1247000.715 |
| 0 | P11233 | Ras-related protein Ral-A OS=Homo sapiens OX=9606 GN=RALA PE=1 SV=1 | 54 | 10 | 11 | 10 | 206 | 23.6 | 30.55 | 396186.0039 |
| 0 | P52907 | F-actin-capping protein subunit alpha-1 OS=Homo sapiens OX=9606 GN=CAPZA1 PE=1 SV=3 | 51 | 9 | 13 | 7 | 286 | 32.9 | 30.47 | 408742.1768 |
| 0 | P62318 | Small nuclear ribonucleoprotein Sm D3 OS=Homo sapiens OX=9606 GN=SNRPD3 PE=1 SV=1 | 60 | 6 | 9 | 6 | 126 | 13.9 | 30.4 | 393730.125 |
| 0 | Q9Y230 | RuvB-like 2 OS=Homo sapiens OX=9606 GN=RUVBL2 PE=1 SV=3 | 26 | 10 | 12 | 10 | 463 | 51.1 | 30.25 | 588159.4492 |
| 0 | P09661 | U2 small nuclear ribonucleoprotein A' OS=Homo sapiens OX=9606 GN=SNRPA1 PE=1 SV=2 | 52 | 8 | 9 | 8 | 255 | 28.4 | 29.94 | 169616.752 |
| 0 | P62753 | 40S ribosomal protein S6 OS=Homo sapiens OX=9606 GN=RPS6 PE=1 SV=1 | 39 | 7 | 10 | 7 | 249 | 28.7 | 29.86 | 458855.7031 |
| 0 | P14678 | Small nuclear ribonucleoprotein-associated proteins B and B' OS=Homo sapiens OX=9606 GN=SNRPB PE=1 SV=2 | 26 | 6 | 9 | 6 | 240 | 24.6 | 29.49 | 436726.3711 |
| 0 | P06753 | Tropomyosin alpha-3 chain OS=Homo sapiens OX=9606 GN=TPM3 PE=1 SV=2 | 31 | 7 | 10 | 2 | 285 | 32.9 | 29.31 | 192581.7813 |
| 0 | P05787 | Keratin, type II cytoskeletal 8 OS=Homo sapiens OX=9606 GN=KRT8 PE=1 SV=7 | 23 | 11 | 11 | 6 | 483 | 53.7 | 29.11 | 156526.8691 |
| 0 | Q9UHV9 | Prefoldin subunit 2 OS=Homo sapiens OX=9606 GN=PFDN2 PE=1 SV=1 | 75 | 10 | 12 | 10 | 154 | 16.6 | 29.06 | 957748.957 |
| 0 | P84103 | Serine/arginine-rich splicing factor 3 OS=Homo sapiens OX=9606 GN=SRSF3 PE=1 SV=1 | 51 | 9 | 12 | 8 | 164 | 19.3 | 29.05 | 501219.0117 |
| 0 | Q9UKV3 | Apoptotic chromatin condensation inducer in the nucleus OS=Homo sapiens OX=9606 GN=ACIN1 PE=1 SV=2 | 11 | 10 | 11 | 10 | 1341 | 151.8 | 28.89 | 162910.415 |
| 0 | P62826 | GTP-binding nuclear protein Ran OS=Homo sapiens OX=9606 GN=RAN PE=1 SV=3 | 42 | 9 | 12 | 9 | 216 | 24.4 | 28.6 | 1005335.125 |
| 0 | P55209 | Nucleosome assembly protein 1-like 1 OS=Homo sapiens OX=9606 GN=NAP1L1 PE=1 SV=1 | 32 | 7 | 9 | 7 | 391 | 45.3 | 28.5 | 84172.76953 |
| 0 | P07305 | Histone H1.0 OS=Homo sapiens OX=9606 GN=H1-0 PE=1 SV=3 | 31 | 9 | 10 | 9 | 194 | 20.9 | 28.41 | 260924.3394 |
| 0 | Q15029 | 116 kDa U5 small nuclear ribonucleoprotein component OS=Homo sapiens OX=9606 GN=EFTUD2 PE=1 SV=1 | 15 | 9 | 9 | 9 | 972 | 109.4 | 28.06 | 109705.8818 |
| 0 | O15020 | Spectrin beta chain, non-erythrocytic 2 OS=Homo sapiens OX=9606 GN=SPTBN2 PE=1 SV=3 | 4 | 7 | 9 | 6 | 2390 | 271.2 | 27.99 | 71069.75 |
| 0 | P32119 | Peroxiredoxin-2 OS=Homo sapiens OX=9606 GN=PRDX2 PE=1 SV=5 | 56 | 7 | 9 | 6 | 198 | 21.9 | 27.96 | 159670.1533 |
| 0 | P62280 | 40S ribosomal protein S11 OS=Homo sapiens OX=9606 GN=RPS11 PE=1 SV=3 | 59 | 13 | 13 | 13 | 158 | 18.4 | 27.55 | 1147125.773 |
| 0 | Q99729 | Heterogeneous nuclear ribonucleoprotein A/B OS=Homo sapiens OX=9606 GN=HNRNPAB PE=1 SV=2 | 23 | 7 | 10 | 7 | 332 | 36.2 | 27.52 | 220154.3135 |
| 0 | Q92499 | ATP-dependent RNA helicase DDX1 OS=Homo sapiens OX=9606 GN=DDX1 PE=1 SV=2 | 21 | 9 | 13 | 9 | 740 | 82.4 | 27.49 | 244989.1846 |
| 0 | Q15717 | ELAV-like protein 1 OS=Homo sapiens OX=9606 GN=ELAVL1 PE=1 SV=2 | 35 | 7 | 8 | 7 | 326 | 36.1 | 26.59 | 241562.5371 |
| 0 | P40926 | Malate dehydrogenase, mitochondrial OS=Homo sapiens OX=9606 GN=MDH2 PE=1 SV=3 | 32 | 6 | 7 | 6 | 338 | 35.5 | 26.59 | 168147.4609 |
| 0 | Q13573 | SNW domain-containing protein 1 OS=Homo sapiens OX=9606 GN=SNW1 PE=1 SV=1 | 22 | 6 | 8 | 6 | 536 | 61.5 | 26.49 | 111091.5332 |
| 0 | Q12765 | Secernin-1 OS=Homo sapiens OX=9606 GN=SCRN1 PE=1 SV=2 | 20 | 7 | 9 | 7 | 414 | 46.4 | 26.3 | 832047.8535 |
| 0 | P26599 | Polypyrimidine tract-binding protein 1 OS=Homo sapiens OX=9606 GN=PTBP1 PE=1 SV=1 | 19 | 9 | 11 | 9 | 531 | 57.2 | 26.07 | 498172.1289 |
| 0 | P15880 | 40S ribosomal protein S2 OS=Homo sapiens OX=9606 GN=RPS2 PE=1 SV=2 | 31 | 10 | 12 | 10 | 293 | 31.3 | 25.22 | 298325.3887 |
| 0 | P18621 | 60S ribosomal protein L17 OS=Homo sapiens OX=9606 GN=RPL17 PE=1 SV=3 | 59 | 8 | 10 | 8 | 184 | 21.4 | 25.16 | 446017.8965 |
| 0 | P51991 | Heterogeneous nuclear ribonucleoprotein A3 OS=Homo sapiens OX=9606 GN=HNRNPA3 PE=1 SV=2 | 31 | 7 | 9 | 7 | 378 | 39.6 | 25.13 | 218736.1816 |
| 0 | P13647 | Keratin, type II cytoskeletal 5 OS=Homo sapiens OX=9606 GN=KRT5 PE=1 SV=3 | 15 | 10 | 11 | 4 | 590 | 62.3 | 25.08 | 165921.3486 |
| 0 | P62917 | 60S ribosomal protein L8 OS=Homo sapiens OX=9606 GN=RPL8 PE=1 SV=2 | 36 | 9 | 10 | 9 | 257 | 28 | 25.05 | 554058.6699 |
| 0 | P62910 | 60S ribosomal protein L32 OS=Homo sapiens OX=9606 GN=RPL32 PE=1 SV=2 | 43 | 6 | 9 | 6 | 135 | 15.9 | 25.03 | 1037090.167 |
| 0 | P09012 | U1 small nuclear ribonucleoprotein A OS=Homo sapiens OX=9606 GN=SNRPA PE=1 SV=3 | 30 | 6 | 9 | 6 | 282 | 31.3 | 24.89 | 256784.1738 |
| 0 | P62701 | 40S ribosomal protein S4, X isoform OS=Homo sapiens OX=9606 GN=RPS4X PE=1 SV=2 | 41 | 11 | 11 | 11 | 263 | 29.6 | 24.89 | 612189.2598 |
| 0 | Q15287 | RNA-binding protein with serine-rich domain 1 OS=Homo sapiens OX=9606 GN=RNPS1 PE=1 SV=1 | 29 | 7 | 9 | 7 | 305 | 34.2 | 24.68 | 264013.9902 |

| | | | | | | | | | | |
|---|---|---|---|---|---|---|---|---|---|---|
| 0 | P04259 | Keratin, type II cytoskeletal 6B OS=Homo sapiens OX=9606 GN=KRT6B PE=1 SV=5 | 13 | 9 | 10 | 1 | 564 | 60 | 24.62 | 47806.45313 |
| 0 | P29692 | Elongation factor 1-delta OS=Homo sapiens OX=9606 GN=EEF1D PE=1 SV=5 | 34 | 8 | 9 | 6 | 281 | 31.1 | 24.57 | 279458.0166 |
| 0 | P02794 | Ferritin heavy chain OS=Homo sapiens OX=9606 GN=FTH1 PE=1 SV=2 | 43 | 6 | 7 | 6 | 183 | 21.2 | 24.36 | 190550.8203 |
| 0 | P42766 | 60S ribosomal protein L35 OS=Homo sapiens OX=9606 GN=RPL35 PE=1 SV=2 | 44 | 7 | 12 | 7 | 123 | 14.5 | 24.3 | 1227941.559 |
| 0 | P62888 | 60S ribosomal protein L30 OS=Homo sapiens OX=9606 GN=RPL30 PE=1 SV=2 | 72 | 7 | 8 | 7 | 115 | 12.8 | 23.93 | 829166.1465 |
| 0 | P83731 | 60S ribosomal protein L24 OS=Homo sapiens OX=9606 GN=RPL24 PE=1 SV=1 | 40 | 8 | 10 | 8 | 157 | 17.8 | 23.75 | 462950.4355 |
| 0 | P02042 | Hemoglobin subunit delta OS=Homo sapiens OX=9606 GN=HBD PE=1 SV=2 | 52 | 7 | 10 | 1 | 147 | 16 | 23.56 | 19177.34766 |
| 0 | Q8N163 | Cell cycle and apoptosis regulator protein 2 OS=Homo sapiens OX=9606 GN=CCAR2 PE=1 SV=2 | 15 | 9 | 9 | 9 | 923 | 102.8 | 23.54 | 257666.4961 |
| 0 | P49327 | Fatty acid synthase OS=Homo sapiens OX=9606 GN=FASN PE=1 SV=3 | 6 | 7 | 8 | 7 | 2511 | 273.3 | 23.43 | 84977.37305 |
| 0 | P62273 | 40S ribosomal protein S29 OS=Homo sapiens OX=9606 GN=RPS29 PE=1 SV=2 | 59 | 5 | 11 | 5 | 56 | 6.7 | 23.26 | 3042404.654 |
| 0 | P37108 | Signal recognition particle 14 kDa protein OS=Homo sapiens OX=9606 GN=SRP14 PE=1 SV=2 | 46 | 7 | 9 | 7 | 136 | 14.6 | 23.11 | 580024.9102 |
| 0 | P06733 | Alpha-enolase OS=Homo sapiens OX=9606 GN=ENO1 PE=1 SV=2 | 28 | 7 | 7 | 7 | 434 | 47.1 | 22.9 | 112449.9805 |
| 0 | Q92522 | Histone H1.10 OS=Homo sapiens OX=9606 GN=H1-10 PE=1 SV=1 | 46 | 10 | 10 | 10 | 213 | 22.5 | 22.82 | 990452.3262 |
| 0 | Q9Y3U8 | 60S ribosomal protein L36 OS=Homo sapiens OX=9606 GN=RPL36 PE=1 SV=3 | 50 | 8 | 11 | 8 | 105 | 12.2 | 22.78 | 625490.957 |
| 0 | P62277 | 40S ribosomal protein S13 OS=Homo sapiens OX=9606 GN=RPS13 PE=1 SV=2 | 47 | 9 | 10 | 9 | 151 | 17.2 | 22.75 | 596523.418 |
| 0 | P60866 | 40S ribosomal protein S20 OS=Homo sapiens OX=9606 GN=RPS20 PE=1 SV=1 | 34 | 4 | 8 | 4 | 119 | 13.4 | 22.74 | 1191812.906 |
| 0 | Q9UHX1 | Poly(U)-binding-splicing factor PUF60 OS=Homo sapiens OX=9606 GN=PUF60 PE=1 SV=1 | 23 | 7 | 9 | 7 | 559 | 59.8 | 22.68 | 229019.5 |
| 0 | Q00839 | Heterogeneous nuclear ribonucleoprotein U OS=Homo sapiens OX=9606 GN=HNRNPU PE=1 SV=6 | 14 | 7 | 8 | 7 | 825 | 90.5 | 22.59 | 137468.0762 |
| 0 | P62316 | Small nuclear ribonucleoprotein Sm D2 OS=Homo sapiens OX=9606 GN=SNRPD2 PE=1 SV=1 | 75 | 8 | 8 | 8 | 118 | 13.5 | 22.47 | 873398.4219 |
| 0 | P62841 | 40S ribosomal protein S15 OS=Homo sapiens OX=9606 GN=RPS15 PE=1 SV=2 | 39 | 6 | 11 | 6 | 145 | 17 | 22.41 | 787165.9277 |
| 0 | Q9ULU4 | Protein kinase C-binding protein 1 OS=Homo sapiens OX=9606 GN=ZMYND8 PE=1 SV=2 | 12 | 7 | 7 | 7 | 1186 | 131.6 | 22.29 | 89727.22852 |
| 0 | P05141 | ADP/ATP translocase 2 OS=Homo sapiens OX=9606 GN=SLC25A5 PE=1 SV=7 | 17 | 6 | 7 | 6 | 298 | 32.8 | 22.25 | 312974.4004 |
| 0 | P63104 | 14-3-3 protein zeta/delta OS=Homo sapiens OX=9606 GN=YWHAZ PE=1 SV=1 | 34 | 7 | 7 | 6 | 245 | 27.7 | 22.25 | 233250.2891 |
| 0 | P53999 | Activated RNA polymerase II transcriptional coactivator p15 OS=Homo sapiens OX=9606 GN=SUB1 PE=1 SV=3 | 43 | 6 | 8 | 6 | 127 | 14.4 | 21.6 | 207912.1758 |
| 0 | C9JLW8 | Mapk-regulated corepressor-interacting protein 1 OS=Homo sapiens OX=9606 GN=MCRIP1 PE=1 SV=1 | 70 | 6 | 7 | 6 | 97 | 10.9 | 21.53 | 292284.1953 |
| 0 | P24534 | Elongation factor 1-beta OS=Homo sapiens OX=9606 GN=EEF1B2 PE=1 SV=3 | 31 | 7 | 9 | 5 | 225 | 24.7 | 21.02 | 255797.9502 |
| 0 | Q9H4A5 | Golgi phosphoprotein 3-like OS=Homo sapiens OX=9606 GN=GOLPH3L PE=1 SV=1 | 31 | 4 | 6 | 4 | 285 | 32.7 | 20.73 | 122585.1719 |
| 0 | P46778 | 60S ribosomal protein L21 OS=Homo sapiens OX=9606 GN=RPL21 PE=1 SV=2 | 47 | 7 | 8 | 7 | 160 | 18.6 | 20.7 | 671526.6836 |
| 0 | P61964 | WD repeat-containing protein 5 OS=Homo sapiens OX=9606 GN=WDR5 PE=1 SV=1 | 18 | 4 | 7 | 4 | 334 | 36.6 | 20.31 | 218115.7051 |
| 0 | P61513 | 60S ribosomal protein L37a OS=Homo sapiens OX=9606 GN=RPL37A PE=1 SV=2 | 68 | 7 | 9 | 7 | 92 | 10.3 | 20.31 | 601632.0352 |
| 0 | Q6I9Y2 | THO complex subunit 7 homolog OS=Homo sapiens OX=9606 GN=THOC7 PE=1 SV=3 | 38 | 6 | 6 | 6 | 204 | 23.7 | 19.99 | 674077.0742 |
| 0 | P42677 | 40S ribosomal protein S27 OS=Homo sapiens OX=9606 GN=RPS27 PE=1 SV=3 | 40 | 5 | 7 | 3 | 84 | 9.5 | 19.67 | 498263.3594 |
| 0 | Q14257 | Reticulocalbin-2 OS=Homo sapiens OX=9606 GN=RCN2 PE=1 SV=1 | 15 | 4 | 7 | 4 | 317 | 36.9 | 19.52 | 146992.2773 |
| 0 | Q8IUE6 | Histone H2A type 2-B OS=Homo sapiens OX=9606 GN=H2AC21 PE=1 SV=3 | 45 | 6 | 8 | 3 | 130 | 14 | 19.52 | 102835.4434 |
| 0 | P63173 | 60S ribosomal protein L38 OS=Homo sapiens OX=9606 GN=RPL38 PE=1 SV=2 | 40 | 4 | 7 | 4 | 70 | 8.2 | 19.46 | 838434.2656 |
| 0 | P49411 | Elongation factor Tu, mitochondrial OS=Homo sapiens OX=9606 GN=TUFM PE=1 SV=2 | 16 | 4 | 6 | 4 | 452 | 49.5 | 19.42 | 83664.72461 |
| 0 | Q07955 | Serine/arginine-rich splicing factor 1 OS=Homo sapiens OX=9606 GN=SRSF1 PE=1 SV=2 | 36 | 7 | 8 | 6 | 248 | 27.7 | 19.29 | 599040.3047 |
| 0 | O14979 | Heterogeneous nuclear ribonucleoprotein D-like OS=Homo sapiens OX=9606 GN=HNRNPDL PE=1 SV=3 | 10 | 4 | 6 | 4 | 420 | 46.4 | 19.27 | 90801.49609 |
| 0 | P05204 | Non-histone chromosomal protein HMG-17 OS=Homo sapiens OX=9606 GN=HMGN2 PE=1 SV=3 | 17 | 1 | 9 | 1 | 90 | 9.4 | 19.21 | 504067.3301 |
| 0 | P14866 | Heterogeneous nuclear ribonucleoprotein L OS=Homo sapiens OX=9606 GN=HNRNPL PE=1 SV=2 | 14 | 4 | 5 | 4 | 589 | 64.1 | 19 | 109529.0557 |
| 0 | E9PRG8 | Uncharacterized protein C11orf98 OS=Homo sapiens OX=9606 GN=C11orf98 PE=4 SV=2 | 34 | 4 | 6 | 4 | 123 | 14.2 | 18.86 | 1722987.172 |
| 0 | P62979 | Ubiquitin-40S ribosomal protein S27a OS=Homo sapiens OX=9606 GN=RPS27A PE=1 SV=2 | 44 | 6 | 8 | 2 | 156 | 18 | 18.45 | 263483.2207 |
| 0 | Q13151 | Heterogeneous nuclear ribonucleoprotein A0 OS=Homo sapiens OX=9606 GN=HNRNPA0 PE=1 SV=1 | 25 | 6 | 7 | 6 | 305 | 30.8 | 18.39 | 315406.3789 |
| 0 | P31946 | 14-3-3 protein beta/alpha OS=Homo sapiens OX=9606 GN=YWHAB PE=1 SV=3 | 32 | 5 | 5 | 3 | 246 | 28.1 | 18.37 | 94890.09668 |
| 0 | P55769 | NHP2-like protein 1 OS=Homo sapiens OX=9606 GN=SNU13 PE=1 SV=3 | 45 | 5 | 6 | 5 | 128 | 14.2 | 18.34 | 220761.9102 |
| 0 | P31689 | DnaJ homolog subfamily A member 1 OS=Homo sapiens OX=9606 GN=DNAJA1 PE=1 SV=2 | 20 | 5 | 8 | 5 | 397 | 44.8 | 18.23 | 135074.8306 |
| 0 | O75643 | U5 small nuclear ribonucleoprotein 200 kDa helicase OS=Homo sapiens OX=9606 GN=SNRNP200 PE=1 SV=2 | 4 | 6 | 6 | 6 | 2136 | 244.4 | 17.92 | 74087.73438 |
| 0 | P35268 | 60S ribosomal protein L22 OS=Homo sapiens OX=9606 GN=RPL22 PE=1 SV=2 | 63 | 5 | 5 | 5 | 128 | 14.8 | 17.79 | 287769 |
| 0 | Q16629 | Serine/arginine-rich splicing factor 7 OS=Homo sapiens OX=9606 GN=SRSF7 PE=1 SV=1 | 19 | 4 | 7 | 3 | 238 | 27.4 | 17.76 | 172188.9043 |
| 0 | Q08380 | Galectin-3-binding protein OS=Homo sapiens OX=9606 GN=LGALS3BP PE=1 SV=1 | 10 | 4 | 6 | 4 | 585 | 65.3 | 17.63 | 54229.79541 |
| 0 | P42771 | Cyclin-dependent kinase inhibitor 2A OS=Homo sapiens OX=9606 GN=CDKN2A PE=1 SV=2 | 51 | 5 | 7 | 5 | 156 | 16.5 | 17.21 | 275251.4512 |
| 0 | Q9C005 | Protein dpy-30 homolog OS=Homo sapiens OX=9606 GN=DPY30 PE=1 SV=1 | 57 | 4 | 5 | 4 | 99 | 11.2 | 17.14 | 477847.5 |
| 0 | Q6UXN9 | WD repeat-containing protein 82 OS=Homo sapiens OX=9606 GN=WDR82 PE=1 SV=1 | 28 | 5 | 6 | 5 | 313 | 35.1 | 17.09 | 93694.39941 |
| 0 | P17844 | Probable ATP-dependent RNA helicase DDX5 OS=Homo sapiens OX=9606 GN=DDX5 PE=1 SV=1 | 12 | 7 | 7 | 6 | 614 | 69.1 | 16.91 | 171241.9834 |
| 0 | Q13185 | Chromobox protein homolog 3 OS=Homo sapiens OX=9606 GN=CBX3 PE=1 SV=4 | 36 | 5 | 5 | 5 | 183 | 20.8 | 16.86 | 96261.93018 |
| 0 | P46777 | 60S ribosomal protein L5 OS=Homo sapiens OX=9606 GN=RPL5 PE=1 SV=3 | 27 | 5 | 6 | 5 | 297 | 34.3 | 16.79 | 92657.46387 |
| 0 | Q53GL7 | Protein mono-ADP-ribosyltransferase PARP10 OS=Homo sapiens OX=9606 GN=PARP10 PE=1 SV=2 | 16 | 5 | 7 | 5 | 1025 | 109.9 | 16.68 | 114951.7217 |
| 0 | Q15233 | Non-POU domain-containing octamer-binding protein OS=Homo sapiens OX=9606 GN=NONO PE=1 SV=4 | 22 | 7 | 7 | 7 | 471 | 54.2 | 16.37 | 162346.4873 |
| 0 | P31943 | Heterogeneous nuclear ribonucleoprotein H OS=Homo sapiens OX=9606 GN=HNRNPH1 PE=1 SV=4 | 14 | 6 | 6 | 6 | 449 | 49.2 | 16.36 | 177692.541 |
| 0 | P46776 | 60S ribosomal protein L27a OS=Homo sapiens OX=9606 GN=RPL27A PE=1 SV=2 | 37 | 4 | 5 | 4 | 148 | 16.6 | 16.3 | 245711.3828 |
| 0 | Q6DD87 | Zinc finger protein 787 OS=Homo sapiens OX=9606 GN=ZNF787 PE=1 SV=4 | 23 | 6 | 6 | 6 | 382 | 40.4 | 16.16 | 123187.3574 |
| 0 | P62258 | 14-3-3 protein epsilon OS=Homo sapiens OX=9606 GN=YWHAE PE=1 SV=1 | 32 | 5 | 6 | 5 | 255 | 29.2 | 16.13 | 430638.9238 |
| 0 | Q14974 | Importin subunit beta-1 OS=Homo sapiens OX=9606 GN=KPNB1 PE=1 SV=2 | 5 | 4 | 6 | 4 | 876 | 97.1 | 16.12 | 93448.36328 |

| | | | | | | | | | | |
|---|---|---|---|---|---|---|---|---|---|---|
| 0 | O00571 | ATP-dependent RNA helicase DDX3X OS=Homo sapiens OX=9606 GN=DDX3X PE=1 SV=3 | 11 | 5 | 6 | 5 | 662 | 73.2 | 16.08 | 90094.53906 |
| 0 | O95793 | Double-stranded RNA-binding protein Staufen homolog 1 OS=Homo sapiens OX=9606 GN=STAU1 PE=1 SV=2 | 21 | 7 | 7 | 7 | 577 | 63.1 | 15.94 | 181672.6553 |
| 0 | O43809 | Cleavage and polyadenylation specificity factor subunit 5 OS=Homo sapiens OX=9606 GN=NUDT21 PE=1 SV=1 | 35 | 4 | 5 | 4 | 227 | 26.2 | 15.87 | 277929.2266 |
| 0 | Q16531 | DNA damage-binding protein 1 OS=Homo sapiens OX=9606 GN=DDB1 PE=1 SV=1 | 6 | 4 | 5 | 4 | 1140 | 126.9 | 15.71 | 52509.60352 |
| 0 | P31040 | Succinate dehydrogenase [ubiquinone] flavoprotein subunit, mitochondrial OS=Homo sapiens OX=9606 GN=SDHA PE=1 SV=2 | 14 | 5 | 6 | 5 | 664 | 72.6 | 15.63 | 57343.20996 |
| 0 | P51149 | Ras-related protein Rab-7a OS=Homo sapiens OX=9606 GN=RAB7A PE=1 SV=1 | 28 | 7 | 7 | 7 | 207 | 23.5 | 15.63 | 224610.9473 |
| 0 | Q9Y266 | Nuclear migration protein nudC OS=Homo sapiens OX=9606 GN=NUDC PE=1 SV=1 | 16 | 5 | 6 | 5 | 331 | 38.2 | 15.62 | 177940.6621 |
| 0 | P67870 | Casein kinase II subunit beta OS=Homo sapiens OX=9606 GN=CSNK2B PE=1 SV=1 | 34 | 4 | 5 | 4 | 215 | 24.9 | 15.44 | 273874.3867 |
| 0 | P61604 | 10 kDa heat shock protein, mitochondrial OS=Homo sapiens OX=9606 GN=HSPE1 PE=1 SV=2 | 51 | 5 | 6 | 5 | 102 | 10.9 | 15.44 | 217583.7422 |
| 0 | Q71UI9 | Histone H2A.V OS=Homo sapiens OX=9606 GN=H2AZ2 PE=1 SV=3 | 31 | 4 | 10 | 2 | 128 | 13.5 | 15.4 | 897464.2773 |
| 0 | P62851 | 40S ribosomal protein S25 OS=Homo sapiens OX=9606 GN=RPS25 PE=1 SV=1 | 48 | 7 | 7 | 7 | 125 | 13.7 | 15.22 | 723008.1455 |
| 0 | Q6PJT7 | Zinc finger CCCH domain-containing protein 14 OS=Homo sapiens OX=9606 GN=ZC3H14 PE=1 SV=1 | 10 | 5 | 5 | 5 | 736 | 82.8 | 15.16 | 121813.8906 |
| 0 | P22392 | Nucleoside diphosphate kinase B OS=Homo sapiens OX=9606 GN=NME2 PE=1 SV=1 | 57 | 6 | 7 | 3 | 152 | 17.3 | 14.71 | 435859.4199 |
| 0 | Q15291 | Retinoblastoma-binding protein 5 OS=Homo sapiens OX=9606 GN=RBBP5 PE=1 SV=2 | 15 | 5 | 5 | 5 | 538 | 59.1 | 14.69 | 159974.9277 |
| 0 | Q13895 | Bystin OS=Homo sapiens OX=9606 GN=BYSL PE=1 SV=3 | 7 | 3 | 5 | 3 | 437 | 49.6 | 14.51 | 21317.26563 |
| 0 | P52926 | High mobility group protein HMGI-C OS=Homo sapiens OX=9606 GN=HMGA2 PE=1 SV=1 | 39 | 4 | 5 | 4 | 109 | 11.8 | 14.46 | 169960.3359 |
| 0 | P52298 | Nuclear cap-binding protein subunit 2 OS=Homo sapiens OX=9606 GN=NCBP2 PE=1 SV=1 | 37 | 5 | 5 | 5 | 156 | 18 | 14.3 | 167871.209 |
| 0 | P41208 | Centrin-2 OS=Homo sapiens OX=9606 GN=CETN2 PE=1 SV=1 | 41 | 4 | 4 | 4 | 172 | 19.7 | 14.3 | 68997.0293 |
| 0 | P61981 | 14-3-3 protein gamma OS=Homo sapiens OX=9606 GN=YWHAG PE=1 SV=2 | 22 | 4 | 5 | 3 | 247 | 28.3 | 14.28 | 45925.79785 |
| 0 | Q9BRD0 | BUD13 homolog OS=Homo sapiens OX=9606 GN=BUD13 PE=1 SV=1 | 12 | 6 | 8 | 6 | 619 | 70.5 | 14.22 | 399284.3252 |
| 0 | Q6IS14 | Eukaryotic translation initiation factor 5A-1-like OS=Homo sapiens OX=9606 GN=EIF5AL1 PE=2 SV=2 | 38 | 4 | 5 | 4 | 154 | 16.8 | 14.15 | 161669.2402 |
| 0 | O60506 | Heterogeneous nuclear ribonucleoprotein Q OS=Homo sapiens OX=9606 GN=SYNCRIP PE=1 SV=2 | 9 | 5 | 5 | 5 | 623 | 69.6 | 14.06 | 94198.53125 |
| 0 | Q5BKY9 | Protein FAM133B OS=Homo sapiens OX=9606 GN=FAM133B PE=1 SV=1 | 25 | 4 | 5 | 4 | 247 | 28.4 | 14.05 | 420808.7012 |
| 0 | P63220 | 40S ribosomal protein S21 OS=Homo sapiens OX=9606 GN=RPS21 PE=1 SV=1 | 55 | 6 | 6 | 6 | 83 | 9.1 | 14.02 | 148107.3242 |
| 0 | Q86YZ3 | Hornerin OS=Homo sapiens OX=9606 GN=HRNR PE=1 SV=2 | 2 | 3 | 3 | 3 | 2850 | 282.2 | 13.76 | 15647.67383 |
| 0 | O00479 | High mobility group nucleosome-binding domain-containing protein 4 OS=Homo sapiens OX=9606 GN=HMGN4 PE=1 SV=3 | 38 | 3 | 4 | 3 | 90 | 9.5 | 13.73 | 86013.875 |
| 0 | P43487 | Ran-specific GTPase-activating protein OS=Homo sapiens OX=9606 GN=RANBP1 PE=1 SV=1 | 21 | 4 | 4 | 4 | 201 | 23.3 | 13.63 | 124342.2109 |
| 0 | P62987 | Ubiquitin-60S ribosomal protein L40 OS=Homo sapiens OX=9606 GN=UBA52 PE=1 SV=2 | 39 | 5 | 6 | 1 | 128 | 14.7 | 13.62 | |
| 0 | Q96SI1 | BTB/POZ domain-containing protein KCTD15 OS=Homo sapiens OX=9606 GN=KCTD15 PE=1 SV=1 | 13 | 3 | 4 | 3 | 283 | 31.9 | 13.61 | 45958.49414 |
| 0 | Q09028 | Histone-binding protein RBBP4 OS=Homo sapiens OX=9606 GN=RBBP4 PE=1 SV=3 | 17 | 5 | 5 | 2 | 425 | 47.6 | 13.44 | 1191082.961 |
| 0 | Q92804 | TATA-binding protein-associated factor 2N OS=Homo sapiens OX=9606 GN=TAF15 PE=1 SV=1 | 8 | 3 | 5 | 2 | 592 | 61.8 | 13.41 | 112965.0156 |
| 0 | Q13247 | Serine/arginine-rich splicing factor 6 OS=Homo sapiens OX=9606 GN=SRSF6 PE=1 SV=2 | 19 | 6 | 6 | 6 | 344 | 39.6 | 13.27 | 136676.9199 |
| 0 | O14828 | Secretory carrier-associated membrane protein 3 OS=Homo sapiens OX=9606 GN=SCAMP3 PE=1 SV=3 | 22 | 5 | 6 | 5 | 347 | 38.3 | 13.19 | 166116.5078 |
| 0 | Q01844 | RNA-binding protein EWS OS=Homo sapiens OX=9606 GN=EWSR1 PE=1 SV=1 | 7 | 3 | 4 | 3 | 656 | 68.4 | 13 | 137259.376 |
| 0 | P18077 | 60S ribosomal protein L35a OS=Homo sapiens OX=9606 GN=RPL35A PE=1 SV=2 | 31 | 6 | 6 | 6 | 110 | 12.5 | 12.9 | 221937.7891 |
| 0 | P04075 | Fructose-bisphosphate aldolase A OS=Homo sapiens OX=9606 GN=ALDOA PE=1 SV=2 | 25 | 5 | 6 | 5 | 364 | 39.4 | 12.84 | 79113.53516 |
| 0 | Q9BWD1 | Acetyl-CoA acetyltransferase, cytosolic OS=Homo sapiens OX=9606 GN=ACAT2 PE=1 SV=2 | 28 | 4 | 4 | 4 | 397 | 41.3 | 12.83 | 101493.9023 |
| 0 | P09874 | Poly [ADP-ribose] polymerase 1 OS=Homo sapiens OX=9606 GN=PARP1 PE=1 SV=4 | 7 | 4 | 4 | 4 | 1014 | 113 | 12.77 | 30959.43359 |
| 0 | Q9NZT1 | Calmodulin-like protein 5 OS=Homo sapiens OX=9606 GN=CALML5 PE=1 SV=2 | 26 | 4 | 4 | 4 | 146 | 15.9 | 12.62 | 195975.8838 |
| 0 | P83881 | 60S ribosomal protein L36a OS=Homo sapiens OX=9606 GN=RPL36A PE=1 SV=2 | 24 | 4 | 5 | 4 | 106 | 12.4 | 12.53 | 135027.2295 |
| 0 | P04406 | Glyceraldehyde-3-phosphate dehydrogenase OS=Homo sapiens OX=9606 GN=GAPDH PE=1 SV=3 | 20 | 4 | 4 | 4 | 335 | 36 | 12.42 | 79520.32031 |
| 0 | Q96J01 | THO complex subunit 3 OS=Homo sapiens OX=9606 GN=THOC3 PE=1 SV=1 | 16 | 5 | 5 | 5 | 351 | 38.7 | 12.36 | 170490.6309 |
| 0 | Q16576 | Histone-binding protein RBBP7 OS=Homo sapiens OX=9606 GN=RBBP7 PE=1 SV=1 | 17 | 5 | 5 | 2 | 425 | 47.8 | 12.34 | |
| 0 | P43243 | Matrin-3 OS=Homo sapiens OX=9606 GN=MATR3 PE=1 SV=2 | 10 | 4 | 4 | 4 | 847 | 94.6 | 12.27 | 39519.07813 |
| 0 | P22061 | Protein-L-isoaspartate(D-aspartate) O-methyltransferase OS=Homo sapiens OX=9606 GN=PCMT1 PE=1 SV=4 | 40 | 4 | 4 | 4 | 227 | 24.6 | 12.24 | 125782.3398 |
| 0 | Q07666 | KH domain-containing, RNA-binding, signal transduction-associated protein 1 OS=Homo sapiens OX=9606 GN=KHDRBS1 PE=1 SV=1 | 16 | 4 | 6 | 4 | 443 | 48.2 | 12.07 | 377732.4883 |
| 0 | Q15637 | Splicing factor 1 OS=Homo sapiens OX=9606 GN=SF1 PE=1 SV=4 | 12 | 5 | 5 | 5 | 639 | 68.3 | 12.02 | 282179.5234 |
| 0 | Q99623 | Prohibitin-2 OS=Homo sapiens OX=9606 GN=PHB2 PE=1 SV=2 | 23 | 4 | 4 | 4 | 299 | 33.3 | 11.97 | 117867.3203 |
| 0 | Q01105 | Protein SET OS=Homo sapiens OX=9606 GN=SET PE=1 SV=3 | 19 | 3 | 4 | 3 | 290 | 33.5 | 11.96 | 92800.13965 |
| 0 | P25705 | ATP synthase subunit alpha, mitochondrial OS=Homo sapiens OX=9606 GN=ATP5F1A PE=1 SV=1 | 9 | 3 | 3 | 3 | 553 | 59.7 | 11.92 | 27260.42969 |
| 0 | Q9H773 | dCTP pyrophosphatase 1 OS=Homo sapiens OX=9606 GN=DCTPP1 PE=1 SV=1 | 42 | 4 | 4 | 4 | 170 | 18.7 | 11.9 | 59643.84375 |
| 0 | Q99459 | Cell division cycle 5-like protein OS=Homo sapiens OX=9606 GN=CDC5L PE=1 SV=2 | 12 | 3 | 4 | 3 | 802 | 92.2 | 11.71 | 57213.58594 |
| 0 | P81605 | Dermcidin OS=Homo sapiens OX=9606 GN=DCD PE=1 SV=2 | 34 | 3 | 4 | 3 | 110 | 11.3 | 11.46 | 85556.78516 |
| 0 | O60232 | Protein ZNRD2 OS=Homo sapiens OX=9606 GN=ZNRD2 PE=1 SV=1 | 37 | 3 | 3 | 3 | 199 | 21.5 | 11.45 | 24051.79004 |
| 0 | Q14444 | Caprin-1 OS=Homo sapiens OX=9606 GN=CAPRIN1 PE=1 SV=2 | 12 | 4 | 4 | 4 | 709 | 78.3 | 11.43 | 95111.86816 |
| 0 | O75494 | Serine/arginine-rich splicing factor 10 OS=Homo sapiens OX=9606 GN=SRSF10 PE=1 SV=1 | 24 | 4 | 4 | 4 | 262 | 31.3 | 11.31 | 108121.2666 |
| 0 | Q6PKG0 | La-related protein 1 OS=Homo sapiens OX=9606 GN=LARP1 PE=1 SV=2 | 7 | 4 | 4 | 4 | 1096 | 123.4 | 11.19 | 28144.10449 |
| 0 | P50914 | 60S ribosomal protein L14 OS=Homo sapiens OX=9606 GN=RPL14 PE=1 SV=4 | 20 | 4 | 4 | 4 | 215 | 23.4 | 11.11 | 427553.6484 |
| 0 | Q86X55 | Histone-arginine methyltransferase CARM1 OS=Homo sapiens OX=9606 GN=CARM1 PE=1 SV=3 | 9 | 4 | 4 | 4 | 608 | 65.8 | 11.02 | 52766.64941 |
| 0 | P62913 | 60S ribosomal protein L11 OS=Homo sapiens OX=9606 GN=RPL11 PE=1 SV=2 | 20 | 3 | 4 | 3 | 178 | 20.2 | 10.88 | 205716.2344 |

| | | | | | | | | | | |
|---|---|---|---|---|---|---|---|---|---|---|
| 0 | P05023 | Sodium/potassium-transporting ATPase subunit alpha-1 OS=Homo sapiens OX=9606 GN=ATP1A1 PE=1 SV=1 | 6 | 4 | 4 | 4 | 1023 | 112.8 | 10.8 | 99177.63867 |
| 0 | P56270 | Myc-associated zinc finger protein OS=Homo sapiens OX=9606 GN=MAZ PE=1 SV=1 | 9 | 3 | 4 | 3 | 477 | 48.6 | 10.76 | 36305.91406 |
| 0 | Q9BVG4 | Protein PBDC1 OS=Homo sapiens OX=9606 GN=PBDC1 PE=1 SV=1 | 17 | 3 | 4 | 3 | 233 | 26 | 10.72 | 41315.68457 |
| 0 | P04080 | Cystatin-B OS=Homo sapiens OX=9606 GN=CSTB PE=1 SV=2 | 58 | 3 | 4 | 3 | 98 | 11.1 | 10.59 | 43457.22852 |
| 0 | P46459 | Vesicle-fusing ATPase OS=Homo sapiens OX=9606 GN=NSF PE=1 SV=3 | 6 | 4 | 4 | 4 | 744 | 82.5 | 10.43 | 155202.0664 |
| 0 | P47755 | F-actin-capping protein subunit alpha-2 OS=Homo sapiens OX=9606 GN=CAPZA2 PE=1 SV=3 | 17 | 4 | 4 | 2 | 286 | 32.9 | 10.43 | 60627.7793 |
| 0 | P41091 | Eukaryotic translation initiation factor 2 subunit 3 OS=Homo sapiens OX=9606 GN=EIF2S3 PE=1 SV=3 | 10 | 3 | 4 | 3 | 472 | 51.1 | 10.29 | 39504.29102 |
| 0 | Q96EB1 | Elongator complex protein 4 OS=Homo sapiens OX=9606 GN=ELP4 PE=1 SV=2 | 12 | 4 | 5 | 4 | 424 | 46.6 | 10.23 | 49228.80957 |
| 0 | Q96CN7 | Isochorismatase domain-containing protein 1 OS=Homo sapiens OX=9606 GN=ISOC1 PE=1 SV=3 | 19 | 5 | 5 | 5 | 298 | 32.2 | 10.19 | 122233.5771 |
| 0 | Q9P0U4 | CXXC-type zinc finger protein 1 OS=Homo sapiens OX=9606 GN=CXXC1 PE=1 SV=2 | 5 | 2 | 4 | 2 | 656 | 75.7 | 10.11 | 43936.0625 |
| 0 | P55735 | Protein SEC13 homolog OS=Homo sapiens OX=9606 GN=SEC13 PE=1 SV=3 | 19 | 3 | 3 | 3 | 322 | 35.5 | 10.06 | 179467.5381 |
| 0 | P26641 | Elongation factor 1-gamma OS=Homo sapiens OX=9606 GN=EEF1G PE=1 SV=3 | 12 | 4 | 4 | 4 | 437 | 50.1 | 9.91 | 77502.93896 |
| 0 | P61289 | Proteasome activator complex subunit 3 OS=Homo sapiens OX=9606 GN=PSME3 PE=1 SV=1 | 16 | 3 | 3 | 3 | 254 | 29.5 | 9.88 | 61182.14746 |
| 0 | Q5SSJ5 | Heterochromatin protein 1-binding protein 3 OS=Homo sapiens OX=9606 GN=HP1BP3 PE=1 SV=1 | 9 | 3 | 3 | 3 | 553 | 61.2 | 9.79 | 40144.55469 |
| 0 | Q9BWJ5 | Splicing factor 3B subunit 5 OS=Homo sapiens OX=9606 GN=SF3B5 PE=1 SV=1 | 60 | 3 | 4 | 3 | 86 | 10.1 | 9.66 | 214830.2305 |
| 0 | Q8N9Q2 | Protein SREK1IP1 OS=Homo sapiens OX=9606 GN=SREK1IP1 PE=1 SV=1 | 25 | 4 | 4 | 4 | 155 | 18.2 | 9.62 | 279425.4883 |
| 0 | P39023 | 60S ribosomal protein L3 OS=Homo sapiens OX=9606 GN=RPL3 PE=1 SV=2 | 12 | 3 | 3 | 3 | 403 | 46.1 | 9.49 | 48338.62402 |
| 0 | P62861 | 40S ribosomal protein S30 OS=Homo sapiens OX=9606 GN=FAU PE=1 SV=1 | 34 | 5 | 7 | 5 | 59 | 6.6 | 9.49 | 1689896.92 |
| 0 | Q92841 | Probable ATP-dependent RNA helicase DDX17 OS=Homo sapiens OX=9606 GN=DDX17 PE=1 SV=2 | 4 | 2 | 3 | 1 | 729 | 80.2 | 9.34 | 21046.22656 |
| 0 | P05114 | Non-histone chromosomal protein HMG-14 OS=Homo sapiens OX=9606 GN=HMGN1 PE=1 SV=3 | 28 | 2 | 4 | 2 | 100 | 10.7 | 9.16 | 215095.0156 |
| 0 | Q9UMS4 | Pre-mRNA-processing factor 19 OS=Homo sapiens OX=9606 GN=PRPF19 PE=1 SV=1 | 16 | 4 | 5 | 4 | 504 | 55.1 | 9.13 | 48675.27148 |
| 0 | O75396 | Vesicle-trafficking protein SEC22b OS=Homo sapiens OX=9606 GN=SEC22B PE=1 SV=4 | 30 | 3 | 3 | 3 | 215 | 24.6 | 9.04 | 15444.61035 |
| 0 | P15531 | Nucleoside diphosphate kinase A OS=Homo sapiens OX=9606 GN=NME1 PE=1 SV=1 | 34 | 4 | 5 | 1 | 152 | 17.1 | 9.03 | |
| 0 | Q71UM5 | 40S ribosomal protein S27-like OS=Homo sapiens OX=9606 GN=RPS27L PE=1 SV=3 | 40 | 3 | 3 | 1 | 84 | 9.5 | 8.92 | 35833.41797 |
| 0 | P17987 | T-complex protein 1 subunit alpha OS=Homo sapiens OX=9606 GN=TCP1 PE=1 SV=1 | 8 | 3 | 3 | 3 | 556 | 60.3 | 8.92 | 30735.2627 |
| 0 | O76021 | Ribosomal L1 domain-containing protein 1 OS=Homo sapiens OX=9606 GN=RSL1D1 PE=1 SV=3 | 10 | 4 | 4 | 4 | 490 | 54.9 | 8.72 | 93170.93555 |
| 0 | Q7Z4W1 | L-xylulose reductase OS=Homo sapiens OX=9606 GN=DCXR PE=1 SV=2 | 9 | 2 | 3 | 2 | 244 | 25.9 | 8.72 | |
| 0 | P02786 | Transferrin receptor protein 1 OS=Homo sapiens OX=9606 GN=TFRC PE=1 SV=2 | 6 | 3 | 3 | 3 | 760 | 84.8 | 8.68 | 102754.2813 |
| 0 | P08590 | Myosin light chain 3 OS=Homo sapiens OX=9606 GN=MYL3 PE=1 SV=3 | 8 | 1 | 2 | 1 | 195 | 21.9 | 8.59 | 280779.5 |
| 0 | Q15370 | Elongin-B OS=Homo sapiens OX=9606 GN=ELOB PE=1 SV=1 | 31 | 1 | 2 | 1 | 118 | 13.1 | 8.47 | 31645.17188 |
| 0 | O60832 | H/ACA ribonucleoprotein complex subunit DKC1 OS=Homo sapiens OX=9606 GN=DKC1 PE=1 SV=3 | 8 | 3 | 3 | 3 | 514 | 57.6 | 8.47 | 25299.92578 |
| 0 | P11441 | Ubiquitin-like protein 4A OS=Homo sapiens OX=9606 GN=UBL4A PE=1 SV=1 | 24 | 3 | 3 | 3 | 157 | 17.8 | 8.45 | 58390.49805 |
| 0 | Q969G3 | SWI/SNF-related matrix-associated actin-dependent regulator of chromatin subfamily E member 1 OS=Homo sapiens OX=9606 GN=SMARCE1 PE=1 SV=2 | 13 | 2 | 2 | 2 | 411 | 46.6 | 8.24 | 16317.84863 |
| 0 | P17096 | High mobility group protein HMG-I/HMG-Y OS=Homo sapiens OX=9606 GN=HMGA1 PE=1 SV=3 | 38 | 2 | 4 | 2 | 107 | 11.7 | 8.21 | 178673.3906 |
| 0 | E9PAV3 | Nascent polypeptide-associated complex subunit alpha, muscle-specific form OS=Homo sapiens OX=9606 GN=NACA PE=1 SV=1 | 3 | 3 | 3 | 3 | 2078 | 205.3 | 8.06 | 28122.70703 |
| 0 | Q92928 | Putative Ras-related protein Rab-1C OS=Homo sapiens OX=9606 GN=RAB1C PE=5 SV=2 | 14 | 2 | 3 | 1 | 201 | 22 | 8.03 | 11898.05469 |
| 0 | Q8WUD4 | Coiled-coil domain-containing protein 12 OS=Homo sapiens OX=9606 GN=CCDC12 PE=1 SV=1 | 30 | 3 | 3 | 3 | 166 | 19.2 | 7.92 | 63613.80859 |
| 0 | P06730 | Eukaryotic translation initiation factor 4E OS=Homo sapiens OX=9606 GN=EIF4E PE=1 SV=2 | 18 | 4 | 4 | 4 | 217 | 25.1 | 7.9 | 127330.9355 |
| 0 | P27635 | 60S ribosomal protein L10 OS=Homo sapiens OX=9606 GN=RPL10 PE=1 SV=4 | 22 | 2 | 2 | 2 | 214 | 24.6 | 7.86 | |
| 0 | Q13616 | Cullin-1 OS=Homo sapiens OX=9606 GN=CUL1 PE=1 SV=2 | 5 | 3 | 3 | 3 | 776 | 89.6 | 7.56 | 116548.1563 |
| 0 | P62136 | Serine/threonine-protein phosphatase PP1-alpha catalytic subunit OS=Homo sapiens OX=9606 GN=PPP1CA PE=1 SV=1 | 11 | 3 | 3 | 2 | 330 | 37.5 | 7.47 | 62614.7959 |
| 0 | P36578 | 60S ribosomal protein L4 OS=Homo sapiens OX=9606 GN=RPL4 PE=1 SV=5 | 8 | 2 | 2 | 2 | 427 | 47.7 | 7.45 | 31581.7793 |
| 0 | P50990 | T-complex protein 1 subunit theta OS=Homo sapiens OX=9606 GN=CCT8 PE=1 SV=4 | 6 | 3 | 3 | 3 | 548 | 59.6 | 7.39 | 15717.36719 |
| 0 | P22087 | rRNA 2'-O-methyltransferase fibrillarin OS=Homo sapiens OX=9606 GN=FBL PE=1 SV=2 | 16 | 4 | 4 | 4 | 321 | 33.8 | 7.37 | 233846.8281 |
| 0 | P05783 | Keratin, type I cytoskeletal 18 OS=Homo sapiens OX=9606 GN=KRT18 PE=1 SV=2 | 21 | 5 | 5 | 5 | 430 | 48 | 7.23 | 70882.30273 |
| 0 | Q13283 | Ras GTPase-activating protein-binding protein 1 OS=Homo sapiens OX=9606 GN=G3BP1 PE=1 SV=1 | 8 | 2 | 2 | 2 | 466 | 52.1 | 7.22 | 25668.80859 |
| 0 | P62820 | Ras-related protein Rab-1A OS=Homo sapiens OX=9606 GN=RAB1A PE=1 SV=3 | 13 | 2 | 4 | 1 | 205 | 22.7 | 7.18 | 46073.51953 |
| 0 | P05386 | 60S acidic ribosomal protein P1 OS=Homo sapiens OX=9606 GN=RPLP1 PE=1 SV=1 | 14 | 1 | 2 | 1 | 114 | 11.5 | 7.16 | 328107.1563 |
| 0 | P62306 | Small nuclear ribonucleoprotein F OS=Homo sapiens OX=9606 GN=SNRPF PE=1 SV=1 | 41 | 2 | 3 | 2 | 86 | 9.7 | 7.15 | 130044.3477 |
| 0 | Q99714 | 3-hydroxyacyl-CoA dehydrogenase type-2 OS=Homo sapiens OX=9606 GN=HSD17B10 PE=1 SV=3 | 19 | 2 | 2 | 2 | 261 | 26.9 | 7.1 | 18554.1875 |
| 0 | Q7Z3C6 | Autophagy-related protein 9A OS=Homo sapiens OX=9606 GN=ATG9A PE=1 SV=3 | 8 | 3 | 3 | 3 | 839 | 94.4 | 7.04 | 62966.2168 |
| 0 | Q15836 | Vesicle-associated membrane protein 3 OS=Homo sapiens OX=9606 GN=VAMP3 PE=1 SV=3 | 25 | 2 | 2 | 2 | 100 | 11.3 | 7.02 | 44900.25 |
| 0 | P35613 | Basigin OS=Homo sapiens OX=9606 GN=BSG PE=1 SV=2 | 10 | 3 | 3 | 3 | 385 | 42.2 | 6.99 | 25322.21094 |
| 0 | P30153 | Serine/threonine-protein phosphatase 2A 65 kDa regulatory subunit A alpha isoform OS=Homo sapiens OX=9606 GN=PPP2R1A PE=1 SV=4 | 4 | 2 | 3 | 2 | 589 | 65.3 | 6.98 | 18220.44727 |
| 0 | P08195 | 4F2 cell-surface antigen heavy chain OS=Homo sapiens OX=9606 GN=SLC3A2 PE=1 SV=3 | 4 | 2 | 3 | 2 | 630 | 68 | 6.94 | 20650.20117 |
| 0 | P61353 | 60S ribosomal protein L27 OS=Homo sapiens OX=9606 GN=RPL27 PE=1 SV=2 | 34 | 5 | 6 | 5 | 136 | 15.8 | 6.9 | 742614.0527 |
| 0 | Q9UHB9 | Signal recognition particle subunit SRP68 OS=Homo sapiens OX=9606 GN=SRP68 PE=1 SV=2 | 7 | 2 | 3 | 2 | 627 | 70.7 | 6.89 | 111259.5859 |
| 0 | P49207 | 60S ribosomal protein L34 OS=Homo sapiens OX=9606 GN=RPL34 PE=1 SV=3 | 32 | 4 | 4 | 4 | 117 | 13.3 | 6.87 | 225828.9961 |
| 0 | Q8N5F7 | NF-kappa-B-activating protein OS=Homo sapiens OX=9606 GN=NKAP PE=1 SV=1 | 4 | 1 | 2 | 1 | 415 | 47.1 | 6.79 | 19694.29102 |

| | | | | | | | | | | |
|---|---|---|---|---|---|---|---|---|---|---|
| 0 | O43684 | Mitotic checkpoint protein BUB3 OS=Homo sapiens OX=9606 GN=BUB3 PE=1 SV=1 | 7 | 2 | 3 | 2 | 328 | 37.1 | 6.76 | 13113.8916 |
| 0 | P18124 | 60S ribosomal protein L7 OS=Homo sapiens OX=9606 GN=RPL7 PE=1 SV=1 | 12 | 2 | 2 | 2 | 248 | 29.2 | 6.72 | 66516.79688 |
| 0 | Q15651 | High mobility group nucleosome-binding domain-containing protein 3 OS=Homo sapiens OX=9606 GN=HMGN3 PE=1 SV=2 | 15 | 1 | 5 | 1 | 99 | 10.7 | 6.7 | 181627.0293 |
| 0 | Q96HQ2 | CDKN2AIP N-terminal-like protein OS=Homo sapiens OX=9606 GN=CDKN2AIPNL PE=1 SV=1 | 24 | 2 | 2 | 2 | 116 | 13.2 | 6.69 | 51613.74219 |
| 0 | P05388 | 60S acidic ribosomal protein P0 OS=Homo sapiens OX=9606 GN=RPLP0 PE=1 SV=1 | 10 | 2 | 2 | 2 | 317 | 34.3 | 6.66 | 62271.86328 |
| 0 | Q15428 | Splicing factor 3A subunit 2 OS=Homo sapiens OX=9606 GN=SF3A2 PE=1 SV=2 | 7 | 3 | 3 | 3 | 464 | 49.2 | 6.65 | 24767.90137 |
| 0 | P62241 | 40S ribosomal protein S8 OS=Homo sapiens OX=9606 GN=RPS8 PE=1 SV=2 | 13 | 2 | 2 | 2 | 208 | 24.2 | 6.59 | 98451.71094 |
| 0 | Q15437 | Protein transport protein Sec23B OS=Homo sapiens OX=9606 GN=SEC23B PE=1 SV=2 | 7 | 3 | 3 | 3 | 767 | 86.4 | 6.56 | 64827.98926 |
| 0 | Q09161 | Nuclear cap-binding protein subunit 1 OS=Homo sapiens OX=9606 GN=NCBP1 PE=1 SV=1 | 5 | 2 | 2 | 2 | 790 | 91.8 | 6.52 | 25750.97852 |
| 0 | Q15365 | Poly(rC)-binding protein 1 OS=Homo sapiens OX=9606 GN=PCBP1 PE=1 SV=2 | 10 | 2 | 2 | 2 | 356 | 37.5 | 6.52 | 16205.99707 |
| 0 | Q9P013 | Spliceosome-associated protein CWC15 homolog OS=Homo sapiens OX=9606 GN=CWC15 PE=1 SV=2 | 22 | 3 | 3 | 3 | 229 | 26.6 | 6.46 | 118347.2031 |
| 0 | P02792 | Ferritin light chain OS=Homo sapiens OX=9606 GN=FTL PE=1 SV=2 | 14 | 2 | 2 | 2 | 175 | 20 | 6.43 | 33796.07813 |
| 0 | Q9H3K6 | BolA-like protein 2 OS=Homo sapiens OX=9606 GN=BOLA2 PE=1 SV=1 | 29 | 2 | 2 | 2 | 86 | 10.1 | 6.42 | 49127.75781 |
| 0 | P51572 | B-cell receptor-associated protein 31 OS=Homo sapiens OX=9606 GN=BCAP31 PE=1 SV=3 | 9 | 2 | 2 | 2 | 246 | 28 | 6.42 | 141693.3867 |
| 0 | Q01130 | Serine/arginine-rich splicing factor 2 OS=Homo sapiens OX=9606 GN=SRSF2 PE=1 SV=4 | 16 | 2 | 2 | 2 | 221 | 25.5 | 6.38 | 39864.01074 |
| 0 | P63000 | Ras-related C3 botulinum toxin substrate 1 OS=Homo sapiens OX=9606 GN=RAC1 PE=1 SV=1 | 13 | 2 | 2 | 2 | 192 | 21.4 | 6.38 | 22633.41016 |
| 0 | P46782 | 40S ribosomal protein S5 OS=Homo sapiens OX=9606 GN=RPS5 PE=1 SV=4 | 11 | 2 | 2 | 2 | 204 | 22.9 | 6.16 | 20560.7832 |
| 0 | Q7RTV0 | PHD finger-like domain-containing protein 5A OS=Homo sapiens OX=9606 GN=PHF5A PE=1 SV=1 | 25 | 2 | 2 | 2 | 110 | 12.4 | 6.14 | 80003.94922 |
| 0 | O00264 | Membrane-associated progesterone receptor component 1 OS=Homo sapiens OX=9606 GN=PGRMC1 PE=1 SV=3 | 8 | 2 | 2 | 2 | 195 | 21.7 | 6.14 | 65654.20313 |
| 0 | Q9H814 | Phosphorylated adapter RNA export protein OS=Homo sapiens OX=9606 GN=PHAX PE=1 SV=1 | 7 | 2 | 2 | 2 | 394 | 44.4 | 6.14 | |
| 0 | Q99700 | Ataxin-2 OS=Homo sapiens OX=9606 GN=ATXN2 PE=1 SV=2 | 3 | 3 | 3 | 3 | 1313 | 140.2 | 6.12 | |
| 0 | P04062 | Lysosomal acid glucosylceramidase OS=Homo sapiens OX=9606 GN=GBA PE=1 SV=3 | 4 | 1 | 2 | 1 | 536 | 59.7 | 6.1 | |
| 0 | Q3MHD2 | Protein LSM12 homolog OS=Homo sapiens OX=9606 GN=LSM12 PE=1 SV=2 | 11 | 2 | 2 | 2 | 195 | 21.7 | 6.06 | 80250.77344 |
| 0 | Q9UN86 | Ras GTPase-activating protein-binding protein 2 OS=Homo sapiens OX=9606 GN=G3BP2 PE=1 SV=2 | 12 | 3 | 3 | 3 | 482 | 54.1 | 6.04 | 40098.49805 |
| 0 | P06702 | Protein S100-A9 OS=Homo sapiens OX=9606 GN=S100A9 PE=1 SV=1 | 25 | 2 | 2 | 2 | 114 | 13.2 | 6.01 | 122695.6953 |
| 0 | Q9NVI7 | ATPase family AAA domain-containing protein 3A OS=Homo sapiens OX=9606 GN=ATAD3A PE=1 SV=2 | 6 | 3 | 3 | 3 | 634 | 71.3 | 5.96 | 16611.19531 |
| 0 | O75934 | Pre-mRNA-splicing factor SPF27 OS=Homo sapiens OX=9606 GN=BCAS2 PE=1 SV=1 | 24 | 3 | 3 | 3 | 225 | 26.1 | 5.93 | 42557.60254 |
| 0 | Q00325 | Phosphate carrier protein, mitochondrial OS=Homo sapiens OX=9606 GN=SLC25A3 PE=1 SV=2 | 3 | 1 | 2 | 1 | 362 | 40.1 | 5.91 | 46842.28516 |
| 0 | P78371 | T-complex protein 1 subunit beta OS=Homo sapiens OX=9606 GN=CCT2 PE=1 SV=4 | 5 | 2 | 2 | 2 | 535 | 57.5 | 5.91 | |
| 0 | Q9H2H8 | Peptidyl-prolyl cis-trans isomerase-like 3 OS=Homo sapiens OX=9606 GN=PPIL3 PE=1 SV=1 | 20 | 2 | 3 | 2 | 161 | 18.1 | 5.85 | 68392.2168 |
| 0 | P20674 | Cytochrome c oxidase subunit 5A, mitochondrial OS=Homo sapiens OX=9606 GN=COX5A PE=1 SV=2 | 18 | 2 | 2 | 2 | 150 | 16.8 | 5.84 | 61754.31055 |
| 0 | Q13601 | KRR1 small subunit processome component homolog OS=Homo sapiens OX=9606 GN=KRR1 PE=1 SV=4 | 7 | 2 | 2 | 2 | 381 | 43.6 | 5.81 | 18659.7793 |
| 0 | O60828 | Polyglutamine-binding protein 1 OS=Homo sapiens OX=9606 GN=PQBP1 PE=1 SV=1 | 20 | 3 | 3 | 3 | 265 | 30.5 | 5.78 | 11172.91992 |
| 0 | P52815 | 39S ribosomal protein L12, mitochondrial OS=Homo sapiens OX=9606 GN=MRPL12 PE=1 SV=2 | 11 | 2 | 2 | 2 | 198 | 21.3 | 5.76 | 42270.69141 |
| 0 | O75367 | Core histone macro-H2A.1 OS=Homo sapiens OX=9606 GN=MACROH2A1 PE=1 SV=4 | 6 | 1 | 2 | 1 | 372 | 39.6 | 5.76 | 17169.12305 |
| 0 | P09211 | Glutathione S-transferase P OS=Homo sapiens OX=9606 GN=GSTP1 PE=1 SV=2 | 15 | 2 | 2 | 2 | 210 | 23.3 | 5.72 | 73508.2334 |
| 0 | Q14103 | Heterogeneous nuclear ribonucleoprotein D0 OS=Homo sapiens OX=9606 GN=HNRNPD PE=1 SV=1 | 6 | 2 | 2 | 2 | 355 | 38.4 | 5.72 | 35717.21094 |
| 0 | P27348 | 14-3-3 protein theta OS=Homo sapiens OX=9606 GN=YWHAQ PE=1 SV=1 | 9 | 2 | 2 | 1 | 245 | 27.7 | 5.7 | 16096.93457 |
| 0 | P30041 | Peroxiredoxin-6 OS=Homo sapiens OX=9606 GN=PRDX6 PE=1 SV=3 | 15 | 2 | 2 | 2 | 224 | 25 | 5.69 | 45316.72852 |
| 0 | O76094 | Signal recognition particle subunit SRP72 OS=Homo sapiens OX=9606 GN=SRP72 PE=1 SV=3 | 4 | 1 | 2 | 1 | 671 | 74.6 | 5.68 | |
| 0 | P62314 | Small nuclear ribonucleoprotein Sm D1 OS=Homo sapiens OX=9606 GN=SNRPD1 PE=1 SV=1 | 29 | 2 | 2 | 2 | 119 | 13.3 | 5.62 | 157598.6255 |
| 0 | P16615 | Sarcoplasmic/endoplasmic reticulum calcium ATPase 2 OS=Homo sapiens OX=9606 GN=ATP2A2 PE=1 SV=1 | 3 | 2 | 2 | 2 | 1042 | 114.7 | 5.61 | |
| 0 | Q15427 | Splicing factor 3B subunit 4 OS=Homo sapiens OX=9606 GN=SF3B4 PE=1 SV=1 | 6 | 2 | 2 | 2 | 424 | 44.4 | 5.61 | 34347.53223 |
| 0 | P33316 | Deoxyuridine 5'-triphosphate nucleotidohydrolase, mitochondrial OS=Homo sapiens OX=9606 GN=DUT PE=1 SV=4 | 8 | 2 | 2 | 2 | 252 | 26.5 | 5.61 | 104326.334 |
| 0 | P63208 | S-phase kinase-associated protein 1 OS=Homo sapiens OX=9606 GN=SKP1 PE=1 SV=2 | 23 | 2 | 3 | 2 | 163 | 18.6 | 5.6 | 181030.0703 |
| 0 | P45880 | Voltage-dependent anion-selective channel protein 2 OS=Homo sapiens OX=9606 GN=VDAC2 PE=1 SV=2 | 9 | 2 | 2 | 2 | 294 | 31.5 | 5.59 | 52911.33789 |
| 0 | P62244 | 40S ribosomal protein S15a OS=Homo sapiens OX=9606 GN=RPS15A PE=1 SV=2 | 17 | 2 | 2 | 2 | 130 | 14.8 | 5.55 | 144426.4063 |
| 0 | P62633 | Cellular nucleic acid-binding protein OS=Homo sapiens OX=9606 GN=CNBP PE=1 SV=1 | 8 | 1 | 2 | 1 | 177 | 19.5 | 5.55 | |
| 0 | P04843 | Dolichyl-diphosphooligosaccharide--protein glycosyltransferase subunit 1 OS=Homo sapiens OX=9606 GN=RPN1 PE=1 SV=1 | 4 | 2 | 2 | 2 | 607 | 68.5 | 5.54 | 51066.16406 |
| 0 | Q9NVT9 | Armadillo repeat-containing protein 1 OS=Homo sapiens OX=9606 GN=ARMC1 PE=1 SV=1 | 7 | 1 | 2 | 1 | 282 | 31.3 | 5.51 | 47476.37988 |
| 0 | Q99959 | Plakophilin-2 OS=Homo sapiens OX=9606 GN=PKP2 PE=1 SV=2 | 4 | 2 | 2 | 2 | 881 | 97.4 | 5.43 | |
| 0 | Q92598 | Heat shock protein 105 kDa OS=Homo sapiens OX=9606 GN=HSPH1 PE=1 SV=1 | 3 | 2 | 2 | 2 | 858 | 96.8 | 5.42 | 10563.05273 |
| 0 | P63167 | Dynein light chain 1, cytoplasmic OS=Homo sapiens OX=9606 GN=DYNLL1 PE=1 SV=1 | 37 | 2 | 2 | 2 | 89 | 10.4 | 5.42 | 58054.15625 |
| 0 | Q16891 | MICOS complex subunit MIC60 OS=Homo sapiens OX=9606 GN=IMMT PE=1 SV=1 | 3 | 2 | 2 | 2 | 758 | 83.6 | 5.41 | 13917.07715 |
| 0 | Q9H6F5 | Coiled-coil domain-containing protein 86 OS=Homo sapiens OX=9606 GN=CCDC86 PE=1 SV=1 | 7 | 2 | 2 | 2 | 360 | 40.2 | 5.4 | 56117.87305 |
| 0 | Q9Y4Z0 | U6 snRNA-associated Sm-like protein LSm4 OS=Homo sapiens OX=9606 GN=LSM4 PE=1 SV=1 | 20 | 2 | 2 | 2 | 139 | 15.3 | 5.36 | 36882.4707 |
| 0 | O15212 | Prefoldin subunit 6 OS=Homo sapiens OX=9606 GN=PFDN6 PE=1 SV=1 | 19 | 2 | 2 | 2 | 129 | 14.6 | 5.29 | 9331.125977 |
| 0 | Q13148 | TAR DNA-binding protein 43 OS=Homo sapiens OX=9606 GN=TARDBP PE=1 SV=1 | 7 | 2 | 3 | 2 | 414 | 44.7 | 5.23 | 11855.24316 |

| | | | | | | | | | | |
|---|---|---|---|---|---|---|---|---|---|---|
| 0 | P36873 | Serine/threonine-protein phosphatase PP1-gamma catalytic subunit OS=Homo sapiens OX=9606 GN=PPP1CC PE=1 SV=1 | 8 | 2 | 2 | 1 | 323 | 37 | 5.2 | 29161.04688 |
| 0 | Q13595 | Transformer-2 protein homolog alpha OS=Homo sapiens OX=9606 GN=TRA2A PE=1 SV=1 | 7 | 2 | 2 | 2 | 282 | 32.7 | 5.18 | 35932.56445 |
| 0 | P26368 | Splicing factor U2AF 65 kDa subunit OS=Homo sapiens OX=9606 GN=U2AF2 PE=1 SV=4 | 9 | 2 | 2 | 2 | 475 | 53.5 | 5.14 | 42166.90234 |
| 0 | Q99832 | T-complex protein 1 subunit eta OS=Homo sapiens OX=9606 GN=CCT7 PE=1 SV=2 | 7 | 2 | 2 | 2 | 543 | 59.3 | 5.11 | 8941.390625 |
| 0 | Q9HB71 | Calcyclin-binding protein OS=Homo sapiens OX=9606 GN=CACYBP PE=1 SV=2 | 8 | 1 | 1 | 1 | 228 | 26.2 | 5.11 | |
| 0 | Q04917 | 14-3-3 protein eta OS=Homo sapiens OX=9606 GN=YWHAH PE=1 SV=4 | 15 | 2 | 2 | 2 | 246 | 28.2 | 5.09 | |
| 0 | P27824 | Calnexin OS=Homo sapiens OX=9606 GN=CANX PE=1 SV=2 | 5 | 2 | 2 | 2 | 592 | 67.5 | 5.06 | 78053.26563 |
| 0 | P37198 | Nuclear pore glycoprotein p62 OS=Homo sapiens OX=9606 GN=NUP62 PE=1 SV=3 | 6 | 2 | 2 | 2 | 522 | 53.2 | 5.05 | 24323.97363 |
| 0 | O15047 | Histone-lysine N-methyltransferase SETD1A OS=Homo sapiens OX=9606 GN=SETD1A PE=1 SV=3 | 2 | 2 | 2 | 2 | 1707 | 185.9 | 5.05 | 26369.48535 |
| 0 | P56537 | Eukaryotic translation initiation factor 6 OS=Homo sapiens OX=9606 GN=EIF6 PE=1 SV=1 | 14 | 2 | 2 | 2 | 245 | 26.6 | 5.03 | 42750.43945 |
| 0 | P60174 | Triosephosphate isomerase OS=Homo sapiens OX=9606 GN=TPI1 PE=1 SV=4 | 11 | 2 | 2 | 2 | 249 | 26.7 | 5.01 | |
| 0 | P09234 | U1 small nuclear ribonucleoprotein C OS=Homo sapiens OX=9606 GN=SNRPC PE=1 SV=1 | 13 | 1 | 1 | 1 | 159 | 17.4 | 4.96 | 14448.96875 |
| 0 | Q14978 | Nucleolar and coiled-body phosphoprotein 1 OS=Homo sapiens OX=9606 GN=NOLC1 PE=1 SV=2 | 4 | 2 | 2 | 2 | 699 | 73.6 | 4.95 | 54652.44531 |
| 0 | P10599 | Thioredoxin OS=Homo sapiens OX=9606 GN=TXN PE=1 SV=3 | 30 | 2 | 2 | 2 | 105 | 11.7 | 4.95 | 123103.2188 |
| 0 | P62304 | Small nuclear ribonucleoprotein E OS=Homo sapiens OX=9606 GN=SNRPE PE=1 SV=1 | 29 | 2 | 2 | 2 | 92 | 10.8 | 4.94 | 154205.0469 |
| 0 | Q9ULR0 | Pre-mRNA-splicing factor ISY1 homolog OS=Homo sapiens OX=9606 GN=ISY1 PE=1 SV=3 | 9 | 2 | 2 | 2 | 285 | 33 | 4.9 | 28455.94238 |
| 0 | Q9NPA8 | Transcription and mRNA export factor ENY2 OS=Homo sapiens OX=9606 GN=ENY2 PE=1 SV=1 | 20 | 2 | 2 | 2 | 101 | 11.5 | 4.89 | 14311.42871 |
| 0 | P62854 | 40S ribosomal protein S26 OS=Homo sapiens OX=9606 GN=RPS26 PE=1 SV=3 | 19 | 2 | 3 | 2 | 115 | 13 | 4.86 | 40231.64063 |
| 0 | Q13162 | Peroxiredoxin-4 OS=Homo sapiens OX=9606 GN=PRDX4 PE=1 SV=1 | 9 | 2 | 2 | 1 | 271 | 30.5 | 4.82 | 24154.03125 |
| 0 | P46379 | Large proline-rich protein BAG6 OS=Homo sapiens OX=9606 GN=BAG6 PE=1 SV=2 | 4 | 3 | 3 | 3 | 1132 | 119.3 | 4.8 | 75728.59766 |
| 0 | O75607 | Nucleoplasmin-3 OS=Homo sapiens OX=9606 GN=NPM3 PE=1 SV=3 | 10 | 1 | 1 | 1 | 178 | 19.3 | 4.75 | 23295.53711 |
| 0 | Q04837 | Single-stranded DNA-binding protein, mitochondrial OS=Homo sapiens OX=9606 GN=SSBP1 PE=1 SV=1 | 16 | 2 | 2 | 2 | 148 | 17.2 | 4.7 | 111196.0742 |
| 0 | P01834 | Immunoglobulin kappa constant OS=Homo sapiens OX=9606 GN=IGKC PE=1 SV=2 | 19 | 1 | 1 | 1 | 107 | 11.8 | 4.7 | 11684.37207 |
| 0 | Q9NY12 | H/ACA ribonucleoprotein complex subunit 1 OS=Homo sapiens OX=9606 GN=GAR1 PE=1 SV=1 | 14 | 2 | 2 | 2 | 217 | 22.3 | 4.69 | 40788.65332 |
| 0 | O14579 | Coatomer subunit epsilon OS=Homo sapiens OX=9606 GN=COPE PE=1 SV=3 | 11 | 2 | 2 | 2 | 308 | 34.5 | 4.64 | 16442.27539 |
| 0 | Q9BQG0 | Myb-binding protein 1A OS=Homo sapiens OX=9606 GN=MYBBP1A PE=1 SV=2 | 3 | 2 | 2 | 2 | 1328 | 148.8 | 4.6 | 12183.59082 |
| 0 | P61244 | Protein max OS=Homo sapiens OX=9606 GN=MAX PE=1 SV=1 | 9 | 1 | 1 | 1 | 160 | 18.3 | 4.55 | 15317.17773 |
| 0 | Q6RW13 | Type-1 angiotensin II receptor-associated protein OS=Homo sapiens OX=9606 GN=AGTRAP PE=1 SV=1 | 14 | 1 | 1 | 1 | 159 | 17.4 | 4.55 | 60828.23828 |
| 0 | Q96BT3 | Centromere protein T OS=Homo sapiens OX=9606 GN=CENPT PE=1 SV=2 | 2 | 1 | 2 | 1 | 561 | 60.4 | 4.52 | 5283455 |
| 0 | Q13557 | Calcium/calmodulin-dependent protein kinase type II subunit delta OS=Homo sapiens OX=9606 GN=CAMK2D PE=1 SV=3 | 4 | 2 | 2 | 2 | 499 | 56.3 | 4.52 | 62813.61133 |
| 0 | P62266 | 40S ribosomal protein S23 OS=Homo sapiens OX=9606 GN=RPS23 PE=1 SV=3 | 15 | 2 | 2 | 2 | 143 | 15.8 | 4.49 | 112799.418 |
| 0 | P62308 | Small nuclear ribonucleoprotein G OS=Homo sapiens OX=9606 GN=SNRPG PE=1 SV=1 | 26 | 2 | 2 | 2 | 76 | 8.5 | 4.49 | 206747.3438 |
| 0 | P34897 | Serine hydroxymethyltransferase, mitochondrial OS=Homo sapiens OX=9606 GN=SHMT2 PE=1 SV=3 | 8 | 1 | 1 | 1 | 504 | 56 | 4.44 | |
| 0 | Q8WXF1 | Paraspeckle component 1 OS=Homo sapiens OX=9606 GN=PSPC1 PE=1 SV=1 | 3 | 1 | 2 | 1 | 523 | 58.7 | 4.43 | 16368.03711 |
| 0 | Q12874 | Splicing factor 3A subunit 3 OS=Homo sapiens OX=9606 GN=SF3A3 PE=1 SV=1 | 3 | 2 | 2 | 2 | 501 | 58.8 | 4.35 | |
| 0 | Q9HCN8 | Stromal cell-derived factor 2-like protein 1 OS=Homo sapiens OX=9606 GN=SDF2L1 PE=1 SV=2 | 9 | 1 | 1 | 1 | 221 | 23.6 | 4.35 | 28298.69531 |
| 0 | Q13242 | Serine/arginine-rich splicing factor 9 OS=Homo sapiens OX=9606 GN=SRSF9 PE=1 SV=1 | 7 | 2 | 2 | 1 | 221 | 25.5 | 4.33 | 17324.97656 |
| 0 | P19388 | DNA-directed RNA polymerases I, II, and III subunit RPABC1 OS=Homo sapiens OX=9606 GN=POLR2E PE=1 SV=4 | 7 | 1 | 2 | 1 | 210 | 24.5 | 4.3 | 15471.46777 |
| 0 | Q8WWM7 | Ataxin-2-like protein OS=Homo sapiens OX=9606 GN=ATXN2L PE=1 SV=2 | 4 | 2 | 2 | 2 | 1075 | 113.3 | 4.27 | 25567.37012 |
| 0 | P62891 | 60S ribosomal protein L39 OS=Homo sapiens OX=9606 GN=RPL39 PE=1 SV=2 | 25 | 2 | 3 | 2 | 51 | 6.4 | 4.22 | 169967.918 |
| 0 | P41223 | Protein BUD31 homolog OS=Homo sapiens OX=9606 GN=BUD31 PE=1 SV=2 | 24 | 3 | 3 | 3 | 144 | 17 | 4.22 | 27725.33789 |
| 0 | Q9Y520 | Protein PRRC2C OS=Homo sapiens OX=9606 GN=PRRC2C PE=1 SV=4 | 1 | 1 | 1 | 1 | 2896 | 316.7 | 4.22 | |
| 0 | O43660 | Pleiotropic regulator 1 OS=Homo sapiens OX=9606 GN=PLRG1 PE=1 SV=1 | 5 | 1 | 1 | 1 | 514 | 57.2 | 4.2 | |
| 0 | P23246 | Splicing factor, proline- and glutamine-rich OS=Homo sapiens OX=9606 GN=SFPQ PE=1 SV=2 | 5 | 2 | 2 | 2 | 707 | 76.1 | 4.17 | 25277.38965 |
| 0 | O75531 | Barrier-to-autointegration factor OS=Homo sapiens OX=9606 GN=BANF1 PE=1 SV=1 | 43 | 2 | 2 | 2 | 89 | 10.1 | 4.12 | 16686.55859 |
| 0 | P62873 | Guanine nucleotide-binding protein G(I)/G(S)/G(T) subunit beta-1 OS=Homo sapiens OX=9606 GN=GNB1 PE=1 SV=3 | 6 | 1 | 1 | 1 | 340 | 37.4 | 4.11 | 19409.77344 |
| 0 | P46781 | 40S ribosomal protein S9 OS=Homo sapiens OX=9606 GN=RPS9 PE=1 SV=3 | 8 | 2 | 2 | 2 | 194 | 22.6 | 4.1 | 28877.67578 |
| 0 | Q9Y3C1 | Nucleolar protein 16 OS=Homo sapiens OX=9606 GN=NOP16 PE=1 SV=2 | 11 | 2 | 2 | 2 | 178 | 21.2 | 4.06 | 23889.45508 |
| 0 | P00492 | Hypoxanthine-guanine phosphoribosyltransferase OS=Homo sapiens OX=9606 GN=HPRT1 PE=1 SV=2 | 24 | 3 | 3 | 3 | 218 | 24.6 | 4.06 | 112441.7344 |
| 0 | Q9UIL1 | Short coiled-coil protein OS=Homo sapiens OX=9606 GN=SCOC PE=1 SV=2 | 18 | 1 | 1 | 1 | 159 | 18 | 4.01 | |
| 0 | Q9H9B4 | Sideroflexin-1 OS=Homo sapiens OX=9606 GN=SFXN1 PE=1 SV=4 | 4 | 1 | 1 | 1 | 322 | 35.6 | 4 | 14556.40625 |
| 0 | P49755 | Transmembrane emp24 domain-containing protein 10 OS=Homo sapiens OX=9606 GN=TMED10 PE=1 SV=2 | 12 | 2 | 2 | 2 | 219 | 25 | 3.99 | 52287.59766 |
| 0 | Q86TB3 | Alpha-protein kinase 2 OS=Homo sapiens OX=9606 GN=ALPK2 PE=1 SV=3 | 2 | 2 | 2 | 2 | 2170 | 236.9 | 3.98 | 14596.77246 |
| 0 | Q99848 | Probable rRNA-processing protein EBP2 OS=Homo sapiens OX=9606 GN=EBNA1BP2 PE=1 SV=2 | 6 | 2 | 2 | 2 | 306 | 34.8 | 3.95 | 19586.31055 |
| 0 | Q66PJ3 | ADP-ribosylation factor-like protein 6-interacting protein 4 OS=Homo sapiens OX=9606 GN=ARL6IP4 PE=1 SV=2 | 4 | 1 | 1 | 1 | 421 | 44.9 | 3.94 | |
| 0 | Q9UII2 | ATPase inhibitor, mitochondrial OS=Homo sapiens OX=9606 GN=ATP5IF1 PE=1 SV=1 | 17 | 2 | 2 | 2 | 106 | 12.2 | 3.89 | 129543.6563 |
| 0 | Q9UM54 | Unconventional myosin-VI OS=Homo sapiens OX=9606 GN=MYO6 PE=1 SV=4 | 1 | 1 | 1 | 1 | 1294 | 149.6 | 3.88 | |
| 0 | P62310 | U6 snRNA-associated Sm-like protein LSm3 OS=Homo sapiens OX=9606 GN=LSM3 PE=1 SV=2 | 22 | 2 | 2 | 2 | 102 | 11.8 | 3.86 | 34685.33691 |
| 0 | P49368 | T-complex protein 1 subunit gamma OS=Homo sapiens OX=9606 GN=CCT3 PE=1 SV=4 | 3 | 1 | 1 | 1 | 545 | 60.5 | 3.84 | 23802.17383 |

| | | | | | | | | | | |
|---|---|---|---|---|---|---|---|---|---|---|
| 0 | P23526 | Adenosylhomocysteinase OS=Homo sapiens OX=9606 GN=AHCY PE=1 SV=4 | 3 | 1 | 1 | 1 | 432 | 47.7 | 3.83 | |
| 0 | Q5RKV6 | Exosome complex component MTR3 OS=Homo sapiens OX=9606 GN=EXOSC6 PE=1 SV=1 | 10 | 1 | 1 | 1 | 272 | 28.2 | 3.8 | 19498.01172 |
| 0 | Q9UJZ1 | Stomatin-like protein 2, mitochondrial OS=Homo sapiens OX=9606 GN=STOML2 PE=1 SV=1 | 9 | 1 | 1 | 1 | 356 | 38.5 | 3.76 | 4411.717773 |
| 0 | Q9UBS4 | DnaJ homolog subfamily B member 11 OS=Homo sapiens OX=9606 GN=DNAJB11 PE=1 SV=1 | 4 | 1 | 1 | 1 | 358 | 40.5 | 3.74 | 12116.55273 |
| 0 | O43504 | Ragulator complex protein LAMTOR5 OS=Homo sapiens OX=9606 GN=LAMTOR5 PE=1 SV=1 | 37 | 1 | 1 | 1 | 91 | 9.6 | 3.74 | |
| 0 | Q9Y6M1 | Insulin-like growth factor 2 mRNA-binding protein 2 OS=Homo sapiens OX=9606 GN=IGF2BP2 PE=1 SV=2 | 8 | 3 | 3 | 2 | 599 | 66.1 | 3.72 | 18747.4082 |
| 0 | Q15005 | Signal peptidase complex subunit 2 OS=Homo sapiens OX=9606 GN=SPCS2 PE=1 SV=3 | 10 | 1 | 1 | 1 | 226 | 25 | 3.71 | |
| 0 | O15260 | Surfeit locus protein 4 OS=Homo sapiens OX=9606 GN=SURF4 PE=1 SV=3 | 7 | 1 | 1 | 1 | 269 | 30.4 | 3.69 | 5543.247559 |
| 0 | Q9UNZ5 | Leydig cell tumor 10 kDa protein homolog OS=Homo sapiens OX=9606 GN=C19orf53 PE=1 SV=1 | 22 | 2 | 2 | 2 | 99 | 10.6 | 3.65 | 22572.73975 |
| 0 | Q96EY5 | Multivesicular body subunit 12A OS=Homo sapiens OX=9606 GN=MVB12A PE=1 SV=1 | 8 | 1 | 1 | 1 | 273 | 28.8 | 3.64 | |
| 0 | P61927 | 60S ribosomal protein L37 OS=Homo sapiens OX=9606 GN=RPL37 PE=1 SV=2 | 19 | 2 | 3 | 2 | 97 | 11.1 | 3.64 | 75653.72266 |
| 0 | P13639 | Elongation factor 2 OS=Homo sapiens OX=9606 GN=EEF2 PE=1 SV=4 | 2 | 1 | 1 | 1 | 858 | 95.3 | 3.6 | |
| 0 | Q02978 | Mitochondrial 2-oxoglutarate/malate carrier protein OS=Homo sapiens OX=9606 GN=SLC25A11 PE=1 SV=3 | 5 | 1 | 1 | 1 | 314 | 34 | 3.59 | 45230.93359 |
| 0 | P29558 | RNA-binding motif, single-stranded-interacting protein 1 OS=Homo sapiens OX=9606 GN=RBMS1 PE=1 SV=3 | 4 | 1 | 1 | 1 | 406 | 44.5 | 3.58 | 15481.56641 |
| 0 | O15160 | DNA-directed RNA polymerases I and III subunit RPAC1 OS=Homo sapiens OX=9606 GN=POLR1C PE=1 SV=1 | 6 | 1 | 1 | 1 | 346 | 39.2 | 3.57 | 16074.89063 |
| 0 | O00193 | Small acidic protein OS=Homo sapiens OX=9606 GN=SMAP PE=1 SV=1 | 13 | 1 | 1 | 1 | 183 | 20.3 | 3.55 | |
| 0 | Q9Y3B4 | Splicing factor 3B subunit 6 OS=Homo sapiens OX=9606 GN=SF3B6 PE=1 SV=1 | 14 | 1 | 1 | 1 | 125 | 14.6 | 3.52 | 18184.85352 |
| 0 | Q92945 | Far upstream element-binding protein 2 OS=Homo sapiens OX=9606 GN=KHSRP PE=1 SV=4 | 3 | 1 | 1 | 1 | 711 | 73.1 | 3.49 | |
| 0 | P49458 | Signal recognition particle 9 kDa protein OS=Homo sapiens OX=9606 GN=SRP9 PE=1 SV=2 | 14 | 1 | 1 | 1 | 86 | 10.1 | 3.49 | 155182.9219 |
| 0 | Q9UK41 | Vacuolar protein sorting-associated protein 28 homolog OS=Homo sapiens OX=9606 GN=VPS28 PE=1 SV=1 | 13 | 1 | 2 | 1 | 221 | 25.4 | 3.48 | |
| 0 | P30048 | Thioredoxin-dependent peroxide reductase, mitochondrial OS=Homo sapiens OX=9606 GN=PRDX3 PE=1 SV=3 | 5 | 1 | 1 | 1 | 256 | 27.7 | 3.42 | 14954.74414 |
| 0 | Q9C037 | E3 ubiquitin-protein ligase TRIM4 OS=Homo sapiens OX=9606 GN=TRIM4 PE=1 SV=2 | 3 | 1 | 1 | 1 | 500 | 57.4 | 3.41 | 9762.773438 |
| 0 | Q9NX24 | H/ACA ribonucleoprotein complex subunit 2 OS=Homo sapiens OX=9606 GN=NHP2 PE=1 SV=1 | 12 | 1 | 1 | 1 | 153 | 17.2 | 3.36 | 23473.70313 |
| 0 | P14854 | Cytochrome c oxidase subunit 6B1 OS=Homo sapiens OX=9606 GN=COX6B1 PE=1 SV=2 | 13 | 1 | 1 | 1 | 86 | 10.2 | 3.31 | 15871.84961 |
| 0 | Q92908 | Transcription factor GATA-6 OS=Homo sapiens OX=9606 GN=GATA6 PE=1 SV=2 | 5 | 1 | 1 | 1 | 595 | 60 | 3.3 | |
| 0 | P33992 | DNA replication licensing factor MCM5 OS=Homo sapiens OX=9606 GN=MCM5 PE=1 SV=5 | 2 | 1 | 1 | 1 | 734 | 82.2 | 3.28 | 22462.98242 |
| 0 | Q8TF09 | Dynein light chain roadblock-type 2 OS=Homo sapiens OX=9606 GN=DYNLRB2 PE=1 SV=1 | 17 | 1 | 1 | 1 | 96 | 10.8 | 3.26 | 14730.41309 |
| 0 | O43852 | Calumenin OS=Homo sapiens OX=9606 GN=CALU PE=1 SV=2 | 4 | 1 | 1 | 1 | 315 | 37.1 | 3.26 | |
| 0 | O96019 | Actin-like protein 6A OS=Homo sapiens OX=9606 GN=ACTL6A PE=1 SV=1 | 7 | 1 | 1 | 1 | 429 | 47.4 | 3.25 | |
| 0 | Q8NAV1 | Pre-mRNA-splicing factor 38A OS=Homo sapiens OX=9606 GN=PRPF38A PE=1 SV=1 | 5 | 1 | 1 | 1 | 312 | 37.5 | 3.23 | 19886.86719 |
| 0 | Q7L5D6 | Golgi to ER traffic protein 4 homolog OS=Homo sapiens OX=9606 GN=GET4 PE=1 SV=1 | 4 | 1 | 1 | 1 | 327 | 36.5 | 3.21 | 70047.03906 |
| 0 | P20645 | Cation-dependent mannose-6-phosphate receptor OS=Homo sapiens OX=9606 GN=M6PR PE=1 SV=1 | 5 | 1 | 1 | 1 | 277 | 31 | 3.21 | 19151.73438 |
| 0 | Q9BRT6 | Protein LLP homolog OS=Homo sapiens OX=9606 GN=LLPH PE=1 SV=1 | 12 | 1 | 1 | 1 | 129 | 15.2 | 3.2 | 18776.38477 |
| 0 | P08579 | U2 small nuclear ribonucleoprotein B'' OS=Homo sapiens OX=9606 GN=SNRPB2 PE=1 SV=1 | 8 | 1 | 1 | 1 | 225 | 25.5 | 3.16 | |
| 0 | P00387 | NADH-cytochrome b5 reductase 3 OS=Homo sapiens OX=9606 GN=CYB5R3 PE=1 SV=3 | 6 | 1 | 1 | 1 | 301 | 34.2 | 3.14 | 14106.44727 |
| 0 | O15234 | Protein CASC3 OS=Homo sapiens OX=9606 GN=CASC3 PE=1 SV=2 | 2 | 1 | 1 | 1 | 703 | 76.2 | 3.08 | 22736.59766 |
| 0 | P21108 | Ribose-phosphate pyrophosphokinase 3 OS=Homo sapiens OX=9606 GN=PRPS1L1 PE=1 SV=2 | 4 | 1 | 1 | 1 | 318 | 34.8 | 3.07 | |
| 0 | P42285 | Exosome RNA helicase MTR4 OS=Homo sapiens OX=9606 GN=MTREX PE=1 SV=3 | 2 | 1 | 1 | 1 | 1042 | 117.7 | 3.07 | |
| 0 | P02549 | Spectrin alpha chain, erythrocytic 1 OS=Homo sapiens OX=9606 GN=SPTA1 PE=1 SV=5 | 0 | 1 | 1 | 1 | 2419 | 279.8 | 3.03 | 26339.08203 |
| 0 | Q16342 | Programmed cell death protein 2 OS=Homo sapiens OX=9606 GN=PDCD2 PE=1 SV=2 | 6 | 1 | 1 | 1 | 344 | 38.6 | 3.02 | 27792.94922 |
| 0 | P22695 | Cytochrome b-c1 complex subunit 2, mitochondrial OS=Homo sapiens OX=9606 GN=UQCRC2 PE=1 SV=3 | 4 | 1 | 1 | 1 | 453 | 48.4 | 3.01 | |
| 0 | Q9UBL3 | Set1/Ash2 histone methyltransferase complex subunit ASH2 OS=Homo sapiens OX=9606 GN=ASH2L PE=1 SV=1 | 4 | 1 | 1 | 1 | 628 | 68.7 | 3 | 14546.33496 |
| 0 | P04844 | Dolichyl-diphosphooligosaccharide--protein glycosyltransferase subunit 2 OS=Homo sapiens OX=9606 GN=RPN2 PE=1 SV=3 | 10 | 3 | 3 | 3 | 631 | 69.2 | 2.98 | 6256.616699 |
| 0 | P48643 | T-complex protein 1 subunit epsilon OS=Homo sapiens OX=9606 GN=CCT5 PE=1 SV=1 | 3 | 1 | 1 | 1 | 541 | 59.6 | 2.93 | 10866.77148 |
| 0 | O94906 | Pre-mRNA-processing factor 6 OS=Homo sapiens OX=9606 GN=PRPF6 PE=1 SV=1 | 1 | 1 | 1 | 1 | 941 | 106.9 | 2.92 | |
| 0 | P06454 | Prothymosin alpha OS=Homo sapiens OX=9606 GN=PTMA PE=1 SV=2 | 14 | 1 | 1 | 1 | 111 | 12.2 | 2.92 | 17705.37891 |
| 0 | Q6N021 | Methylcytosine dioxygenase TET2 OS=Homo sapiens OX=9606 GN=TET2 PE=1 SV=3 | 1 | 1 | 1 | 1 | 2002 | 223.7 | 2.91 | |
| 0 | Q14739 | Delta(14)-sterol reductase LBR OS=Homo sapiens OX=9606 GN=LBR PE=1 SV=2 | 2 | 1 | 1 | 1 | 615 | 70.7 | 2.89 | |
| 0 | P62857 | 40S ribosomal protein S28 OS=Homo sapiens OX=9606 GN=RPS28 PE=1 SV=1 | 17 | 1 | 1 | 1 | 69 | 7.8 | 2.87 | 23506.88477 |
| 0 | P25205 | DNA replication licensing factor MCM3 OS=Homo sapiens OX=9606 GN=MCM3 PE=1 SV=3 | 4 | 1 | 1 | 1 | 808 | 90.9 | 2.86 | 8362.578125 |
| 0 | P28799 | Progranulin OS=Homo sapiens OX=9606 GN=GRN PE=1 SV=2 | 3 | 1 | 1 | 1 | 593 | 63.5 | 2.85 | 18351.24414 |
| 0 | O95071 | E3 ubiquitin-protein ligase UBR5 OS=Homo sapiens OX=9606 GN=UBR5 PE=1 SV=2 | 1 | 1 | 1 | 1 | 2799 | 309.2 | 2.85 | |
| 0 | Q9NPL8 | Complex I assembly factor TIMMDC1, mitochondrial OS=Homo sapiens OX=9606 GN=TIMMDC1 PE=1 SV=2 | 9 | 1 | 1 | 1 | 285 | 32.2 | 2.84 | |
| 0 | Q9NPE3 | H/ACA ribonucleoprotein complex subunit 3 OS=Homo sapiens OX=9606 GN=NOP10 PE=1 SV=1 | 20 | 1 | 1 | 1 | 64 | 7.7 | 2.83 | 36170.66797 |
| 0 | P17480 | Nucleolar transcription factor 1 OS=Homo sapiens OX=9606 GN=UBTF PE=1 SV=1 | 2 | 1 | 1 | 1 | 764 | 89.4 | 2.83 | |
| 0 | Q15758 | Neutral amino acid transporter B(0) OS=Homo sapiens OX=9606 GN=SLC1A5 PE=1 SV=2 | 4 | 1 | 1 | 1 | 541 | 56.6 | 2.83 | 26491.1543 |
| 0 | Q9BUT9 | MAPK regulated corepressor interacting protein 2 OS=Homo sapiens OX=9606 GN=MCRIP2 PE=1 SV=2 | 19 | 1 | 1 | 1 | 160 | 17.8 | 2.78 | |
| 0 | Q8IYI6 | Exocyst complex component 8 OS=Homo sapiens OX=9606 GN=EXOC8 PE=1 SV=2 | 2 | 1 | 1 | 1 | 725 | 81.7 | 2.77 | |
| 0 | P0DN79 | Cystathionine beta-synthase-like protein OS=Homo sapiens OX=9606 GN=CBSL PE=1 SV=1 | 2 | 1 | 1 | 1 | 551 | 60.5 | 2.77 | 12520.10742 |
| 0 | P31151 | Protein S100-A7 OS=Homo sapiens OX=9606 GN=S100A7 PE=1 SV=4 | 11 | 1 | 1 | 1 | 101 | 11.5 | 2.77 | 17197.38086 |
| 0 | P49773 | Histidine triad nucleotide-binding protein 1 OS=Homo sapiens OX=9606 GN=HINT1 PE=1 SV=2 | 11 | 1 | 1 | 1 | 126 | 13.8 | 2.72 | 49326.28516 |
| 0 | Q13561 | Dynactin subunit 2 OS=Homo sapiens OX=9606 GN=DCTN2 PE=1 SV=4 | 3 | 1 | 1 | 1 | 401 | 44.2 | 2.71 | 15519.89355 |

| | | | | | | | | | | |
|---|---|---|---|---|---|---|---|---|---|---|
| 0 | Q9BQA1 | Methylosome protein 50 OS=Homo sapiens OX=9606 GN=WDR77 PE=1 SV=1 | 5 | 1 | 1 | 1 | 342 | 36.7 | 2.69 | 17484.86914 |
| 0 | Q9Y5M8 | Signal recognition particle receptor subunit beta OS=Homo sapiens OX=9606 GN=SRPRB PE=1 SV=3 | 7 | 1 | 1 | 1 | 271 | 29.7 | 2.69 | 11855.69141 |
| 0 | Q15007 | Pre-mRNA-splicing regulator WTAP OS=Homo sapiens OX=9606 GN=WTAP PE=1 SV=2 | 4 | 1 | 1 | 1 | 396 | 44.2 | 2.66 | 19062.83789 |
| 0 | Q7RTR0 | NACHT, LRR and PYD domains-containing protein 9 OS=Homo sapiens OX=9606 GN=NLRP9 PE=1 SV=1 | 1 | 1 | 1 | 1 | 991 | 113.2 | 2.65 | |
| 0 | Q9NP66 | High mobility group protein 20A OS=Homo sapiens OX=9606 GN=HMG20A PE=1 SV=1 | 5 | 1 | 1 | 1 | 347 | 40.1 | 2.65 | |
| 0 | P46060 | Ran GTPase-activating protein 1 OS=Homo sapiens OX=9606 GN=RANGAP1 PE=1 SV=1 | 3 | 1 | 1 | 1 | 587 | 63.5 | 2.63 | 9362.350586 |
| 0 | Q92560 | Ubiquitin carboxyl-terminal hydrolase BAP1 OS=Homo sapiens OX=9606 GN=BAP1 PE=1 SV=2 | 2 | 1 | 1 | 1 | 729 | 80.3 | 2.63 | 19188.70313 |
| 0 | Q9NPF4 | Probable tRNA N6-adenosine threonylcarbamoyltransferase OS=Homo sapiens OX=9606 GN=OSGEP PE=1 SV=1 | 4 | 1 | 1 | 1 | 335 | 36.4 | 2.61 | 11204.16309 |
| 0 | P62942 | Peptidyl-prolyl cis-trans isomerase FKBP1A OS=Homo sapiens OX=9606 GN=FKBP1A PE=1 SV=2 | 13 | 1 | 1 | 1 | 108 | 11.9 | 2.61 | 19356.07031 |
| 0 | P60468 | Protein transport protein Sec61 subunit beta OS=Homo sapiens OX=9606 GN=SEC61B PE=1 SV=2 | 17 | 1 | 1 | 1 | 96 | 10 | 2.6 | 23964.82422 |
| 0 | P62714 | Serine/threonine-protein phosphatase 2A catalytic subunit beta isoform OS=Homo sapiens OX=9606 GN=PPP2CB PE=1 SV=1 | 8 | 1 | 1 | 1 | 309 | 35.6 | 2.6 | |
| 0 | P00338 | L-lactate dehydrogenase A chain OS=Homo sapiens OX=9606 GN=LDHA PE=1 SV=2 | 4 | 1 | 1 | 1 | 332 | 36.7 | 2.6 | |
| 0 | Q99496 | E3 ubiquitin-protein ligase RING2 OS=Homo sapiens OX=9606 GN=RNF2 PE=1 SV=1 | 4 | 1 | 1 | 1 | 336 | 37.6 | 2.6 | |
| 0 | Q2KHR3 | Glutamine and serine-rich protein 1 OS=Homo sapiens OX=9606 GN=QSER1 PE=1 SV=3 | 1 | 1 | 1 | 1 | 1735 | 189.9 | 2.58 | 9938.038086 |
| 0 | Q13547 | Histone deacetylase 1 OS=Homo sapiens OX=9606 GN=HDAC1 PE=1 SV=1 | 2 | 1 | 1 | 1 | 482 | 55.1 | 2.57 | 27986.18945 |
| 0 | O75348 | V-type proton ATPase subunit G 1 OS=Homo sapiens OX=9606 GN=ATP6V1G1 PE=1 SV=3 | 12 | 1 | 1 | 1 | 118 | 13.7 | 2.57 | |
| 0 | P78406 | mRNA export factor OS=Homo sapiens OX=9606 GN=RAE1 PE=1 SV=1 | 2 | 1 | 1 | 1 | 368 | 40.9 | 2.54 | 20885.07031 |
| 0 | P49756 | RNA-binding protein 25 OS=Homo sapiens OX=9606 GN=RBM25 PE=1 SV=3 | 2 | 1 | 1 | 1 | 843 | 100.1 | 2.54 | |
| 0 | Q96ND8 | Zinc finger protein 583 OS=Homo sapiens OX=9606 GN=ZNF583 PE=2 SV=2 | 3 | 1 | 1 | 1 | 569 | 66 | 2.53 | 19718.0957 |
| 0 | Q8N573 | Oxidation resistance protein 1 OS=Homo sapiens OX=9606 GN=OXR1 PE=1 SV=2 | 2 | 1 | 1 | 1 | 874 | 97.9 | 2.53 | |
| 0 | Q9UGM3 | Deleted in malignant brain tumors 1 protein OS=Homo sapiens OX=9606 GN=DMBT1 PE=1 SV=2 | 7 | 1 | 1 | 1 | 2413 | 260.6 | 2.51 | 46415.53906 |
| 0 | P61626 | Lysozyme C OS=Homo sapiens OX=9606 GN=LYZ PE=1 SV=1 | 7 | 2 | 2 | 2 | 148 | 16.5 | 2.51 | 13489.24902 |
| 0 | O75190 | DnaJ homolog subfamily B member 6 OS=Homo sapiens OX=9606 GN=DNAJB6 PE=1 SV=2 | 3 | 1 | 1 | 1 | 326 | 36.1 | 2.51 | 14830.27637 |
| 0 | O60925 | Prefoldin subunit 1 OS=Homo sapiens OX=9606 GN=PFDN1 PE=1 SV=2 | 9 | 1 | 1 | 1 | 122 | 14.2 | 2.51 | 27118.89258 |
| 0 | Q09666 | Neuroblast differentiation-associated protein AHNAK OS=Homo sapiens OX=9606 GN=AHNAK PE=1 SV=2 | 1 | 1 | 1 | 1 | 5890 | 628.7 | 2.5 | 16044.2627 |
| 0 | P48378 | DNA-binding protein RFX2 OS=Homo sapiens OX=9606 GN=RFX2 PE=1 SV=2 | 2 | 1 | 1 | 1 | 723 | 79.9 | 2.5 | 88117.33594 |
| 0 | Q9BRJ7 | Tudor-interacting repair regulator protein OS=Homo sapiens OX=9606 GN=NUDT16L1 PE=1 SV=1 | 8 | 1 | 1 | 1 | 211 | 23.3 | 2.5 | 11321.75684 |
| 0 | Q9BT78 | COP9 signalosome complex subunit 4 OS=Homo sapiens OX=9606 GN=COPS4 PE=1 SV=1 | 2 | 1 | 1 | 1 | 406 | 46.2 | 2.49 | 12857.16309 |
| 0 | P61163 | Alpha-centractin OS=Homo sapiens OX=9606 GN=ACTR1A PE=1 SV=1 | 3 | 1 | 1 | 1 | 376 | 42.6 | 2.48 | 32052.57422 |
| 0 | P07919 | Cytochrome b-c1 complex subunit 6, mitochondrial OS=Homo sapiens OX=9606 GN=UQCRH PE=1 SV=2 | 10 | 1 | 1 | 1 | 91 | 10.7 | 2.45 | 24744.17383 |
| 0 | Q06587 | E3 ubiquitin-protein ligase RING1 OS=Homo sapiens OX=9606 GN=RING1 PE=1 SV=2 | 3 | 1 | 1 | 1 | 406 | 42.4 | 2.45 | 12336.98633 |
| 0 | Q14244 | Ensconsin OS=Homo sapiens OX=9606 GN=MAP7 PE=1 SV=1 | 1 | 1 | 1 | 1 | 749 | 84 | 2.42 | 109842.4375 |
| 0 | Q96DI7 | U5 small nuclear ribonucleoprotein 40 kDa protein OS=Homo sapiens OX=9606 GN=SNRNP40 PE=1 SV=1 | 3 | 1 | 1 | 1 | 357 | 39.3 | 2.41 | 27103.36133 |
| 0 | Q14684 | Ribosomal RNA processing protein 1 homolog B OS=Homo sapiens OX=9606 GN=RRP1B PE=1 SV=3 | 4 | 1 | 1 | 1 | 758 | 84.4 | 2.4 | 11522.35059 |
| 0 | P52565 | Rho GDP-dissociation inhibitor 1 OS=Homo sapiens OX=9606 GN=ARHGDIA PE=1 SV=3 | 16 | 1 | 1 | 1 | 204 | 23.2 | 2.4 | 13888.12305 |
| 0 | P53396 | ATP-citrate synthase OS=Homo sapiens OX=9606 GN=ACLY PE=1 SV=3 | 1 | 1 | 1 | 1 | 1101 | 120.8 | 2.39 | 14504.4082 |
| 0 | Q9NQC3 | Reticulon-4 OS=Homo sapiens OX=9606 GN=RTN4 PE=1 SV=2 | 3 | 1 | 1 | 1 | 1192 | 129.9 | 2.38 | 21494.03711 |
| 0 | P49006 | MARCKS-related protein OS=Homo sapiens OX=9606 GN=MARCKSL1 PE=1 SV=2 | 4 | 1 | 1 | 1 | 195 | 19.5 | 2.38 | 29465.19922 |
| 0 | P40429 | 60S ribosomal protein L13a OS=Homo sapiens OX=9606 GN=RPL13A PE=1 SV=2 | 5 | 1 | 1 | 1 | 203 | 23.6 | 2.38 | 18337.76172 |
| 0 | O75475 | PC4 and SFRS1-interacting protein OS=Homo sapiens OX=9606 GN=PSIP1 PE=1 SV=1 | 5 | 2 | 2 | 2 | 530 | 60.1 | 2.38 | 43674.53125 |
| 0 | Q9GZZ1 | N-alpha-acetyltransferase 50 OS=Homo sapiens OX=9606 GN=NAA50 PE=1 SV=1 | 10 | 1 | 1 | 1 | 169 | 19.4 | 2.35 | |
| 0 | Q8TA86 | Retinitis pigmentosa 9 protein OS=Homo sapiens OX=9606 GN=RP9 PE=1 SV=2 | 7 | 1 | 1 | 1 | 221 | 26.1 | 2.35 | 27602.12305 |
| 0 | P62995 | Transformer-2 protein homolog beta OS=Homo sapiens OX=9606 GN=TRA2B PE=1 SV=1 | 3 | 1 | 1 | 1 | 288 | 33.6 | 2.31 | 77513.11719 |
| 0 | Q9P258 | Protein RCC2 OS=Homo sapiens OX=9606 GN=RCC2 PE=1 SV=2 | 3 | 1 | 1 | 1 | 522 | 56 | 2.3 | 11181.06934 |
| 0 | P61962 | DDB1- and CUL4-associated factor 7 OS=Homo sapiens OX=9606 GN=DCAF7 PE=1 SV=1 | 4 | 1 | 1 | 1 | 342 | 38.9 | 2.3 | 9351.013672 |
| 0 | Q9GZT3 | SRA stem-loop-interacting RNA-binding protein, mitochondrial OS=Homo sapiens OX=9606 GN=SLIRP PE=1 SV=1 | 11 | 1 | 1 | 1 | 109 | 12.3 | 2.3 | 10955.91406 |
| 0 | P48047 | ATP synthase subunit O, mitochondrial OS=Homo sapiens OX=9606 GN=ATP5PO PE=1 SV=1 | 11 | 2 | 2 | 2 | 213 | 23.3 | 2.28 | 42282.07715 |
| 0 | O15145 | Actin-related protein 2/3 complex subunit 3 OS=Homo sapiens OX=9606 GN=ARPC3 PE=1 SV=3 | 6 | 1 | 1 | 1 | 178 | 20.5 | 2.28 | 11857.52441 |
| 0 | Q9Y608 | Leucine-rich repeat flightless-interacting protein 2 OS=Homo sapiens OX=9606 GN=LRRFIP2 PE=1 SV=1 | 2 | 1 | 1 | 1 | 721 | 82.1 | 2.27 | 17973.43164 |
| 0 | Q6P2Q9 | Pre-mRNA-processing-splicing factor 8 OS=Homo sapiens OX=9606 GN=PRPF8 PE=1 SV=2 | 1 | 1 | 1 | 1 | 2335 | 273.4 | 2.27 | 21212.61328 |
| 0 | P21802 | Fibroblast growth factor receptor 2 OS=Homo sapiens OX=9606 GN=FGFR2 PE=1 SV=1 | 2 | 1 | 1 | 1 | 821 | 92 | 2.25 | 35914.44531 |
| 0 | Q9Y3B7 | 39S ribosomal protein L11, mitochondrial OS=Homo sapiens OX=9606 GN=MRPL11 PE=1 SV=1 | 11 | 1 | 1 | 1 | 192 | 20.7 | 2.25 | |
| 0 | Q9ULA0 | Aspartyl aminopeptidase OS=Homo sapiens OX=9606 GN=DNPEP PE=1 SV=2 | 6 | 2 | 2 | 2 | 485 | 53.4 | 2.24 | 28725.6792 |
| 0 | P24539 | ATP synthase F(0) complex subunit B1, mitochondrial OS=Homo sapiens OX=9606 GN=ATP5PB PE=1 SV=2 | 4 | 1 | 1 | 1 | 256 | 28.9 | 2.24 | |
| 0 | O95816 | BAG family molecular chaperone regulator 2 OS=Homo sapiens OX=9606 GN=BAG2 PE=1 SV=1 | 5 | 1 | 1 | 1 | 211 | 23.8 | 2.24 | |
| 0 | D6REC4 | Cilia- and flagella-associated protein 99 OS=Homo sapiens OX=9606 GN=CFAP99 PE=3 SV=1 | 2 | 1 | 1 | 1 | 459 | 52.3 | 2.23 | 73775.125 |
| 0 | O14904 | Protein Wnt-9a OS=Homo sapiens OX=9606 GN=WNT9A PE=1 SV=2 | 5 | 1 | 1 | 1 | 365 | 40.3 | 2.22 | 27935.5625 |
| 0 | Q9Y2W2 | WW domain-binding protein 11 OS=Homo sapiens OX=9606 GN=WBP11 PE=1 SV=1 | 4 | 1 | 1 | 1 | 641 | 70 | 2.2 | |
| 0 | Q15369 | Elongin-C OS=Homo sapiens OX=9606 GN=ELOC PE=1 SV=1 | 12 | 1 | 1 | 1 | 112 | 12.5 | 2.2 | 35156.07813 |
| 0 | Q9ULD5 | Zinc finger protein 777 OS=Homo sapiens OX=9606 GN=ZNF777 PE=1 SV=3 | 1 | 1 | 1 | 1 | 831 | 93.7 | 2.19 | 29753.6582 |
| 0 | Q9HB58 | Sp110 nuclear body protein OS=Homo sapiens OX=9606 GN=SP110 PE=1 SV=5 | 1 | 1 | 1 | 1 | 689 | 78.3 | 2.16 | 39490.66406 |
| 0 | P10606 | Cytochrome c oxidase subunit 5B, mitochondrial OS=Homo sapiens OX=9606 GN=COX5B PE=1 SV=2 | 9 | 1 | 1 | 1 | 129 | 13.7 | 2.14 | 18848.74023 |

| | | | | | | | | | | |
|---|---|---|---|---|---|---|---|---|---|---|
| 0 | O43143 | Pre-mRNA-splicing factor ATP-dependent RNA helicase DHX15 OS=Homo sapiens OX=9606 GN=DHX15 PE=1 SV=2 | 1 | 1 | 1 | 1 | 795 | 90.9 | 2.12 | 9258.073242 |
| 0 | Q9NWU2 | Glucose-induced degradation protein 8 homolog OS=Homo sapiens OX=9606 GN=GID8 PE=1 SV=1 | 11 | 1 | 1 | 1 | 228 | 26.7 | 2.11 | |
| 0 | P54709 | Sodium/potassium-transporting ATPase subunit beta-3 OS=Homo sapiens OX=9606 GN=ATP1B3 PE=1 SV=1 | 4 | 1 | 1 | 1 | 279 | 31.5 | 2.11 | 35797.80469 |
| 0 | Q9NXG2 | THUMP domain-containing protein 1 OS=Homo sapiens OX=9606 GN=THUMPD1 PE=1 SV=2 | 5 | 1 | 1 | 1 | 353 | 39.3 | 2.11 | |
| 0 | O95782 | AP-2 complex subunit alpha-1 OS=Homo sapiens OX=9606 GN=AP2A1 PE=1 SV=3 | 1 | 1 | 1 | 1 | 977 | 107.5 | 2.1 | |
| 0 | Q9Y666 | Solute carrier family 12 member 7 OS=Homo sapiens OX=9606 GN=SLC12A7 PE=1 SV=3 | 3 | 1 | 1 | 1 | 1083 | 119 | 2.07 | 16670.86328 |
| 0 | O15126 | Secretory carrier-associated membrane protein 1 OS=Homo sapiens OX=9606 GN=SCAMP1 PE=1 SV=2 | 16 | 2 | 2 | 2 | 338 | 37.9 | 2.07 | 7456.182617 |
| 0 | P09110 | 3-ketoacyl-CoA thiolase, peroxisomal OS=Homo sapiens OX=9606 GN=ACAA1 PE=1 SV=2 | 2 | 1 | 1 | 1 | 424 | 44.3 | 2.05 | 14749.22461 |
| 0 | Q6XD76 | Achaete-scute homolog 4 OS=Homo sapiens OX=9606 GN=ASCL4 PE=1 SV=1 | 7 | 1 | 1 | 1 | 172 | 19.2 | 2.05 | 13496.80469 |
| 0 | O75947 | ATP synthase subunit d, mitochondrial OS=Homo sapiens OX=9606 GN=ATP5PD PE=1 SV=3 | 11 | 1 | 2 | 1 | 161 | 18.5 | 2.05 | 24282.29688 |
| 0 | Q15058 | Kinesin-like protein KIF14 OS=Homo sapiens OX=9606 GN=KIF14 PE=1 SV=1 | 1 | 1 | 1 | 1 | 1648 | 186.4 | 2.05 | 174100.875 |
| 0 | P34932 | Heat shock 70 kDa protein 4 OS=Homo sapiens OX=9606 GN=HSPA4 PE=1 SV=4 | 2 | 1 | 1 | 1 | 840 | 94.3 | 2.03 | |
| 0 | Q8NCA5 | Protein FAM98A OS=Homo sapiens OX=9606 GN=FAM98A PE=1 SV=2 | 2 | 1 | 1 | 1 | 518 | 55.2 | 2 | 40568.88672 |
| 0 | Q8WUW1 | Protein BRICK1 OS=Homo sapiens OX=9606 GN=BRK1 PE=1 SV=1 | 11 | 1 | 1 | 1 | 75 | 8.7 | 1.96 | 12875.19043 |
| 0 | Q9NW07 | Zinc finger protein 358 OS=Homo sapiens OX=9606 GN=ZNF358 PE=1 SV=2 | 2 | 1 | 1 | 1 | 568 | 59.3 | 1.95 | 10173.90137 |
| 0 | Q9BRP8 | Partner of Y14 and mago OS=Homo sapiens OX=9606 GN=PYM1 PE=1 SV=1 | 8 | 1 | 1 | 1 | 204 | 22.6 | 1.95 | 8939.061523 |
| 0 | Q07020 | 60S ribosomal protein L18 OS=Homo sapiens OX=9606 GN=RPL18 PE=1 SV=2 | 5 | 1 | 1 | 1 | 188 | 21.6 | 1.95 | 37674.04688 |
| 0 | Q6EEV6 | Small ubiquitin-related modifier 4 OS=Homo sapiens OX=9606 GN=SUMO4 PE=1 SV=2 | 13 | 1 | 1 | 1 | 95 | 10.7 | 1.94 | 22520.20703 |
| 0 | P06493 | Cyclin-dependent kinase 1 OS=Homo sapiens OX=9606 GN=CDK1 PE=1 SV=3 | 5 | 1 | 1 | 1 | 297 | 34.1 | 1.94 | |
| 0 | Q93074 | Mediator of RNA polymerase II transcription subunit 12 OS=Homo sapiens OX=9606 GN=MED12 PE=1 SV=4 | 0 | 1 | 1 | 1 | 2177 | 242.9 | 1.92 | 61981.33594 |
| 0 | Q13724 | Mannosyl-oligosaccharide glucosidase OS=Homo sapiens OX=9606 GN=MOGS PE=1 SV=5 | 10 | 4 | 4 | 4 | 837 | 91.9 | 1.9 | 69411.6543 |
| 0 | P42167 | Lamina-associated polypeptide 2, isoforms beta/gamma OS=Homo sapiens OX=9606 GN=TMPO PE=1 SV=2 | 3 | 1 | 1 | 1 | 454 | 50.6 | 1.89 | 20598.01172 |
| 0 | Q9HBD1 | Roquin-2 OS=Homo sapiens OX=9606 GN=RC3H2 PE=1 SV=2 | 1 | 1 | 1 | 1 | 1191 | 131.6 | 1.88 | 5095.147461 |
| 0 | Q8TE02 | Elongator complex protein 5 OS=Homo sapiens OX=9606 GN=ELP5 PE=1 SV=2 | 4 | 1 | 1 | 1 | 316 | 34.8 | 1.88 | 472103.7188 |
| 0 | O43760 | Synaptogyrin-2 OS=Homo sapiens OX=9606 GN=SYNGR2 PE=1 SV=1 | 4 | 1 | 1 | 1 | 224 | 24.8 | 1.85 | 105400.6875 |
| 0 | Q9ULC4 | Malignant T-cell-amplified sequence 1 OS=Homo sapiens OX=9606 GN=MCTS1 PE=1 SV=1 | 6 | 1 | 1 | 1 | 181 | 20.5 | 1.85 | |
| 0 | Q2TAY7 | WD40 repeat-containing protein SMU1 OS=Homo sapiens OX=9606 GN=SMU1 PE=1 SV=2 | 2 | 1 | 1 | 1 | 513 | 57.5 | 1.84 | 31088.05664 |
| 0 | O60264 | SWI/SNF-related matrix-associated actin-dependent regulator of chromatin subfamily A member 5 OS=Homo sapiens OX=9606 GN=SMARCA5 PE=1 SV=1 | 1 | 1 | 1 | 1 | 1052 | 121.8 | 1.84 | 13411.39551 |
| 0 | Q8IZT6 | Abnormal spindle-like microcephaly-associated protein OS=Homo sapiens OX=9606 GN=ASPM PE=1 SV=2 | 0 | 1 | 1 | 1 | 3477 | 409.5 | 1.83 | 43379.27344 |
| 0 | O14559 | Rho GTPase-activating protein 33 OS=Homo sapiens OX=9606 GN=ARHGAP33 PE=1 SV=2 | 1 | 1 | 1 | 1 | 1287 | 137.1 | 1.82 | 140967.3594 |
| 0 | O14964 | Hepatocyte growth factor-regulated tyrosine kinase substrate OS=Homo sapiens OX=9606 GN=HGS PE=1 SV=1 | 1 | 1 | 1 | 1 | 777 | 86.1 | 1.82 | 32618.62891 |
| 0 | P02765 | Alpha-2-HS-glycoprotein OS=Homo sapiens OX=9606 GN=AHSG PE=1 SV=2 | 2 | 1 | 1 | 1 | 367 | 39.3 | 1.81 | 12159.50293 |
| 0 | Q9ULJ1 | Protein BCAP OS=Homo sapiens OX=9606 GN=ODF2L PE=2 SV=2 | 2 | 1 | 1 | 1 | 636 | 73.7 | 1.8 | 34299.05859 |
| 0 | O75528 | Transcriptional adapter 3 OS=Homo sapiens OX=9606 GN=TADA3 PE=1 SV=1 | 3 | 1 | 1 | 1 | 432 | 48.9 | 1.8 | |
| 0 | Q96DN5 | TBC1 domain family member 31 OS=Homo sapiens OX=9606 GN=TBC1D31 PE=1 SV=2 | 1 | 1 | 1 | 1 | 1066 | 124.1 | 1.8 | 15853.64453 |
| 0 | Q8N309 | Leucine-rich repeat-containing protein 43 OS=Homo sapiens OX=9606 GN=LRRC43 PE=2 SV=2 | 2 | 1 | 1 | 1 | 656 | 73 | 1.78 | 36242.17578 |
| 0 | O75817 | Ribonuclease P protein subunit p20 OS=Homo sapiens OX=9606 GN=POP7 PE=1 SV=2 | 11 | 1 | 1 | 1 | 140 | 15.6 | 1.77 | 32755.36914 |
| 0 | Q9NVP1 | ATP-dependent RNA helicase DDX18 OS=Homo sapiens OX=9606 GN=DDX18 PE=1 SV=2 | 6 | 2 | 2 | 2 | 670 | 75.4 | 1.75 | 13608.88672 |
| 0 | P62491 | Ras-related protein Rab-11A OS=Homo sapiens OX=9606 GN=RAB11A PE=1 SV=3 | 6 | 1 | 1 | 1 | 216 | 24.4 | 1.75 | 16150.64551 |
| 0 | Q9Y4H2 | Insulin receptor substrate 2 OS=Homo sapiens OX=9606 GN=IRS2 PE=1 SV=2 | 1 | 1 | 1 | 1 | 1338 | 137.2 | 1.75 | |
| 0 | P47929 | Galectin-7 OS=Homo sapiens OX=9606 GN=LGALS7 PE=1 SV=2 | 8 | 1 | 1 | 1 | 136 | 15.1 | 1.74 | 14847.60547 |
| 0 | Q6N043 | Zinc finger protein 280D OS=Homo sapiens OX=9606 GN=ZNF280D PE=1 SV=3 | 1 | 1 | 1 | 1 | 979 | 109.2 | 1.73 | 78386.44531 |
| 0 | A6NC57 | Ankyrin repeat domain-containing protein 62 OS=Homo sapiens OX=9606 GN=ANKRD62 PE=2 SV=4 | 2 | 1 | 1 | 1 | 917 | 106.4 | 1.73 | 53702.89063 |
| 0 | Q9BZE4 | Nucleolar GTP-binding protein 1 OS=Homo sapiens OX=9606 GN=GTPBP4 PE=1 SV=3 | 2 | 1 | 1 | 1 | 634 | 73.9 | 1.73 | |
| 0 | Q8WY50 | Placenta-specific protein 4 OS=Homo sapiens OX=9606 GN=PLAC4 PE=2 SV=2 | 5 | 1 | 1 | 1 | 150 | 16.7 | 1.73 | 45307.61719 |
| 0 | O43148 | mRNA cap guanine-N7 methyltransferase OS=Homo sapiens OX=9606 GN=RNMT PE=1 SV=1 | 4 | 1 | 1 | 1 | 476 | 54.8 | 1.73 | |
| 0 | P02652 | Apolipoprotein A-II OS=Homo sapiens OX=9606 GN=APOA2 PE=1 SV=1 | 8 | 1 | 1 | 1 | 100 | 11.2 | 1.68 | 548552.125 |
| 0 | Q9UI47 | Catenin alpha-3 OS=Homo sapiens OX=9606 GN=CTNNA3 PE=1 SV=2 | 2 | 1 | 1 | 1 | 895 | 99.7 | 1.67 | 783421.25 |
| 0 | P40939 | Trifunctional enzyme subunit alpha, mitochondrial OS=Homo sapiens OX=9606 GN=HADHA PE=1 SV=2 | 2 | 1 | 1 | 1 | 763 | 82.9 | 1.66 | 31272.40625 |
| 0 | Q7Z478 | ATP-dependent RNA helicase DHX29 OS=Homo sapiens OX=9606 GN=DHX29 PE=1 SV=2 | 0 | 1 | 1 | 1 | 1369 | 155.1 | 1.66 | |
| 0 | Q9NZ01 | Very-long-chain enoyl-CoA reductase OS=Homo sapiens OX=9606 GN=TECR PE=1 SV=1 | 3 | 1 | 2 | 1 | 308 | 36 | 1.65 | 23923.89453 |
| 0 | P50991 | T-complex protein 1 subunit delta OS=Homo sapiens OX=9606 GN=CCT4 PE=1 SV=4 | 2 | 1 | 1 | 1 | 539 | 57.9 | 1.64 | |
| 0 | Q99961 | Endophilin-A2 OS=Homo sapiens OX=9606 GN=SH3GL1 PE=1 SV=1 | 3 | 1 | 1 | 1 | 368 | 41.5 | 1.63 | 37090.92578 |
| 0 | O60573 | Eukaryotic translation initiation factor 4E type 2 OS=Homo sapiens OX=9606 GN=EIF4E2 PE=1 SV=1 | 7 | 1 | 1 | 1 | 245 | 28.3 | 1.62 | |
| 0 | O95359 | Transforming acidic coiled-coil-containing protein 2 OS=Homo sapiens OX=9606 GN=TACC2 PE=1 SV=3 | 1 | 1 | 1 | 1 | 2948 | 309.2 | 0 | 18889.91797 |
| 0 | Q5JSZ5 | Protein PRRC2B OS=Homo sapiens OX=9606 GN=PRRC2B PE=1 SV=2 | 1 | 1 | 1 | 1 | 2229 | 242.8 | 0 | 36682.65234 |
| 0 | Q8N0Y7 | Probable phosphoglycerate mutase 4 OS=Homo sapiens OX=9606 GN=PGAM4 PE=3 SV=1 | 6 | 1 | 1 | 1 | 254 | 28.8 | 0 | 18203.95508 |
| 0 | Q6ZNJ1 | Neurobeachin-like protein 2 OS=Homo sapiens OX=9606 GN=NBEAL2 PE=1 SV=2 | 1 | 1 | 1 | 1 | 2754 | 302.3 | 0 | |
| 0 | Q16181 | Septin-7 OS=Homo sapiens OX=9606 GN=SEPTIN7 PE=1 SV=2 | 2 | 1 | 1 | 1 | 437 | 50.6 | 0 | 6676.350098 |
| 0 | Q9Y4Y9 | U6 snRNA-associated Sm-like protein LSm5 OS=Homo sapiens OX=9606 GN=LSM5 PE=1 SV=3 | 30 | 1 | 1 | 1 | 91 | 9.9 | 0 | 20466.37891 |
| 0 | P78367 | Homeobox protein Nkx-3.2 OS=Homo sapiens OX=9606 GN=NKX3-2 PE=1 SV=2 | 14 | 1 | 1 | 1 | 333 | 34.8 | 0 | 865292.75 |

| | | | | | | | | | | |
|---|---|---|---|---|---|---|---|---|---|---|
| 0 | P51790 | H(+)/Cl(-) exchange transporter 3 OS=Homo sapiens OX=9606 GN=CLCN3 PE=1 SV=2 | 1 | 1 | 1 | 1 | 818 | 90.9 | 0 | |
| 0 | O43157 | Plexin-B1 OS=Homo sapiens OX=9606 GN=PLXNB1 PE=1 SV=3 | 0 | 1 | 1 | 1 | 2135 | 232.2 | 0 | 32116.01172 |
| 0 | Q9UKN8 | General transcription factor 3C polypeptide 4 OS=Homo sapiens OX=9606 GN=GTF3C4 PE=1 SV=2 | 4 | 1 | 1 | 1 | 822 | 91.9 | 0 | 156948.8906 |
| 0 | Q9NR99 | Matrix-remodeling-associated protein 5 OS=Homo sapiens OX=9606 GN=MXRA5 PE=1 SV=3 | 0 | 1 | 1 | 1 | 2828 | 312 | 0 | 238410.4844 |
| 0 | Q9P0W2 | SWI/SNF-related matrix-associated actin-dependent regulator of chromatin subfamily E member 1-related OS=Homo sapiens OX=9606 GN=HMG20B PE=1 SV=1 | 13 | 1 | 1 | 1 | 317 | 35.8 | 0 | |
| 0 | Q5VVJ2 | Deubiquitinase MYSM1 OS=Homo sapiens OX=9606 GN=MYSM1 PE=1 SV=1 | 2 | 1 | 1 | 1 | 828 | 95 | 0 | 37902.07813 |
| 0 | Q96F81 | Protein dispatched homolog 1 OS=Homo sapiens OX=9606 GN=DISP1 PE=1 SV=3 | 1 | 1 | 1 | 1 | 1524 | 170.8 | 0 | 13693.96289 |
| 0 | Q5VST9 | Obscurin OS=Homo sapiens OX=9606 GN=OBSCN PE=1 SV=3 | 0 | 1 | 1 | 1 | 7968 | 867.9 | 0 | 68283.84375 |
| 0 | Q14515 | SPARC-like protein 1 OS=Homo sapiens OX=9606 GN=SPARCL1 PE=1 SV=2 | 5 | 1 | 1 | 1 | 664 | 75.2 | 0 | 31386.68359 |
| 0 | P63244 | Receptor of activated protein C kinase 1 OS=Homo sapiens OX=9606 GN=RACK1 PE=1 SV=3 | 10 | 2 | 2 | 2 | 317 | 35.1 | 0 | |
| 0 | Q9NWD8 | Transmembrane protein 248 OS=Homo sapiens OX=9606 GN=TMEM248 PE=1 SV=1 | 5 | 1 | 1 | 1 | 314 | 35 | 0 | 26521.71094 |
| 0 | P32969 | 60S ribosomal protein L9 OS=Homo sapiens OX=9606 GN=RPL9 PE=1 SV=1 | 9 | 1 | 1 | 1 | 192 | 21.9 | 0 | 59753.49609 |
| 0 | P41180 | Extracellular calcium-sensing receptor OS=Homo sapiens OX=9606 GN=CASR PE=1 SV=3 | 2 | 1 | 1 | 1 | 1078 | 120.6 | 0 | |
| 0 | Q4ZHG4 | Fibronectin type III domain-containing protein 1 OS=Homo sapiens OX=9606 GN=FNDC1 PE=2 SV=4 | 0 | 1 | 1 | 1 | 1894 | 205.4 | 0 | 25916.36719 |
| 0 | P13073 | Cytochrome c oxidase subunit 4 isoform 1, mitochondrial OS=Homo sapiens OX=9606 GN=COX4I1 PE=1 SV=1 | 7 | 1 | 1 | 1 | 169 | 19.6 | 0 | 43488.86328 |
| 0 | Q8WZ42 | Titin OS=Homo sapiens OX=9606 GN=TTN PE=1 SV=4 | 0 | 1 | 1 | 1 | 34350 | 3813.7 | 0 | 22914.69141 |
| 0 | Q7RTP6 | [F-actin]-monooxygenase MICAL3 OS=Homo sapiens OX=9606 GN=MICAL3 PE=1 SV=2 | 1 | 1 | 1 | 1 | 2002 | 224.2 | 0 | |
| 0 | P35568 | Insulin receptor substrate 1 OS=Homo sapiens OX=9606 GN=IRS1 PE=1 SV=1 | 1 | 1 | 1 | 1 | 1242 | 131.5 | 0 | 17398.06641 |
| 0 | O94864 | STAGA complex 65 subunit gamma OS=Homo sapiens OX=9606 GN=SUPT7L PE=1 SV=1 | 7 | 1 | 1 | 1 | 414 | 46.2 | 0 | |
| 0 | Q96DT5 | Dynein heavy chain 11, axonemal OS=Homo sapiens OX=9606 GN=DNAH11 PE=1 SV=4 | 1 | 1 | 1 | 1 | 4516 | 520 | 0 | 94982.52344 |
| 0 | P61421 | V-type proton ATPase subunit d 1 OS=Homo sapiens OX=9606 GN=ATP6V0D1 PE=1 SV=1 | 3 | 1 | 1 | 1 | 351 | 40.3 | 0 | 25656.75391 |
| 0 | Q92839 | Hyaluronan synthase 1 OS=Homo sapiens OX=9606 GN=HAS1 PE=1 SV=2 | 5 | 1 | 1 | 1 | 578 | 64.8 | 0 | |
| 0 | Q13347 | Eukaryotic translation initiation factor 3 subunit I OS=Homo sapiens OX=9606 GN=EIF3I PE=1 SV=1 | 3 | 1 | 1 | 1 | 325 | 36.5 | 0 | 15682.25098 |
| 0 | Q8TBR7 | TLC domain-containing protein 3A OS=Homo sapiens OX=9606 GN=TLCD3A PE=1 SV=2 | 14 | 1 | 1 | 1 | 257 | 29.4 | 0 | 19248.0332 |

Table S2 Label-free quantitative mass spectrometry reveals an OGT- 2CS interactome

| # Protein Pathway Groups | Accession | Description | Coverage [%] | # Peptides | # PSMs | # Unique Peptides | # AAs | MW [kDa] | Score Sequest HT: Sequest HT | Abundance: F3: Sample |
|---|---|---|---|---|---|---|---|---|---|---|
| 0 | P35579 | Myosin-9 OS=Homo sapiens OX=9606 GN=MYH9 PE=1 SV=4 | 60 | 126 | 179 | 117 | 1960 | 226.4 | 630.21 | 12916810.47 |
| 0 | O15294 | UDP-N-acetylglucosamine--peptide N-acetylglucosaminyltransferase 110 kDa subunit OS=Homo sapiens OX=9606 GN=OGT PE=1 SV=3 | 66 | 80 | 156 | 80 | 1046 | 116.9 | 488.6 | 27978288.63 |
| 0 | P14174 | Macrophage migration inhibitory factor OS=Homo sapiens OX=9606 GN=MIF PE=1 SV=4 | 52 | 9 | 153 | 9 | 115 | 12.5 | 401.75 | 43822655.61 |
| 0 | Q13813 | Spectrin alpha chain, non-erythrocytic 1 OS=Homo sapiens OX=9606 GN=SPTAN1 PE=1 SV=3 | 41 | 78 | 100 | 78 | 2472 | 284.4 | 339.65 | 4359488.286 |
| 0 | P08670 | Vimentin OS=Homo sapiens OX=9606 GN=VIM PE=1 SV=4 | 83 | 57 | 109 | 54 | 466 | 53.6 | 306.4 | 19824768.26 |
| 0 | P04264 | Keratin, type II cytoskeletal 1 OS=Homo sapiens OX=9606 GN=KRT1 PE=1 SV=6 | 68 | 48 | 94 | 43 | 644 | 66 | 305.53 | 17184705.56 |
| 0 | P63261 | Actin, cytoplasmic 2 OS=Homo sapiens OX=9606 GN=ACTG1 PE=1 SV=1 | 76 | 35 | 79 | 2 | 375 | 41.8 | 295.34 | 499032.833 |
| 0 | P60709 | Actin, cytoplasmic 1 OS=Homo sapiens OX=9606 GN=ACTB PE=1 SV=1 | 76 | 35 | 78 | 2 | 375 | 41.7 | 289.19 | 20996560.75 |
| 0 | P35527 | Keratin, type I cytoskeletal 9 OS=Homo sapiens OX=9606 GN=KRT9 PE=1 SV=3 | 61 | 32 | 72 | 32 | 623 | 62 | 284.84 | 8394035.882 |
| 0 | P0DMV9 | Heat shock 70 kDa protein 1B OS=Homo sapiens OX=9606 GN=HSPA1B PE=1 SV=1 | 69 | 51 | 82 | 38 | 641 | 70 | 272.02 | 12445287.98 |
| 0 | P35580 | Myosin-10 OS=Homo sapiens OX=9606 GN=MYH10 PE=1 SV=3 | 34 | 56 | 75 | 47 | 1976 | 228.9 | 257.62 | 2140014.11 |
| 0 | P11142 | Heat shock cognate 71 kDa protein OS=Homo sapiens OX=9606 GN=HSPA8 PE=1 SV=1 | 57 | 42 | 68 | 34 | 646 | 70.9 | 231.91 | 8355437.957 |
| 0 | P68133 | Actin, alpha skeletal muscle OS=Homo sapiens OX=9606 GN=ACTA1 PE=1 SV=1 | 42 | 20 | 62 | 4 | 377 | 42 | 219.79 | 300095.5942 |
| 0 | P13645 | Keratin, type I cytoskeletal 10 OS=Homo sapiens OX=9606 GN=KRT10 PE=1 SV=6 | 58 | 32 | 59 | 27 | 584 | 58.8 | 219.65 | 7166358.258 |
| 0 | P07910 | Heterogeneous nuclear ribonucleoproteins C1/C2 OS=Homo sapiens OX=9606 GN=HNRNPC PE=1 SV=4 | 58 | 26 | 66 | 24 | 306 | 33.7 | 208.86 | 14322165.51 |
| 0 | P16403 | Histone H1.2 OS=Homo sapiens OX=9606 GN=H1-2 PE=1 SV=2 | 65 | 26 | 66 | 10 | 213 | 21.4 | 181.85 | 26373089.63 |
| 0 | P35908 | Keratin, type II cytoskeletal 2 epidermal OS=Homo sapiens OX=9606 GN=KRT2 PE=1 SV=2 | 78 | 37 | 52 | 28 | 639 | 65.4 | 158.79 | 3545332.308 |
| 0 | P51610 | Host cell factor 1 OS=Homo sapiens OX=9606 GN=HCFC1 PE=1 SV=2 | 28 | 39 | 50 | 39 | 2035 | 208.6 | 148.63 | 3002191.35 |
| 0 | P07197 | Neurofilament medium polypeptide OS=Homo sapiens OX=9606 GN=NEFM PE=1 SV=3 | 35 | 27 | 45 | 25 | 916 | 102.4 | 139.87 | 1518662.399 |
| 0 | P12956 | X-ray repair cross-complementing protein 6 OS=Homo sapiens OX=9606 GN=XRCC6 PE=1 SV=2 | 36 | 24 | 43 | 24 | 609 | 69.8 | 134.31 | 2285164.061 |
| 0 | P19338 | Nucleolin OS=Homo sapiens OX=9606 GN=NCL PE=1 SV=3 | 39 | 26 | 39 | 26 | 710 | 76.6 | 133.09 | 2908062.188 |
| 0 | P10412 | Histone H1.4 OS=Homo sapiens OX=9606 GN=H1-4 PE=1 SV=2 | 58 | 22 | 47 | 5 | 219 | 21.9 | 132.43 | 4914115.26 |
| 0 | P46109 | Crk-like protein OS=Homo sapiens OX=9606 GN=CRKL PE=1 SV=1 | 86 | 22 | 39 | 22 | 303 | 33.8 | 129.13 | 3253379.125 |
| 0 | P16402 | Histone H1.3 OS=Homo sapiens OX=9606 GN=H1-3 PE=1 SV=2 | 48 | 19 | 43 | 3 | 221 | 22.3 | 125.96 | 383648.3857 |
| 0 | P17028 | Zinc finger protein 24 OS=Homo sapiens OX=9606 GN=ZNF24 PE=1 SV=4 | 70 | 22 | 41 | 22 | 368 | 42.1 | 122.37 | 6926237.824 |
| 0 | Q9BY77 | Polymerase delta-interacting protein 3 OS=Homo sapiens OX=9606 GN=POLDIP3 PE=1 SV=2 | 76 | 34 | 45 | 34 | 421 | 46.1 | 119.99 | 5491914.195 |
| 0 | P0DJI8 | Serum amyloid A-1 protein OS=Homo sapiens OX=9606 GN=SAA1 PE=1 SV=1 | 79 | 11 | 39 | 11 | 122 | 13.5 | 119.64 | 12678586.08 |
| 0 | Q86V81 | THO complex subunit 4 OS=Homo sapiens OX=9606 GN=ALYREF PE=1 SV=3 | 66 | 18 | 34 | 18 | 257 | 26.9 | 111.78 | 8022198.086 |
| 0 | P11021 | Endoplasmic reticulum chaperone BiP OS=Homo sapiens OX=9606 GN=HSPA5 PE=1 SV=2 | 48 | 28 | 35 | 27 | 654 | 72.3 | 111.38 | 2139181.481 |
| 0 | Q01082 | Spectrin beta chain, non-erythrocytic 1 OS=Homo sapiens OX=9606 GN=SPTBN1 PE=1 SV=2 | 16 | 24 | 31 | 23 | 2364 | 274.4 | 111.3 | 550576.582 |
| 0 | P11940 | Polyadenylate-binding protein 1 OS=Homo sapiens OX=9606 GN=PABPC1 PE=1 SV=2 | 48 | 29 | 36 | 18 | 636 | 70.6 | 108.18 | 2489825.059 |
| 0 | P38919 | Eukaryotic initiation factor 4A-III OS=Homo sapiens OX=9606 GN=EIF4A3 PE=1 SV=4 | 55 | 25 | 36 | 23 | 411 | 46.8 | 107.72 | 4008360.004 |
| 0 | P08779 | Keratin, type I cytoskeletal 16 OS=Homo sapiens OX=9606 GN=KRT16 PE=1 SV=4 | 47 | 23 | 34 | 10 | 473 | 51.2 | 105.94 | 1284567.92 |
| 0 | P13647 | Keratin, type II cytoskeletal 5 OS=Homo sapiens OX=9606 GN=KRT5 PE=1 SV=3 | 33 | 23 | 32 | 12 | 590 | 62.3 | 103.99 | 986287.5811 |
| 0 | Q9Y3Y2 | Chromatin target of PRMT1 protein OS=Homo sapiens OX=9606 GN=CHTOP PE=1 SV=2 | 53 | 18 | 37 | 18 | 248 | 26.4 | 103.97 | 14088930.43 |
| 0 | P02538 | Keratin, type II cytoskeletal 6A OS=Homo sapiens OX=9606 GN=KRT6A PE=1 SV=3 | 43 | 24 | 32 | 4 | 564 | 60 | 103.13 | 1277320.293 |
| 0 | P04259 | Keratin, type II cytoskeletal 6B OS=Homo sapiens OX=9606 GN=KRT6B PE=1 SV=5 | 39 | 23 | 31 | 1 | 564 | 60 | 97.93 | |
| 0 | P60660 | Myosin light polypeptide 6 OS=Homo sapiens OX=9606 GN=MYL6 PE=1 SV=2 | 68 | 17 | 30 | 14 | 151 | 16.9 | 96.09 | 4456859.245 |
| 0 | O60814 | Histone H2B type 1-K OS=Homo sapiens OX=9606 GN=H2BC12 PE=1 SV=3 | 84 | 19 | 37 | 2 | 126 | 13.9 | 93.25 | 8727442.927 |
| 0 | Q96FV9 | THO complex subunit 1 OS=Homo sapiens OX=9606 GN=THOC1 PE=1 SV=1 | 48 | 20 | 25 | 20 | 657 | 75.6 | 89.63 | 1256081.992 |
| 0 | P62269 | 40S ribosomal protein S18 OS=Homo sapiens OX=9606 GN=RPS18 PE=1 SV=3 | 76 | 23 | 34 | 23 | 152 | 17.7 | 88.57 | 10074227.9 |
| 0 | Q5QNW6 | Histone H2B type 2-F OS=Homo sapiens OX=9606 GN=H2BC18 PE=1 SV=3 | 84 | 18 | 34 | 1 | 126 | 13.9 | 87.67 | 134330.1719 |
| 0 | P58876 | Histone H2B type 1-D OS=Homo sapiens OX=9606 GN=H2BC5 PE=1 SV=2 | 84 | 18 | 34 | 1 | 126 | 13.9 | 87.17 | 208277.8203 |
| 0 | P67809 | Y-box-binding protein 1 OS=Homo sapiens OX=9606 GN=YBX1 PE=1 SV=3 | 57 | 15 | 28 | 10 | 324 | 35.9 | 87.1 | 1212366.74 |
| 0 | Q16778 | Histone H2B type 2-E OS=Homo sapiens OX=9606 GN=H2BC21 PE=1 SV=3 | 84 | 17 | 31 | 0 | 126 | 13.9 | 84.51 | |
| 0 | P38159 | RNA-binding motif protein, X chromosome OS=Homo sapiens OX=9606 GN=RBMX PE=1 SV=3 | 44 | 18 | 28 | 18 | 391 | 42.3 | 82.71 | 1074642.763 |
| 0 | P33778 | Histone H2B type 1-B OS=Homo sapiens OX=9606 GN=H2BC3 PE=1 SV=2 | 84 | 17 | 30 | 1 | 126 | 13.9 | 82.36 | 324754.0078 |
| 0 | P06748 | Nucleophosmin OS=Homo sapiens OX=9606 GN=NPM1 PE=1 SV=2 | 70 | 15 | 25 | 15 | 294 | 32.6 | 79.59 | 1654122.33 |
| 0 | Q9NYU2 | UDP-glucose:glycoprotein glucosyltransferase 1 OS=Homo sapiens OX=9606 GN=UGGT1 PE=1 SV=3 | 25 | 25 | 26 | 25 | 1555 | 177.1 | 78.94 | 595287.9834 |
| 0 | Q8NC51 | Plasminogen activator inhibitor 1 RNA-binding protein OS=Homo sapiens OX=9606 GN=SERBP1 PE=1 SV=2 | 40 | 17 | 23 | 17 | 408 | 44.9 | 77.34 | 2298037.672 |
| 0 | Q02539 | Histone H1.1 OS=Homo sapiens OX=9606 GN=H1-1 PE=1 SV=3 | 22 | 10 | 24 | 2 | 215 | 21.8 | 75.32 | 63444.96094 |
| 0 | P22626 | Heterogeneous nuclear ribonucleoproteins A2/B1 OS=Homo sapiens OX=9606 GN=HNRNPA2B1 PE=1 SV=2 | 58 | 18 | 25 | 18 | 353 | 37.4 | 75.04 | 1944629.859 |
| 0 | Q12906 | Interleukin enhancer-binding factor 3 OS=Homo sapiens OX=9606 GN=ILF3 PE=1 SV=3 | 26 | 17 | 25 | 17 | 894 | 95.3 | 74.47 | 1549585.389 |
| 0 | P02533 | Keratin, type I cytoskeletal 14 OS=Homo sapiens OX=9606 GN=KRT14 PE=1 SV=4 | 39 | 19 | 27 | 6 | 472 | 51.5 | 72.1 | 527999.4512 |
| 0 | P05787 | Keratin, type II cytoskeletal 8 OS=Homo sapiens OX=9606 GN=KRT8 PE=1 SV=7 | 38 | 16 | 23 | 11 | 483 | 53.7 | 71.47 | 274350.2329 |
| 0 | P62805 | Histone H4 OS=Homo sapiens OX=9606 GN=H4C1 PE=1 SV=2 | 61 | 13 | 26 | 13 | 103 | 11.4 | 68.96 | 6739130.123 |
| 0 | Q15436 | Protein transport protein Sec23A OS=Homo sapiens OX=9606 GN=SEC23A PE=1 SV=2 | 37 | 18 | 24 | 16 | 765 | 86.1 | 68.29 | 771031.8115 |
| 0 | Q9NZI8 | Insulin-like growth factor 2 mRNA-binding protein 1 OS=Homo sapiens OX=9606 GN=IGF2BP1 PE=1 SV=2 | 39 | 19 | 23 | 16 | 577 | 63.4 | 67.25 | 1015748.732 |
| 0 | P84090 | Enhancer of rudimentary homolog OS=Homo sapiens OX=9606 GN=ERH PE=1 SV=1 | 63 | 11 | 25 | 11 | 104 | 12.3 | 67.21 | 26694747.54 |
| 0 | Q8NI27 | THO complex subunit 2 OS=Homo sapiens OX=9606 GN=THOC2 PE=1 SV=2 | 16 | 19 | 22 | 19 | 1593 | 182.7 | 67.19 | 799634.1904 |
| 0 | P16989 | Y-box-binding protein 3 OS=Homo sapiens OX=9606 GN=YBX3 PE=1 SV=4 | 33 | 9 | 18 | 4 | 372 | 40.1 | 66.3 | 193311.873 |
| 0 | Q13310 | Polyadenylate-binding protein 4 OS=Homo sapiens OX=9606 GN=PABPC4 PE=1 SV=1 | 27 | 17 | 19 | 6 | 644 | 70.7 | 66.17 | 262209.002 |
| 0 | P62750 | 60S ribosomal protein L23a OS=Homo sapiens OX=9606 GN=RPL23A PE=1 SV=1 | 56 | 15 | 24 | 15 | 156 | 17.7 | 66.1 | 3685451.969 |

| | | | | | | | | | | |
|---|---|---|---|---|---|---|---|---|---|---|
| 0 | Q8NEU8 | DCC-interacting protein 13-beta OS=Homo sapiens OX=9606 GN=APPL2 PE=1 SV=3 | 41 | 16 | 17 | 16 | 664 | 74.4 | 65.78 | 484281.3867 |
| 0 | P26373 | 60S ribosomal protein L13 OS=Homo sapiens OX=9606 GN=RPL13 PE=1 SV=4 | 57 | 17 | 22 | 17 | 211 | 24.2 | 65.67 | 4222586.648 |
| 0 | P61247 | 40S ribosomal protein S3a OS=Homo sapiens OX=9606 GN=RPS3A PE=1 SV=2 | 65 | 20 | 23 | 20 | 264 | 29.9 | 63.78 | 1308907.432 |
| 0 | Q13838 | Spliceosome RNA helicase DDX39B OS=Homo sapiens OX=9606 GN=DDX39B PE=1 SV=1 | 50 | 15 | 21 | 15 | 428 | 49 | 63.35 | 3202839.498 |
| 0 | P62829 | 60S ribosomal protein L23 OS=Homo sapiens OX=9606 GN=RPL23 PE=1 SV=1 | 49 | 9 | 19 | 9 | 140 | 14.9 | 59.67 | 799574.3828 |
| 0 | Q9BRP1 | Programmed cell death protein 2-like OS=Homo sapiens OX=9606 GN=PDCD2L PE=1 SV=1 | 46 | 14 | 20 | 14 | 358 | 39.4 | 58.29 | 1614575.268 |
| 0 | Q9NYF8 | Bcl-2-associated transcription factor 1 OS=Homo sapiens OX=9606 GN=BCLAF1 PE=1 SV=2 | 23 | 19 | 22 | 19 | 920 | 106.1 | 57.72 | 888922.6162 |
| 0 | P68400 | Casein kinase II subunit alpha OS=Homo sapiens OX=9606 GN=CSNK2A1 PE=1 SV=1 | 42 | 12 | 15 | 12 | 391 | 45.1 | 57.67 | 378792.9883 |
| 0 | P09493 | Tropomyosin alpha-1 chain OS=Homo sapiens OX=9606 GN=TPM1 PE=1 SV=2 | 37 | 13 | 18 | 3 | 284 | 32.7 | 57.59 | 1262113.643 |
| 0 | Q06830 | Peroxiredoxin-1 OS=Homo sapiens OX=9606 GN=PRDX1 PE=1 SV=1 | 73 | 13 | 17 | 12 | 199 | 22.1 | 56.76 | 2313117.617 |
| 0 | P09651 | Heterogeneous nuclear ribonucleoprotein A1 OS=Homo sapiens OX=9606 GN=HNRNPA1 PE=1 SV=5 | 38 | 14 | 19 | 14 | 372 | 38.7 | 56.76 | 1872508.441 |
| 0 | Q13263 | Transcription intermediary factor 1-beta OS=Homo sapiens OX=9606 GN=TRIM28 PE=1 SV=5 | 30 | 12 | 18 | 12 | 835 | 88.5 | 56.16 | 410445.5449 |
| 0 | Q99878 | Histone H2A type 1-J OS=Homo sapiens OX=9606 GN=H2AC14 PE=1 SV=3 | 54 | 11 | 25 | 3 | 128 | 13.9 | 56.14 | 6446907.812 |
| 0 | Q9Y5S9 | RNA-binding protein 8A OS=Homo sapiens OX=9606 GN=RBM8A PE=1 SV=1 | 40 | 9 | 15 | 9 | 174 | 19.9 | 55.85 | 1449155.969 |
| 0 | Q13769 | THO complex subunit 5 homolog OS=Homo sapiens OX=9606 GN=THOC5 PE=1 SV=2 | 30 | 17 | 21 | 17 | 683 | 78.5 | 55.68 | 651707.8135 |
| 0 | Q9Y3I0 | RNA-splicing ligase RtcB homolog OS=Homo sapiens OX=9606 GN=RTCB PE=1 SV=1 | 43 | 13 | 17 | 13 | 505 | 55.2 | 55.66 | 618297.9341 |
| 0 | Q9UHX1 | Poly(U)-binding-splicing factor PUF60 OS=Homo sapiens OX=9606 GN=PUF60 PE=1 SV=1 | 28 | 8 | 15 | 8 | 559 | 59.8 | 54.2 | 342998.8105 |
| 0 | Q9UJV9 | Probable ATP-dependent RNA helicase DDX41 OS=Homo sapiens OX=9606 GN=DDX41 PE=1 SV=2 | 34 | 17 | 19 | 17 | 622 | 69.8 | 54.03 | 655748.1484 |
| 0 | Q99497 | Parkinson disease protein 7 OS=Homo sapiens OX=9606 GN=PARK7 PE=1 SV=2 | 71 | 11 | 15 | 11 | 189 | 19.9 | 53.2 | 721590.3076 |
| 0 | Q9UKV3 | Apoptotic chromatin condensation inducer in the nucleus OS=Homo sapiens OX=9606 GN=ACIN1 PE=1 SV=2 | 15 | 13 | 17 | 13 | 1341 | 151.8 | 53.01 | 205686.9971 |
| 0 | O15020 | Spectrin beta chain, non-erythrocytic 2 OS=Homo sapiens OX=9606 GN=SPTBN2 PE=1 SV=3 | 9 | 13 | 15 | 12 | 2390 | 271.2 | 51.75 | 96647.95117 |
| 0 | P08708 | 40S ribosomal protein S17 OS=Homo sapiens OX=9606 GN=RPS17 PE=1 SV=2 | 70 | 11 | 15 | 11 | 135 | 15.5 | 51.6 | 2242218.537 |
| 0 | P17066 | Heat shock 70 kDa protein 6 OS=Homo sapiens OX=9606 GN=HSPA6 PE=1 SV=2 | 17 | 12 | 17 | 1 | 643 | 71 | 51.38 | 396176.0313 |
| 0 | Q13435 | Splicing factor 3B subunit 2 OS=Homo sapiens OX=9606 GN=SF3B2 PE=1 SV=2 | 30 | 14 | 19 | 14 | 895 | 100.2 | 51.36 | 483089.1787 |
| 0 | P07437 | Tubulin beta chain OS=Homo sapiens OX=9606 GN=TUBB PE=1 SV=2 | 41 | 13 | 16 | 3 | 444 | 49.6 | 51.35 | 418893.6406 |
| 0 | P05387 | 60S acidic ribosomal protein P2 OS=Homo sapiens OX=9606 GN=RPLP2 PE=1 SV=1 | 85 | 9 | 13 | 9 | 115 | 11.7 | 50.84 | 240396.75 |
| 0 | Q15084 | Protein disulfide-isomerase A6 OS=Homo sapiens OX=9606 GN=PDIA6 PE=1 SV=1 | 30 | 12 | 14 | 12 | 440 | 48.1 | 49.36 | 700287.5044 |
| 0 | P07951 | Tropomyosin beta chain OS=Homo sapiens OX=9606 GN=TPM2 PE=1 SV=1 | 26 | 11 | 16 | 1 | 284 | 32.8 | 49.14 | 10408.60352 |
| 0 | P07196 | Neurofilament light polypeptide OS=Homo sapiens OX=9606 GN=NEFL PE=1 SV=3 | 38 | 17 | 18 | 16 | 543 | 61.5 | 48.38 | 490017.9658 |
| 0 | Q9Y2W1 | Thyroid hormone receptor-associated protein 3 OS=Homo sapiens OX=9606 GN=THRAP3 PE=1 SV=2 | 21 | 15 | 17 | 15 | 955 | 108.6 | 48.26 | 973534.751 |
| 0 | Q96QD9 | UAP56-interacting factor OS=Homo sapiens OX=9606 GN=FYTTD1 PE=1 SV=3 | 59 | 15 | 17 | 15 | 318 | 35.8 | 48.19 | 886516.582 |
| 0 | P08865 | 40S ribosomal protein SA OS=Homo sapiens OX=9606 GN=RPSA PE=1 SV=4 | 57 | 14 | 16 | 14 | 295 | 32.8 | 48.09 | 1071476.564 |
| 0 | Q9ULU4 | Protein kinase C-binding protein 1 OS=Homo sapiens OX=9606 GN=ZMYND8 PE=1 SV=2 | 21 | 13 | 13 | 13 | 1186 | 131.6 | 46.57 | 299550.6377 |
| 0 | Q12905 | Interleukin enhancer-binding factor 2 OS=Homo sapiens OX=9606 GN=ILF2 PE=1 SV=2 | 51 | 13 | 18 | 13 | 390 | 43 | 46.16 | 917102.9746 |
| 0 | Q9UKM9 | RNA-binding protein Raly OS=Homo sapiens OX=9606 GN=RALY PE=1 SV=1 | 56 | 15 | 16 | 15 | 306 | 32.4 | 45.49 | 403807.3652 |
| 0 | P61254 | 60S ribosomal protein L26 OS=Homo sapiens OX=9606 GN=RPL26 PE=1 SV=1 | 59 | 11 | 20 | 4 | 145 | 17.2 | 45.41 | 3095012.66 |
| 0 | P68371 | Tubulin beta-4B chain OS=Homo sapiens OX=9606 GN=TUBB4B PE=1 SV=1 | 34 | 11 | 14 | 1 | 445 | 49.8 | 44.98 | 22620.70508 |
| 0 | Q9BRD0 | BUD13 homolog OS=Homo sapiens OX=9606 GN=BUD13 PE=1 SV=1 | 23 | 14 | 17 | 14 | 619 | 70.5 | 43.85 | 1057584.686 |
| 0 | Q08211 | ATP-dependent RNA helicase A OS=Homo sapiens OX=9606 GN=DHX9 PE=1 SV=4 | 14 | 15 | 17 | 15 | 1270 | 140.9 | 43.75 | 325606.4844 |
| 0 | P13010 | X-ray repair cross-complementing protein 5 OS=Homo sapiens OX=9606 GN=XRCC5 PE=1 SV=3 | 25 | 11 | 12 | 11 | 732 | 82.7 | 42.91 | 397084.4512 |
| 0 | O00422 | Histone deacetylase complex subunit SAP18 OS=Homo sapiens OX=9606 GN=SAP18 PE=1 SV=1 | 64 | 11 | 16 | 11 | 153 | 17.6 | 42.56 | 2005582.773 |
| 0 | Q86U42 | Polyadenylate-binding protein 2 OS=Homo sapiens OX=9606 GN=PABPN1 PE=1 SV=3 | 44 | 8 | 14 | 8 | 306 | 32.7 | 42.37 | 3584632.799 |
| 0 | P62081 | 40S ribosomal protein S7 OS=Homo sapiens OX=9606 GN=RPS7 PE=1 SV=1 | 56 | 9 | 13 | 9 | 194 | 22.1 | 42 | 692331.0107 |
| 0 | Q13573 | SNW domain-containing protein 1 OS=Homo sapiens OX=9606 GN=SNW1 PE=1 SV=1 | 41 | 12 | 14 | 12 | 536 | 61.5 | 41.81 | 100801.2627 |
| 0 | Q93077 | Histone H2A type 1-C OS=Homo sapiens OX=9606 GN=H2AC6 PE=1 SV=3 | 53 | 9 | 18 | 1 | 130 | 14.1 | 41.1 | 79243.26953 |
| 0 | P52907 | F-actin-capping protein subunit alpha-1 OS=Homo sapiens OX=9606 GN=CAPZA1 PE=1 SV=3 | 48 | 10 | 13 | 6 | 286 | 32.9 | 40.99 | 641633.623 |
| 0 | P62318 | Small nuclear ribonucleoprotein Sm D3 OS=Homo sapiens OX=9606 GN=SNRPD3 PE=1 SV=1 | 62 | 7 | 15 | 7 | 126 | 13.9 | 40.77 | 972016.834 |
| 0 | O75533 | Splicing factor 3B subunit 1 OS=Homo sapiens OX=9606 GN=SF3B1 PE=1 SV=3 | 17 | 12 | 15 | 12 | 1304 | 145.7 | 40.75 | 403198.9258 |
| 0 | Q9Y230 | RuvB-like 2 OS=Homo sapiens OX=9606 GN=RUVBL2 PE=1 SV=3 | 29 | 12 | 14 | 12 | 463 | 51.1 | 40.59 | 684811.2637 |
| 0 | Q15029 | 116 kDa U5 small nuclear ribonucleoprotein component OS=Homo sapiens OX=9606 GN=EFTUD2 PE=1 SV=1 | 19 | 12 | 13 | 12 | 972 | 109.4 | 40.5 | 189835.6504 |
| 0 | P06753 | Tropomyosin alpha-3 chain OS=Homo sapiens OX=9606 GN=TPM3 PE=1 SV=2 | 24 | 8 | 11 | 1 | 285 | 32.9 | 40.19 | 127750.5586 |
| 0 | P23396 | 40S ribosomal protein S3 OS=Homo sapiens OX=9606 GN=RPS3 PE=1 SV=2 | 46 | 11 | 14 | 11 | 243 | 26.7 | 39.74 | 716742.5293 |
| 0 | P12268 | Inosine-5'-monophosphate dehydrogenase 2 OS=Homo sapiens OX=9606 GN=IMPDH2 PE=1 SV=2 | 28 | 10 | 10 | 10 | 514 | 55.8 | 39.24 | 463615.1982 |
| 0 | P62424 | 60S ribosomal protein L7a OS=Homo sapiens OX=9606 GN=RPL7A PE=1 SV=2 | 55 | 13 | 19 | 13 | 266 | 30 | 39.22 | 1327987.047 |
| 0 | Q07955 | Serine/arginine-rich splicing factor 1 OS=Homo sapiens OX=9606 GN=SRSF1 PE=1 SV=2 | 56 | 11 | 15 | 10 | 248 | 27.7 | 39.19 | 1504574.858 |
| 0 | P67936 | Tropomyosin alpha-4 chain OS=Homo sapiens OX=9606 GN=TPM4 PE=1 SV=3 | 29 | 8 | 12 | 1 | 248 | 28.5 | 39 | |
| 0 | P61978 | Heterogeneous nuclear ribonucleoprotein K OS=Homo sapiens OX=9606 GN=HNRNPK PE=1 SV=1 | 39 | 12 | 15 | 12 | 463 | 50.9 | 38.47 | 624254.8691 |
| 0 | P62826 | GTP-binding nuclear protein Ran OS=Homo sapiens OX=9606 GN=RAN PE=1 SV=3 | 40 | 8 | 14 | 8 | 216 | 24.4 | 38.33 | 670623.4834 |
| 0 | Q92522 | Histone H1.10 OS=Homo sapiens OX=9606 GN=H1-10 PE=1 SV=1 | 48 | 13 | 15 | 13 | 213 | 22.5 | 37.1 | 1493754.504 |
| 0 | P22234 | Multifunctional protein ADE2 OS=Homo sapiens OX=9606 GN=PAICS PE=1 SV=3 | 33 | 9 | 11 | 9 | 425 | 47 | 37.07 | 739480.7842 |
| 0 | Q9Y224 | RNA transcription, translation and transport factor protein OS=Homo sapiens OX=9606 GN=RTRAF PE=1 SV=1 | 34 | 6 | 9 | 6 | 244 | 28.1 | 36.72 | 213994.9785 |
| 0 | P31040 | Succinate dehydrogenase [ubiquinone] flavoprotein subunit, mitochondrial OS=Homo sapiens OX=9606 GN=SDHA PE=1 SV=2 | 25 | 9 | 11 | 9 | 664 | 72.6 | 36.69 | 352959.1406 |
| 0 | P84243 | Histone H3.3 OS=Homo sapiens OX=9606 GN=H3-3A PE=1 SV=2 | 64 | 11 | 15 | 2 | 136 | 15.3 | 36.6 | 4840525.446 |
| 0 | Q9UHV9 | Prefoldin subunit 2 OS=Homo sapiens OX=9606 GN=PFDN2 PE=1 SV=1 | 59 | 7 | 10 | 7 | 154 | 16.6 | 36.31 | 726907.5898 |
| 0 | O60884 | DnaJ homolog subfamily A member 2 OS=Homo sapiens OX=9606 GN=DNAJA2 PE=1 SV=1 | 32 | 10 | 13 | 10 | 412 | 45.7 | 35.77 | 564244.188 |

| | | | | | | | | | | |
|---|---|---|---|---|---|---|---|---|---|---|
| 0 | P29692 | Elongation factor 1-delta OS=Homo sapiens OX=9606 GN=EEF1D PE=1 SV=5 | 34 | 8 | 10 | 6 | 281 | 31.1 | 35.74 | 214540.5742 |
| 0 | Q00839 | Heterogeneous nuclear ribonucleoprotein U OS=Homo sapiens OX=9606 GN=HNRNPU PE=1 SV=6 | 20 | 10 | 11 | 10 | 825 | 90.5 | 35.35 | 254519.8379 |
| 0 | P0DP25 | Calmodulin-3 OS=Homo sapiens OX=9606 GN=CALM3 PE=1 SV=1 | 46 | 6 | 11 | 6 | 149 | 16.8 | 34.84 | 1233150.45 |
| 0 | P62263 | 40S ribosomal protein S14 OS=Homo sapiens OX=9606 GN=RPS14 PE=1 SV=3 | 42 | 8 | 11 | 8 | 151 | 16.3 | 34.8 | 1055959.219 |
| 0 | P62753 | 40S ribosomal protein S6 OS=Homo sapiens OX=9606 GN=RPS6 PE=1 SV=1 | 34 | 8 | 12 | 8 | 249 | 28.7 | 34.78 | 386985.9375 |
| 0 | O75152 | Zinc finger CCCH domain-containing protein 11A OS=Homo sapiens OX=9606 GN=ZC3H11A PE=1 SV=3 | 21 | 12 | 13 | 12 | 810 | 89.1 | 34.63 | 248697.6172 |
| 0 | Q96J01 | THO complex subunit 3 OS=Homo sapiens OX=9606 GN=THOC3 PE=1 SV=1 | 40 | 10 | 12 | 10 | 351 | 38.7 | 34.49 | 449659.6113 |
| 0 | P10809 | 60 kDa heat shock protein, mitochondrial OS=Homo sapiens OX=9606 GN=HSPD1 PE=1 SV=2 | 21 | 7 | 10 | 7 | 573 | 61 | 34.41 | 52378.25195 |
| 0 | Q9H307 | Pinin OS=Homo sapiens OX=9606 GN=PNN PE=1 SV=5 | 16 | 10 | 12 | 10 | 717 | 81.6 | 34.25 | 399474.0156 |
| 0 | P48634 | Protein PRRC2A OS=Homo sapiens OX=9606 GN=PRRC2A PE=1 SV=3 | 8 | 9 | 11 | 9 | 2157 | 228.7 | 34.06 | 312472.1904 |
| 0 | P46779 | 60S ribosomal protein L28 OS=Homo sapiens OX=9606 GN=RPL28 PE=1 SV=3 | 59 | 8 | 14 | 8 | 137 | 15.7 | 33.83 | 1071479.432 |
| 0 | P23528 | Cofilin-1 OS=Homo sapiens OX=9606 GN=CFL1 PE=1 SV=3 | 69 | 9 | 10 | 5 | 166 | 18.5 | 33.61 | 342708.1523 |
| 0 | P25398 | 40S ribosomal protein S12 OS=Homo sapiens OX=9606 GN=RPS12 PE=1 SV=3 | 83 | 10 | 11 | 10 | 132 | 14.5 | 33.12 | 627559.2861 |
| 0 | P62906 | 60S ribosomal protein L10a OS=Homo sapiens OX=9606 GN=RPL10A PE=1 SV=2 | 43 | 9 | 11 | 9 | 217 | 24.8 | 32.87 | 621366.8135 |
| 0 | P60866 | 40S ribosomal protein S20 OS=Homo sapiens OX=9606 GN=RPS20 PE=1 SV=1 | 34 | 5 | 11 | 5 | 119 | 13.4 | 32.82 | 1857605.34 |
| 0 | P09661 | U2 small nuclear ribonucleoprotein A' OS=Homo sapiens OX=9606 GN=SNRPA1 PE=1 SV=2 | 56 | 9 | 10 | 9 | 255 | 28.4 | 32.52 | 148509.7949 |
| 0 | P52272 | Heterogeneous nuclear ribonucleoprotein M OS=Homo sapiens OX=9606 GN=HNRNPM PE=1 SV=3 | 19 | 11 | 12 | 11 | 730 | 77.5 | 32.41 | 408005.0898 |
| 0 | Q96PK6 | RNA-binding protein 14 OS=Homo sapiens OX=9606 GN=RBM14 PE=1 SV=2 | 22 | 11 | 13 | 11 | 669 | 69.4 | 32.34 | 1263498.988 |
| 0 | Q6PJT7 | Zinc finger CCCH domain-containing protein 14 OS=Homo sapiens OX=9606 GN=ZC3H14 PE=1 SV=1 | 20 | 10 | 11 | 10 | 736 | 82.8 | 32.33 | 308603.834 |
| 0 | P84103 | Serine/arginine-rich splicing factor 3 OS=Homo sapiens OX=9606 GN=SRSF3 PE=1 SV=1 | 51 | 10 | 13 | 9 | 164 | 19.3 | 32.28 | 853810.918 |
| 0 | Q9UNX3 | 60S ribosomal protein L26-like 1 OS=Homo sapiens OX=9606 GN=RPL26L1 PE=1 SV=1 | 50 | 8 | 13 | 1 | 145 | 17.2 | 32.27 | 43928.28516 |
| 0 | P68104 | Elongation factor 1-alpha 1 OS=Homo sapiens OX=9606 GN=EEF1A1 PE=1 SV=1 | 32 | 7 | 9 | 7 | 462 | 50.1 | 31.94 | 287217.3994 |
| 0 | P05455 | Lupus La protein OS=Homo sapiens OX=9606 GN=SSB PE=1 SV=2 | 21 | 8 | 9 | 8 | 408 | 46.8 | 31.94 | 127281.2783 |
| 0 | Q08380 | Galectin-3-binding protein OS=Homo sapiens OX=9606 GN=LGALS3BP PE=1 SV=1 | 22 | 8 | 10 | 8 | 585 | 65.3 | 31.6 | 156262.1836 |
| 0 | P14866 | Heterogeneous nuclear ribonucleoprotein L OS=Homo sapiens OX=9606 GN=HNRNPL PE=1 SV=2 | 28 | 8 | 8 | 8 | 589 | 64.1 | 31.3 | 104377.7813 |
| 0 | P51991 | Heterogeneous nuclear ribonucleoprotein A3 OS=Homo sapiens OX=9606 GN=HNRNPA3 PE=1 SV=2 | 32 | 7 | 10 | 7 | 378 | 39.6 | 31.14 | 328770.3262 |
| 0 | Q71DI3 | Histone H3.2 OS=Homo sapiens OX=9606 GN=H3C15 PE=1 SV=3 | 64 | 10 | 14 | 1 | 136 | 15.4 | 30.8 | 6204.594727 |
| 0 | P62847 | 40S ribosomal protein S24 OS=Homo sapiens OX=9606 GN=RPS24 PE=1 SV=1 | 56 | 10 | 12 | 10 | 133 | 15.4 | 30.69 | 379503.8389 |
| 0 | P68431 | Histone H3.1 OS=Homo sapiens OX=9606 GN=H3C1 PE=1 SV=2 | 64 | 10 | 14 | 1 | 136 | 15.4 | 30.67 | 12068.2207 |
| 0 | P62899 | 60S ribosomal protein L31 OS=Homo sapiens OX=9606 GN=RPL31 PE=1 SV=1 | 52 | 10 | 12 | 10 | 125 | 14.5 | 30.61 | 1070995.947 |
| 0 | Q15459 | Splicing factor 3A subunit 1 OS=Homo sapiens OX=9606 GN=SF3A1 PE=1 SV=1 | 9 | 6 | 8 | 6 | 793 | 88.8 | 30.26 | 218725.2734 |
| 0 | Q15233 | Non-POU domain-containing octamer-binding protein OS=Homo sapiens OX=9606 GN=NONO PE=1 SV=4 | 22 | 8 | 10 | 8 | 471 | 54.2 | 29.87 | 211910.2715 |
| 0 | Q9Y3C6 | Peptidyl-prolyl cis-trans isomerase-like 1 OS=Homo sapiens OX=9606 GN=PPIL1 PE=1 SV=1 | 33 | 5 | 9 | 5 | 166 | 18.2 | 29.21 | 929985.9785 |
| 0 | P61326 | Protein mago nashi homolog OS=Homo sapiens OX=9606 GN=MAGOH PE=1 SV=1 | 39 | 6 | 11 | 6 | 146 | 17.2 | 28.96 | 1378058.492 |
| 0 | P11233 | Ras-related protein Ral-A OS=Homo sapiens OX=9606 GN=RALA PE=1 SV=1 | 54 | 10 | 10 | 10 | 206 | 23.6 | 28.89 | 451226.1895 |
| 0 | P62910 | 60S ribosomal protein L32 OS=Homo sapiens OX=9606 GN=RPL32 PE=1 SV=2 | 44 | 5 | 8 | 5 | 135 | 15.9 | 28.79 | 1009672.084 |
| 0 | Q86W42 | THO complex subunit 6 homolog OS=Homo sapiens OX=9606 GN=THOC6 PE=1 SV=1 | 24 | 7 | 9 | 7 | 341 | 37.5 | 28.6 | 441998.9004 |
| 0 | O95793 | Double-stranded RNA-binding protein Staufen homolog 1 OS=Homo sapiens OX=9606 GN=STAU1 PE=1 SV=2 | 19 | 6 | 9 | 6 | 577 | 63.1 | 28.51 | 143392.8379 |
| 0 | P24534 | Elongation factor 1-beta OS=Homo sapiens OX=9606 GN=EEF1B2 PE=1 SV=3 | 30 | 6 | 9 | 4 | 225 | 24.7 | 28.49 | 273212.6406 |
| 0 | P62979 | Ubiquitin-40S ribosomal protein S27a OS=Homo sapiens OX=9606 GN=RPS27A PE=1 SV=2 | 46 | 6 | 10 | 3 | 156 | 18 | 28.35 | 201755.8848 |
| 0 | Q9Y3U8 | 60S ribosomal protein L36 OS=Homo sapiens OX=9606 GN=RPL36 PE=1 SV=3 | 42 | 6 | 10 | 6 | 105 | 12.2 | 28.34 | 574399.8311 |
| 0 | Q15287 | RNA-binding protein with serine-rich domain 1 OS=Homo sapiens OX=9606 GN=RNPS1 PE=1 SV=1 | 32 | 10 | 10 | 10 | 305 | 34.2 | 27.93 | 620764.7344 |
| 0 | P09874 | Poly [ADP-ribose] polymerase 1 OS=Homo sapiens OX=9606 GN=PARP1 PE=1 SV=4 | 12 | 8 | 9 | 8 | 1014 | 113 | 27.41 | 107828.7061 |
| 0 | P63104 | 14-3-3 protein zeta/delta OS=Homo sapiens OX=9606 GN=YWHAZ PE=1 SV=1 | 26 | 6 | 9 | 6 | 245 | 27.7 | 27.36 | 172893.1943 |
| 0 | P62277 | 40S ribosomal protein S13 OS=Homo sapiens OX=9606 GN=RPS13 PE=1 SV=2 | 52 | 10 | 11 | 10 | 151 | 17.2 | 26.95 | 528915.625 |
| 0 | P62280 | 40S ribosomal protein S11 OS=Homo sapiens OX=9606 GN=RPS11 PE=1 SV=3 | 47 | 10 | 10 | 10 | 158 | 18.4 | 26.95 | 1179130.517 |
| 0 | P02794 | Ferritin heavy chain OS=Homo sapiens OX=9606 GN=FTH1 PE=1 SV=2 | 37 | 4 | 7 | 4 | 183 | 21.2 | 26.42 | 202885.6846 |
| 0 | P39019 | 40S ribosomal protein S19 OS=Homo sapiens OX=9606 GN=RPS19 PE=1 SV=2 | 44 | 10 | 10 | 10 | 145 | 16.1 | 26.38 | 817012.4531 |
| 0 | Q12765 | Secernin-1 OS=Homo sapiens OX=9606 GN=SCRN1 PE=1 SV=2 | 20 | 7 | 8 | 7 | 414 | 46.4 | 25.99 | 872988.6328 |
| 0 | P02768 | Albumin OS=Homo sapiens OX=9606 GN=ALB PE=1 SV=2 | 13 | 8 | 10 | 8 | 609 | 69.3 | 25.68 | 2026935.336 |
| 0 | P14649 | Myosin light chain 6B OS=Homo sapiens OX=9606 GN=MYL6B PE=1 SV=1 | 22 | 4 | 7 | 1 | 208 | 22.8 | 25.59 | 15489.125 |
| 0 | P62249 | 40S ribosomal protein S16 OS=Homo sapiens OX=9606 GN=RPS16 PE=1 SV=2 | 48 | 7 | 10 | 7 | 146 | 16.4 | 25.55 | 587181.1436 |
| 0 | Q02878 | 60S ribosomal protein L6 OS=Homo sapiens OX=9606 GN=RPL6 PE=1 SV=3 | 31 | 9 | 13 | 9 | 288 | 32.7 | 25.49 | 775567.3047 |
| 0 | P14618 | Pyruvate kinase PKM OS=Homo sapiens OX=9606 GN=PKM PE=1 SV=4 | 25 | 7 | 7 | 7 | 531 | 57.9 | 25.43 | 544783.1523 |
| 0 | Q86YZ3 | Hornerin OS=Homo sapiens OX=9606 GN=HRNR PE=1 SV=2 | 8 | 7 | 8 | 7 | 2850 | 282.2 | 25.16 | 71657.93994 |
| 0 | O14950 | Myosin regulatory light chain 12B OS=Homo sapiens OX=9606 GN=MYL12B PE=1 SV=2 | 47 | 7 | 7 | 7 | 172 | 19.8 | 25.14 | 901569.0898 |
| 0 | Q13151 | Heterogeneous nuclear ribonucleoprotein A0 OS=Homo sapiens OX=9606 GN=HNRNPA0 PE=1 SV=1 | 28 | 6 | 8 | 6 | 305 | 30.8 | 24.56 | 385398.7051 |
| 0 | O14979 | Heterogeneous nuclear ribonucleoprotein D-like OS=Homo sapiens OX=9606 GN=HNRNPDL PE=1 SV=3 | 21 | 7 | 9 | 6 | 420 | 46.4 | 24.48 | 224940.0049 |
| 0 | P67870 | Casein kinase II subunit beta OS=Homo sapiens OX=9606 GN=CSNK2B PE=1 SV=1 | 38 | 5 | 7 | 5 | 215 | 24.9 | 24.24 | 347950.5742 |
| 0 | P09012 | U1 small nuclear ribonucleoprotein A OS=Homo sapiens OX=9606 GN=SNRPA PE=1 SV=3 | 27 | 5 | 7 | 5 | 282 | 31.3 | 24.2 | 280629.8496 |
| 0 | P49327 | Fatty acid synthase OS=Homo sapiens OX=9606 GN=FASN PE=1 SV=3 | 4 | 5 | 7 | 5 | 2511 | 273.3 | 24.19 | 61123.94336 |
| 0 | Q9Y265 | RuvB-like 1 OS=Homo sapiens OX=9606 GN=RUVBL1 PE=1 SV=1 | 26 | 7 | 9 | 7 | 456 | 50.2 | 23.75 | 226647.0527 |
| 0 | P62841 | 40S ribosomal protein S15 OS=Homo sapiens OX=9606 GN=RPS15 PE=1 SV=2 | 54 | 7 | 8 | 7 | 145 | 17 | 23.73 | 611684.8779 |
| 0 | P62316 | Small nuclear ribonucleoprotein Sm D2 OS=Homo sapiens OX=9606 GN=SNRPD2 PE=1 SV=1 | 62 | 6 | 9 | 6 | 118 | 13.5 | 23.66 | 870714.0859 |
| 0 | P37108 | Signal recognition particle 14 kDa protein OS=Homo sapiens OX=9606 GN=SRP14 PE=1 SV=2 | 41 | 6 | 7 | 6 | 136 | 14.6 | 23.33 | 530261.9395 |
| 0 | P41208 | Centrin-2 OS=Homo sapiens OX=9606 GN=CETN2 PE=1 SV=1 | 42 | 6 | 6 | 6 | 172 | 19.7 | 23.2 | 184821.2324 |

| | | | | | | | | | | |
|---|---|---|---|---|---|---|---|---|---|---|
| 0 | P26599 | Polypyrimidine tract-binding protein 1 OS=Homo sapiens OX=9606 GN=PTBP1 PE=1 SV=1 | 17 | 7 | 8 | 7 | 531 | 57.2 | 23.06 | 358103.0088 |
| 0 | P62888 | 60S ribosomal protein L30 OS=Homo sapiens OX=9606 GN=RPL30 PE=1 SV=2 | 67 | 6 | 6 | 6 | 115 | 12.8 | 22.87 | 539968.3496 |
| 0 | P62937 | Peptidyl-prolyl cis-trans isomerase A OS=Homo sapiens OX=9606 GN=PPIA PE=1 SV=2 | 66 | 7 | 7 | 7 | 165 | 18 | 22.55 | 356001.0977 |
| 0 | P18621 | 60S ribosomal protein L17 OS=Homo sapiens OX=9606 GN=RPL17 PE=1 SV=3 | 56 | 7 | 8 | 7 | 184 | 21.4 | 22.52 | 403487.4395 |
| 0 | Q9UMS4 | Pre-mRNA-processing factor 19 OS=Homo sapiens OX=9606 GN=PRPF19 PE=1 SV=1 | 22 | 7 | 8 | 7 | 504 | 55.1 | 22.19 | 197034.0957 |
| 0 | Q07666 | KH domain-containing, RNA-binding, signal transduction-associated protein 1 OS=Homo sapiens OX=9606 GN=KHDRBS1 PE=1 SV=1 | 16 | 5 | 8 | 5 | 443 | 48.2 | 22.16 | 394483.7559 |
| 0 | P05783 | Keratin, type I cytoskeletal 18 OS=Homo sapiens OX=9606 GN=KRT18 PE=1 SV=2 | 26 | 8 | 9 | 7 | 430 | 48 | 22.1 | 153168.1436 |
| 0 | P55209 | Nucleosome assembly protein 1-like 1 OS=Homo sapiens OX=9606 GN=NAP1L1 PE=1 SV=1 | 24 | 6 | 6 | 6 | 391 | 45.3 | 21.81 | 95903.85645 |
| 0 | O14828 | Secretory carrier-associated membrane protein 3 OS=Homo sapiens OX=9606 GN=SCAMP3 PE=1 SV=3 | 30 | 6 | 7 | 6 | 347 | 38.3 | 21.73 | 336672.6406 |
| 0 | P62273 | 40S ribosomal protein S29 OS=Homo sapiens OX=9606 GN=RPS29 PE=1 SV=2 | 59 | 5 | 11 | 5 | 56 | 6.7 | 21.64 | 2410517.813 |
| 0 | P83731 | 60S ribosomal protein L24 OS=Homo sapiens OX=9606 GN=RPL24 PE=1 SV=1 | 46 | 7 | 8 | 7 | 157 | 17.8 | 21.31 | 605336.6777 |
| 0 | P31943 | Heterogeneous nuclear ribonucleoprotein H OS=Homo sapiens OX=9606 GN=HNRNPH1 PE=1 SV=4 | 16 | 6 | 6 | 6 | 449 | 49.2 | 21.31 | 242484.042 |
| 0 | P62701 | 40S ribosomal protein S4, X isoform OS=Homo sapiens OX=9606 GN=RPS4X PE=1 SV=2 | 27 | 8 | 9 | 8 | 263 | 29.6 | 21.29 | 639520.1289 |
| 0 | P17844 | Probable ATP-dependent RNA helicase DDX5 OS=Homo sapiens OX=9606 GN=DDX5 PE=1 SV=1 | 14 | 7 | 8 | 7 | 614 | 69.1 | 21.24 | 94654.97266 |
| 0 | P46778 | 60S ribosomal protein L21 OS=Homo sapiens OX=9606 GN=RPL21 PE=1 SV=2 | 47 | 7 | 7 | 7 | 160 | 18.6 | 21.19 | 533571.3203 |
| 0 | P62917 | 60S ribosomal protein L8 OS=Homo sapiens OX=9606 GN=RPL8 PE=1 SV=2 | 36 | 8 | 9 | 8 | 257 | 28 | 21.13 | 502897.8281 |
| 0 | Q15717 | ELAV-like protein 1 OS=Homo sapiens OX=9606 GN=ELAVL1 PE=1 SV=2 | 35 | 7 | 7 | 7 | 326 | 36.1 | 20.93 | 195528.0762 |
| 0 | P32119 | Peroxiredoxin-2 OS=Homo sapiens OX=9606 GN=PRDX2 PE=1 SV=5 | 39 | 5 | 7 | 4 | 198 | 21.9 | 20.79 | 101508.0098 |
| 0 | P61353 | 60S ribosomal protein L27 OS=Homo sapiens OX=9606 GN=RPL27 PE=1 SV=2 | 46 | 7 | 8 | 7 | 136 | 15.8 | 20.73 | 909007.0723 |
| 0 | P63173 | 60S ribosomal protein L38 OS=Homo sapiens OX=9606 GN=RPL38 PE=1 SV=2 | 57 | 5 | 6 | 5 | 70 | 8.2 | 20.46 | 985653.7695 |
| 0 | P14678 | Small nuclear ribonucleoprotein-associated proteins B and B' OS=Homo sapiens OX=9606 GN=SNRPB PE=1 SV=2 | 26 | 6 | 6 | 6 | 240 | 24.6 | 20.17 | 446990.3984 |
| 0 | Q6I9Y2 | THO complex subunit 7 homolog OS=Homo sapiens OX=9606 GN=THOC7 PE=1 SV=3 | 41 | 7 | 7 | 7 | 204 | 23.7 | 20.09 | 722226.7695 |
| 0 | Q14257 | Reticulocalbin-2 OS=Homo sapiens OX=9606 GN=RCN2 PE=1 SV=1 | 20 | 5 | 6 | 5 | 317 | 36.9 | 20.07 | 212321.2139 |
| 0 | P05204 | Non-histone chromosomal protein HMG-17 OS=Homo sapiens OX=9606 GN=HMGN2 PE=1 SV=3 | 17 | 1 | 9 | 1 | 90 | 9.4 | 19.73 | 320739.2329 |
| 0 | P42771 | Cyclin-dependent kinase inhibitor 2A OS=Homo sapiens OX=9606 GN=CDKN2A PE=1 SV=2 | 63 | 6 | 8 | 6 | 156 | 16.5 | 19.34 | 393151.7891 |
| 0 | Q96QV6 | Histone H2A type 1-A OS=Homo sapiens OX=9606 GN=H2AC1 PE=1 SV=3 | 45 | 6 | 9 | 1 | 131 | 14.2 | 19.03 | 16623.67773 |
| 0 | Q9NZT1 | Calmodulin-like protein 5 OS=Homo sapiens OX=9606 GN=CALML5 PE=1 SV=2 | 34 | 6 | 6 | 6 | 146 | 15.9 | 18.97 | 570321.9316 |
| 0 | Q99729 | Heterogeneous nuclear ribonucleoprotein A/B OS=Homo sapiens OX=9606 GN=HNRNPAB PE=1 SV=2 | 18 | 4 | 6 | 4 | 332 | 36.2 | 18.92 | 149044.9414 |
| 0 | Q6DD87 | Zinc finger protein 787 OS=Homo sapiens OX=9606 GN=ZNF787 PE=1 SV=4 | 19 | 5 | 6 | 5 | 382 | 40.4 | 18.8 | 157710.2979 |
| 0 | Q9Y281 | Cofilin-2 OS=Homo sapiens OX=9606 GN=CFL2 PE=1 SV=1 | 37 | 5 | 6 | 1 | 166 | 18.7 | 18.78 | |
| 0 | O43809 | Cleavage and polyadenylation specificity factor subunit 5 OS=Homo sapiens OX=9606 GN=NUDT21 PE=1 SV=1 | 21 | 3 | 5 | 3 | 227 | 26.2 | 18.65 | 390928.7148 |
| 0 | P68363 | Tubulin alpha-1B chain OS=Homo sapiens OX=9606 GN=TUBA1B PE=1 SV=1 | 19 | 6 | 6 | 6 | 451 | 50.1 | 18.62 | 222448.4043 |
| 0 | P50914 | 60S ribosomal protein L14 OS=Homo sapiens OX=9606 GN=RPL14 PE=1 SV=4 | 28 | 6 | 8 | 6 | 215 | 23.4 | 18.22 | 386349.0566 |
| 0 | P47914 | 60S ribosomal protein L29 OS=Homo sapiens OX=9606 GN=RPL29 PE=1 SV=2 | 26 | 6 | 8 | 6 | 159 | 17.7 | 18.18 | 965186.3955 |
| 0 | P46776 | 60S ribosomal protein L27a OS=Homo sapiens OX=9606 GN=RPL27A PE=1 SV=2 | 37 | 4 | 6 | 4 | 148 | 16.6 | 18 | 175293.7715 |
| 0 | P05386 | 60S acidic ribosomal protein P1 OS=Homo sapiens OX=9606 GN=RPLP1 PE=1 SV=1 | 56 | 3 | 4 | 3 | 114 | 11.5 | 17.82 | 303510.75 |
| 0 | P62913 | 60S ribosomal protein L11 OS=Homo sapiens OX=9606 GN=RPL11 PE=1 SV=2 | 34 | 6 | 6 | 6 | 178 | 20.2 | 17.67 | 263866.5723 |
| 0 | P23246 | Splicing factor, proline- and glutamine-rich OS=Homo sapiens OX=9606 GN=SFPQ PE=1 SV=2 | 9 | 4 | 6 | 4 | 707 | 76.1 | 17.58 | 75134.42188 |
| 0 | Q9C005 | Protein dpy-30 homolog OS=Homo sapiens OX=9606 GN=DPY30 PE=1 SV=1 | 71 | 5 | 6 | 5 | 99 | 11.2 | 17.42 | 228795.9395 |
| 0 | O75643 | U5 small nuclear ribonucleoprotein 200 kDa helicase OS=Homo sapiens OX=9606 GN=SNRNP200 PE=1 SV=2 | 4 | 5 | 5 | 5 | 2136 | 244.4 | 17.3 | 121369.3145 |
| 0 | Q8N163 | Cell cycle and apoptosis regulator protein 2 OS=Homo sapiens OX=9606 GN=CCAR2 PE=1 SV=2 | 12 | 5 | 5 | 5 | 923 | 102.8 | 17.27 | 121774.4023 |
| 0 | E9PRG8 | Uncharacterized protein C11orf98 OS=Homo sapiens OX=9606 GN=C11orf98 PE=4 SV=2 | 36 | 5 | 5 | 5 | 123 | 14.2 | 17.24 | 468575.8594 |
| 0 | Q09028 | Histone-binding protein RBBP4 OS=Homo sapiens OX=9606 GN=RBBP4 PE=1 SV=3 | 15 | 4 | 6 | 2 | 425 | 47.6 | 16.81 | 126499.4404 |
| 0 | Q13724 | Mannosyl-oligosaccharide glucosidase OS=Homo sapiens OX=9606 GN=MOGS PE=1 SV=5 | 11 | 5 | 8 | 5 | 837 | 91.9 | 16.81 | 85845.7793 |
| 0 | Q8WUD4 | Coiled-coil domain-containing protein 12 OS=Homo sapiens OX=9606 GN=CCDC12 PE=1 SV=1 | 40 | 4 | 5 | 4 | 166 | 19.2 | 16.76 | 113539.5137 |
| 0 | Q92499 | ATP-dependent RNA helicase DDX1 OS=Homo sapiens OX=9606 GN=DDX1 PE=1 SV=2 | 17 | 7 | 7 | 7 | 740 | 82.4 | 16.7 | 116868.6074 |
| 0 | O60506 | Heterogeneous nuclear ribonucleoprotein Q OS=Homo sapiens OX=9606 GN=SYNCRIP PE=1 SV=2 | 11 | 6 | 6 | 6 | 623 | 69.6 | 16.4 | 82243.90039 |
| 0 | P43243 | Matrin-3 OS=Homo sapiens OX=9606 GN=MATR3 PE=1 SV=2 | 13 | 4 | 5 | 4 | 847 | 94.6 | 16.22 | 45455.08984 |
| 0 | Q9BWJ5 | Splicing factor 3B subunit 5 OS=Homo sapiens OX=9606 GN=SF3B5 PE=1 SV=1 | 60 | 3 | 5 | 3 | 86 | 10.1 | 16.19 | 237263.293 |
| 0 | Q9HCN8 | Stromal cell-derived factor 2-like protein 1 OS=Homo sapiens OX=9606 GN=SDF2L1 PE=1 SV=2 | 26 | 3 | 4 | 3 | 221 | 23.6 | 16.14 | |
| 0 | P31689 | DnaJ homolog subfamily A member 1 OS=Homo sapiens OX=9606 GN=DNAJA1 PE=1 SV=2 | 17 | 3 | 5 | 3 | 397 | 44.8 | 16.08 | 58857.27246 |
| 0 | Q16629 | Serine/arginine-rich splicing factor 7 OS=Homo sapiens OX=9606 GN=SRSF7 PE=1 SV=1 | 20 | 4 | 5 | 3 | 238 | 27.4 | 15.85 | 114040.418 |
| 0 | P62987 | Ubiquitin-60S ribosomal protein L40 OS=Homo sapiens OX=9606 GN=UBA52 PE=1 SV=2 | 32 | 4 | 6 | 1 | 128 | 14.7 | 15.81 | |
| 0 | P15880 | 40S ribosomal protein S2 OS=Homo sapiens OX=9606 GN=RPS2 PE=1 SV=2 | 17 | 5 | 6 | 5 | 293 | 31.3 | 15.78 | 125126.5068 |
| 0 | P07305 | Histone H1.0 OS=Homo sapiens OX=9606 GN=H1-0 PE=1 SV=3 | 21 | 4 | 5 | 4 | 194 | 20.9 | 15.74 | 167031.7656 |
| 0 | P30050 | 60S ribosomal protein L12 OS=Homo sapiens OX=9606 GN=RPL12 PE=1 SV=1 | 50 | 6 | 6 | 6 | 165 | 17.8 | 15.55 | 258252.7324 |
| 0 | Q99459 | Cell division cycle 5-like protein OS=Homo sapiens OX=9606 GN=CDC5L PE=1 SV=2 | 13 | 4 | 5 | 4 | 802 | 92.2 | 15.46 | 83790.00195 |
| 0 | Q16531 | DNA damage-binding protein 1 OS=Homo sapiens OX=9606 GN=DDB1 PE=1 SV=1 | 6 | 4 | 4 | 4 | 1140 | 126.9 | 15.32 | 30643.21191 |
| 0 | P60842 | Eukaryotic initiation factor 4A-I OS=Homo sapiens OX=9606 GN=EIF4A1 PE=1 SV=1 | 9 | 3 | 5 | 1 | 406 | 46.1 | 15.13 | 19229.31445 |
| 0 | P52298 | Nuclear cap-binding protein subunit 2 OS=Homo sapiens OX=9606 GN=NCBP2 PE=1 SV=1 | 35 | 5 | 5 | 5 | 156 | 18 | 14.93 | 120316.876 |
| 0 | Q13247 | Serine/arginine-rich splicing factor 6 OS=Homo sapiens OX=9606 GN=SRSF6 PE=1 SV=2 | 17 | 6 | 6 | 4 | 344 | 39.6 | 14.87 | 294396.9375 |
| 0 | O15126 | Secretory carrier-associated membrane protein 1 OS=Homo sapiens OX=9606 GN=SCAMP1 PE=1 SV=2 | 28 | 5 | 5 | 5 | 338 | 37.9 | 14.86 | 18009.7793 |
| 0 | Q92804 | TATA-binding protein-associated factor 2N OS=Homo sapiens OX=9606 GN=TAF15 PE=1 SV=1 | 7 | 3 | 5 | 2 | 592 | 61.8 | 14.82 | 48306.23047 |
| 0 | P46459 | Vesicle-fusing ATPase OS=Homo sapiens OX=9606 GN=NSF PE=1 SV=3 | 6 | 4 | 5 | 4 | 744 | 82.5 | 14.74 | 120691.7754 |

| | | | | | | | | | | |
|---|---|---|---|---|---|---|---|---|---|---|
| 0 | P43487 | Ran-specific GTPase-activating protein OS=Homo sapiens OX=9606 GN=RANBP1 PE=1 SV=1 | 28 | 4 | 4 | 4 | 201 | 23.3 | 14.67 | 68777.51758 |
| 0 | Q9Y266 | Nuclear migration protein nudC OS=Homo sapiens OX=9606 GN=NUDC PE=1 SV=1 | 16 | 4 | 5 | 4 | 331 | 38.2 | 14.65 | 125567.499 |
| 0 | C9JLW8 | Mapk-regulated corepressor-interacting protein 1 OS=Homo sapiens OX=9606 GN=MCRIP1 PE=1 SV=1 | 59 | 5 | 5 | 5 | 97 | 10.9 | 14.63 | 125016.709 |
| 0 | P42766 | 60S ribosomal protein L35 OS=Homo sapiens OX=9606 GN=RPL35 PE=1 SV=2 | 50 | 6 | 8 | 6 | 123 | 14.5 | 14.54 | 968891.4268 |
| 0 | P62136 | Serine/threonine-protein phosphatase PP1-alpha catalytic subunit OS=Homo sapiens OX=9606 GN=PPP1CA PE=1 SV=1 | 25 | 6 | 6 | 4 | 330 | 37.5 | 14.51 | 76565.94434 |
| 0 | P62851 | 40S ribosomal protein S25 OS=Homo sapiens OX=9606 GN=RPS25 PE=1 SV=1 | 42 | 6 | 6 | 6 | 125 | 13.7 | 14.17 | 566331.3398 |
| 0 | Q9Y4Y9 | U6 snRNA-associated Sm-like protein LSm5 OS=Homo sapiens OX=9606 GN=LSM5 PE=1 SV=3 | 57 | 3 | 6 | 3 | 91 | 9.9 | 14.13 | 43145.03711 |
| 0 | O75934 | Pre-mRNA-splicing factor SPF27 OS=Homo sapiens OX=9606 GN=BCAS2 PE=1 SV=1 | 24 | 3 | 5 | 3 | 225 | 26.1 | 14.11 | 59426.9668 |
| 0 | P61244 | Protein max OS=Homo sapiens OX=9606 GN=MAX PE=1 SV=1 | 23 | 2 | 3 | 2 | 160 | 18.3 | 13.74 | 48108.63086 |
| 0 | Q09161 | Nuclear cap-binding protein subunit 1 OS=Homo sapiens OX=9606 GN=NCBP1 PE=1 SV=1 | 5 | 2 | 4 | 2 | 790 | 91.8 | 13.58 | 34632.24609 |
| 0 | P47755 | F-actin-capping protein subunit alpha-2 OS=Homo sapiens OX=9606 GN=CAPZA2 PE=1 SV=3 | 15 | 5 | 5 | 1 | 286 | 32.9 | 13.52 | 34494.15234 |
| 0 | P02786 | Transferrin receptor protein 1 OS=Homo sapiens OX=9606 GN=TFRC PE=1 SV=2 | 8 | 4 | 4 | 4 | 760 | 84.8 | 13.06 | 139460.5996 |
| 0 | Q9H4A5 | Golgi phosphoprotein 3-like OS=Homo sapiens OX=9606 GN=GOLPH3L PE=1 SV=1 | 20 | 2 | 3 | 2 | 285 | 32.7 | 12.94 | 110646.9922 |
| 0 | Q15637 | Splicing factor 1 OS=Homo sapiens OX=9606 GN=SF1 PE=1 SV=4 | 8 | 3 | 4 | 3 | 639 | 68.3 | 12.88 | 50572.61328 |
| 0 | Q12874 | Splicing factor 3A subunit 3 OS=Homo sapiens OX=9606 GN=SF3A3 PE=1 SV=1 | 9 | 4 | 4 | 4 | 501 | 58.8 | 12.77 | 28162.92383 |
| 0 | P63027 | Vesicle-associated membrane protein 2 OS=Homo sapiens OX=9606 GN=VAMP2 PE=1 SV=3 | 24 | 3 | 4 | 1 | 116 | 12.7 | 12.69 | 30390.45117 |
| 0 | P38646 | Stress-70 protein, mitochondrial OS=Homo sapiens OX=9606 GN=HSPA9 PE=1 SV=2 | 9 | 4 | 6 | 4 | 679 | 73.6 | 12.61 | 41886.18262 |
| 0 | P31946 | 14-3-3 protein beta/alpha OS=Homo sapiens OX=9606 GN=YWHAB PE=1 SV=3 | 12 | 2 | 3 | 1 | 246 | 28.1 | 12.59 | |
| 0 | Q71UI9 | Histone H2A.V OS=Homo sapiens OX=9606 GN=H2AZ2 PE=1 SV=3 | 31 | 4 | 5 | 2 | 128 | 13.5 | 12.44 | 434536.2539 |
| 0 | Q01105 | Protein SET OS=Homo sapiens OX=9606 GN=SET PE=1 SV=3 | 19 | 3 | 4 | 3 | 290 | 33.5 | 12.25 | 84237.08496 |
| 0 | Q9BWD1 | Acetyl-CoA acetyltransferase, cytosolic OS=Homo sapiens OX=9606 GN=ACAT2 PE=1 SV=2 | 17 | 3 | 3 | 3 | 397 | 41.3 | 12.05 | 95154.92188 |
| 0 | P35268 | 60S ribosomal protein L22 OS=Homo sapiens OX=9606 GN=RPL22 PE=1 SV=2 | 40 | 3 | 3 | 3 | 128 | 14.8 | 11.83 | 197452.5039 |
| 0 | P22392 | Nucleoside diphosphate kinase B OS=Homo sapiens OX=9606 GN=NME2 PE=1 SV=1 | 38 | 4 | 4 | 4 | 152 | 17.3 | 11.81 | 245980.5566 |
| 0 | Q9H0D6 | 5'-3' exoribonuclease 2 OS=Homo sapiens OX=9606 GN=XRN2 PE=1 SV=1 | 8 | 4 | 6 | 4 | 950 | 108.5 | 11.46 | 821366.7441 |
| 0 | P55735 | Protein SEC13 homolog OS=Homo sapiens OX=9606 GN=SEC13 PE=1 SV=3 | 19 | 3 | 3 | 3 | 322 | 35.5 | 11.17 | 161884.374 |
| 0 | P62244 | 40S ribosomal protein S15a OS=Homo sapiens OX=9606 GN=RPS15A PE=1 SV=2 | 24 | 3 | 4 | 3 | 130 | 14.8 | 11.13 | 219282.5117 |
| 0 | P61513 | 60S ribosomal protein L37a OS=Homo sapiens OX=9606 GN=RPL37A PE=1 SV=2 | 50 | 4 | 4 | 4 | 92 | 10.3 | 11.05 | 489002.7539 |
| 0 | Q8N1N4 | Keratin, type II cytoskeletal 78 OS=Homo sapiens OX=9606 GN=KRT78 PE=1 SV=2 | 6 | 4 | 4 | 1 | 520 | 56.8 | 11 | |
| 0 | P42677 | 40S ribosomal protein S27 OS=Homo sapiens OX=9606 GN=RPS27 PE=1 SV=3 | 40 | 4 | 4 | 2 | 84 | 9.5 | 10.98 | 311859.9121 |
| 0 | Q15836 | Vesicle-associated membrane protein 3 OS=Homo sapiens OX=9606 GN=VAMP3 PE=1 SV=3 | 29 | 3 | 4 | 1 | 100 | 11.3 | 10.96 | |
| 0 | Q4VCS5 | Angiomotin OS=Homo sapiens OX=9606 GN=AMOT PE=1 SV=1 | 5 | 3 | 3 | 3 | 1084 | 118 | 10.96 | 18398.47852 |
| 0 | Q9BVG4 | Protein PBDC1 OS=Homo sapiens OX=9606 GN=PBDC1 PE=1 SV=1 | 17 | 3 | 3 | 3 | 233 | 26 | 10.87 | 19806.44531 |
| 0 | P61626 | Lysozyme C OS=Homo sapiens OX=9606 GN=LYZ PE=1 SV=1 | 34 | 5 | 5 | 5 | 148 | 16.5 | 10.84 | 45292.7373 |
| 0 | Q5BKY9 | Protein FAM133B OS=Homo sapiens OX=9606 GN=FAM133B PE=1 SV=1 | 10 | 3 | 4 | 3 | 247 | 28.4 | 10.8 | 196311.9453 |
| 0 | Q5T749 | Keratinocyte proline-rich protein OS=Homo sapiens OX=9606 GN=KPRP PE=1 SV=1 | 8 | 3 | 4 | 3 | 579 | 64.1 | 10.67 | 111493.4531 |
| 0 | P06733 | Alpha-enolase OS=Homo sapiens OX=9606 GN=ENO1 PE=1 SV=2 | 12 | 3 | 3 | 3 | 434 | 47.1 | 10.47 | 41409.87695 |
| 0 | Q9H2H8 | Peptidyl-prolyl cis-trans isomerase-like 3 OS=Homo sapiens OX=9606 GN=PPIL3 PE=1 SV=1 | 34 | 3 | 4 | 3 | 161 | 18.1 | 10.25 | 73683.04102 |
| 0 | O60232 | Protein ZNRD2 OS=Homo sapiens OX=9606 GN=ZNRD2 PE=1 SV=1 | 20 | 2 | 3 | 2 | 199 | 21.5 | 10.21 | 41138.46875 |
| 0 | P35637 | RNA-binding protein FUS OS=Homo sapiens OX=9606 GN=FUS PE=1 SV=1 | 7 | 2 | 3 | 1 | 526 | 53.4 | 10.06 | |
| 0 | P00492 | Hypoxanthine-guanine phosphoribosyltransferase OS=Homo sapiens OX=9606 GN=HPRT1 PE=1 SV=2 | 26 | 3 | 3 | 3 | 218 | 24.6 | 10.02 | 45699.21094 |
| 0 | Q96CN7 | Isochorismatase domain-containing protein 1 OS=Homo sapiens OX=9606 GN=ISOC1 PE=1 SV=3 | 16 | 3 | 4 | 3 | 298 | 32.2 | 9.95 | 122117.0972 |
| 0 | Q14103 | Heterogeneous nuclear ribonucleoprotein D0 OS=Homo sapiens OX=9606 GN=HNRNPD PE=1 SV=1 | 11 | 3 | 3 | 2 | 355 | 38.4 | 9.94 | 57944.70313 |
| 0 | Q16576 | Histone-binding protein RBBP7 OS=Homo sapiens OX=9606 GN=RBBP7 PE=1 SV=1 | 8 | 3 | 4 | 1 | 425 | 47.8 | 9.94 | |
| 0 | P46777 | 60S ribosomal protein L5 OS=Homo sapiens OX=9606 GN=RPL5 PE=1 SV=3 | 15 | 3 | 4 | 3 | 297 | 34.3 | 9.88 | 87587.60156 |
| 0 | Q13185 | Chromobox protein homolog 3 OS=Homo sapiens OX=9606 GN=CBX3 PE=1 SV=4 | 20 | 3 | 3 | 3 | 183 | 20.8 | 9.87 | 27889.88477 |
| 0 | Q9UQ35 | Serine/arginine repetitive matrix protein 2 OS=Homo sapiens OX=9606 GN=SRRM2 PE=1 SV=2 | 2 | 4 | 4 | 4 | 2752 | 299.4 | 9.81 | 110456.6641 |
| 0 | P06730 | Eukaryotic translation initiation factor 4E OS=Homo sapiens OX=9606 GN=EIF4E PE=1 SV=2 | 39 | 5 | 5 | 5 | 217 | 25.1 | 9.76 | 216544.1465 |
| 0 | P39023 | 60S ribosomal protein L3 OS=Homo sapiens OX=9606 GN=RPL3 PE=1 SV=2 | 9 | 3 | 3 | 3 | 403 | 46.1 | 9.75 | 57425.86035 |
| 0 | P61981 | 14-3-3 protein gamma OS=Homo sapiens OX=9606 GN=YWHAG PE=1 SV=2 | 17 | 3 | 3 | 2 | 247 | 28.3 | 9.71 | 132146.3184 |
| 0 | P05388 | 60S acidic ribosomal protein P0 OS=Homo sapiens OX=9606 GN=RPLP0 PE=1 SV=1 | 15 | 3 | 3 | 3 | 317 | 34.3 | 9.57 | 34454.80469 |
| 0 | P53999 | Activated RNA polymerase II transcriptional coactivator p15 OS=Homo sapiens OX=9606 GN=SUB1 PE=1 SV=3 | 31 | 2 | 3 | 2 | 127 | 14.4 | 9.49 | 119829.9375 |
| 0 | P62241 | 40S ribosomal protein S8 OS=Homo sapiens OX=9606 GN=RPS8 PE=1 SV=2 | 18 | 3 | 3 | 3 | 208 | 24.2 | 9.39 | 116302.1719 |
| 0 | Q6IS14 | Eukaryotic translation initiation factor 5A-1-like OS=Homo sapiens OX=9606 GN=EIF5AL1 PE=2 SV=2 | 25 | 2 | 3 | 2 | 154 | 16.8 | 9.37 | 104435.9648 |
| 0 | P62258 | 14-3-3 protein epsilon OS=Homo sapiens OX=9606 GN=YWHAE PE=1 SV=1 | 20 | 3 | 3 | 3 | 255 | 29.2 | 9.36 | 249259.3867 |
| 0 | Q9UHB6 | LIM domain and actin-binding protein 1 OS=Homo sapiens OX=9606 GN=LIMA1 PE=1 SV=1 | 7 | 3 | 5 | 3 | 759 | 85.2 | 9.16 | 35631.30859 |
| 0 | P62861 | 40S ribosomal protein S30 OS=Homo sapiens OX=9606 GN=FAU PE=1 SV=1 | 34 | 4 | 5 | 4 | 59 | 6.6 | 9.14 | 1496041.73 |
| 0 | P81605 | Dermcidin OS=Homo sapiens OX=9606 GN=DCD PE=1 SV=2 | 25 | 2 | 3 | 2 | 110 | 11.3 | 8.98 | 131719.8926 |
| 0 | Q9NPA8 | Transcription and mRNA export factor ENY2 OS=Homo sapiens OX=9606 GN=ENY2 PE=1 SV=1 | 26 | 2 | 3 | 2 | 101 | 11.5 | 8.97 | 46216.96289 |
| 0 | O76021 | Ribosomal L1 domain-containing protein 1 OS=Homo sapiens OX=9606 GN=RSL1D1 PE=1 SV=3 | 9 | 4 | 4 | 4 | 490 | 54.9 | 8.96 | 66432.55371 |
| 0 | P49458 | Signal recognition particle 9 kDa protein OS=Homo sapiens OX=9606 GN=SRP9 PE=1 SV=2 | 30 | 2 | 3 | 2 | 86 | 10.1 | 8.86 | 141983.7266 |
| 0 | Q03111 | Protein ENL OS=Homo sapiens OX=9606 GN=MLLT1 PE=1 SV=2 | 9 | 2 | 2 | 2 | 559 | 62 | 8.8 | 44715.45313 |
| 0 | P33316 | Deoxyuridine 5'-triphosphate nucleotidohydrolase, mitochondrial OS=Homo sapiens OX=9606 GN=DUT PE=1 SV=4 | 16 | 3 | 3 | 3 | 252 | 26.5 | 8.77 | 204778.1445 |
| 0 | P63220 | 40S ribosomal protein S21 OS=Homo sapiens OX=9606 GN=RPS21 PE=1 SV=1 | 45 | 3 | 3 | 3 | 83 | 9.1 | 8.63 | 108377.5098 |
| 0 | O00571 | ATP-dependent RNA helicase DDX3X OS=Homo sapiens OX=9606 GN=DDX3X PE=1 SV=3 | 9 | 4 | 4 | 4 | 662 | 73.2 | 8.6 | 97315.66406 |

| | | | | | | | | | | |
|---|---|---|---|---|---|---|---|---|---|---|
| 0 | P36873 | Serine/threonine-protein phosphatase PP1-gamma catalytic subunit OS=Homo sapiens OX=9606 GN=PPP1CC PE=1 SV=1 | 14 | 3 | 3 | 1 | 323 | 37 | 8.59 | |
| 0 | Q13595 | Transformer-2 protein homolog alpha OS=Homo sapiens OX=9606 GN=TRA2A PE=1 SV=1 | 11 | 3 | 3 | 3 | 282 | 32.7 | 8.59 | 62032.55957 |
| 0 | Q15428 | Splicing factor 3A subunit 2 OS=Homo sapiens OX=9606 GN=SF3A2 PE=1 SV=2 | 12 | 4 | 4 | 4 | 464 | 49.2 | 8.52 | 150155.1553 |
| 0 | P61964 | WD repeat-containing protein 5 OS=Homo sapiens OX=9606 GN=WDR5 PE=1 SV=1 | 7 | 2 | 3 | 2 | 334 | 36.6 | 8.51 | 34882.40918 |
| 0 | Q13242 | Serine/arginine-rich splicing factor 9 OS=Homo sapiens OX=9606 GN=SRSF9 PE=1 SV=1 | 18 | 4 | 5 | 3 | 221 | 25.5 | 8.41 | 40713.95117 |
| 0 | Q01844 | RNA-binding protein EWS OS=Homo sapiens OX=9606 GN=EWSR1 PE=1 SV=1 | 3 | 1 | 2 | 1 | 656 | 68.4 | 8.38 | 40202.66797 |
| 0 | P18077 | 60S ribosomal protein L35a OS=Homo sapiens OX=9606 GN=RPL35A PE=1 SV=2 | 24 | 4 | 4 | 4 | 110 | 12.5 | 8.37 | 127789.7207 |
| 0 | Q86X55 | Histone-arginine methyltransferase CARM1 OS=Homo sapiens OX=9606 GN=CARM1 PE=1 SV=3 | 7 | 3 | 3 | 3 | 608 | 65.8 | 8.37 | 32020.95215 |
| 0 | O14744 | Protein arginine N-methyltransferase 5 OS=Homo sapiens OX=9606 GN=PRMT5 PE=1 SV=4 | 7 | 3 | 3 | 3 | 637 | 72.6 | 8.27 | 55573.09375 |
| 0 | Q86SE5 | RNA-binding Raly-like protein OS=Homo sapiens OX=9606 GN=RALYL PE=1 SV=2 | 8 | 3 | 3 | 1 | 291 | 32.3 | 8.23 | 30887.15625 |
| 0 | Q9P258 | Protein RCC2 OS=Homo sapiens OX=9606 GN=RCC2 PE=1 SV=2 | 3 | 1 | 3 | 1 | 522 | 56 | 8.19 | 18373.75 |
| 0 | P62314 | Small nuclear ribonucleoprotein Sm D1 OS=Homo sapiens OX=9606 GN=SNRPD1 PE=1 SV=1 | 34 | 3 | 4 | 3 | 119 | 13.3 | 8.15 | 260230.4712 |
| 0 | Q15437 | Protein transport protein Sec23B OS=Homo sapiens OX=9606 GN=SEC23B PE=1 SV=2 | 5 | 3 | 3 | 1 | 767 | 86.4 | 8.13 | |
| 0 | Q7RTV0 | PHD finger-like domain-containing protein 5A OS=Homo sapiens OX=9606 GN=PHF5A PE=1 SV=1 | 25 | 2 | 2 | 2 | 110 | 12.4 | 8.03 | 91409.48047 |
| 0 | P14923 | Junction plakoglobin OS=Homo sapiens OX=9606 GN=JUP PE=1 SV=3 | 4 | 2 | 2 | 2 | 745 | 81.7 | 7.99 | |
| 0 | Q3MHD2 | Protein LSM12 homolog OS=Homo sapiens OX=9606 GN=LSM12 PE=1 SV=2 | 18 | 3 | 3 | 3 | 195 | 21.7 | 7.95 | 91262.60938 |
| 0 | Q9UM54 | Unconventional myosin-VI OS=Homo sapiens OX=9606 GN=MYO6 PE=1 SV=4 | 3 | 2 | 3 | 2 | 1294 | 149.6 | 7.94 | 40472.65723 |
| 0 | Q71UM5 | 40S ribosomal protein S27-like OS=Homo sapiens OX=9606 GN=RPS27L PE=1 SV=3 | 40 | 3 | 3 | 1 | 84 | 9.5 | 7.93 | 30156.49414 |
| 0 | Q13283 | Ras GTPase-activating protein-binding protein 1 OS=Homo sapiens OX=9606 GN=G3BP1 PE=1 SV=1 | 8 | 2 | 3 | 2 | 466 | 52.1 | 7.91 | |
| 0 | Q9Y6M1 | Insulin-like growth factor 2 mRNA-binding protein 2 OS=Homo sapiens OX=9606 GN=IGF2BP2 PE=1 SV=2 | 6 | 4 | 4 | 1 | 599 | 66.1 | 7.82 | |
| 0 | P35244 | Replication protein A 14 kDa subunit OS=Homo sapiens OX=9606 GN=RPA3 PE=1 SV=1 | 32 | 2 | 3 | 2 | 121 | 13.6 | 7.79 | 36737.97266 |
| 0 | Q6RW13 | Type-1 angiotensin II receptor-associated protein OS=Homo sapiens OX=9606 GN=AGTRAP PE=1 SV=1 | 25 | 2 | 2 | 2 | 159 | 17.4 | 7.74 | 65971.78125 |
| 0 | Q9C037 | E3 ubiquitin-protein ligase TRIM4 OS=Homo sapiens OX=9606 GN=TRIM4 PE=1 SV=2 | 8 | 3 | 3 | 3 | 500 | 57.4 | 7.65 | 91003.41797 |
| 0 | Q9BY49 | Peroxisomal trans-2-enoyl-CoA reductase OS=Homo sapiens OX=9606 GN=PECR PE=1 SV=2 | 12 | 2 | 2 | 2 | 303 | 32.5 | 7.59 | |
| 0 | Q16342 | Programmed cell death protein 2 OS=Homo sapiens OX=9606 GN=PDCD2 PE=1 SV=2 | 14 | 2 | 2 | 2 | 344 | 38.6 | 7.44 | 43433.03125 |
| 0 | P83881 | 60S ribosomal protein L36a OS=Homo sapiens OX=9606 GN=RPL36A PE=1 SV=2 | 28 | 4 | 4 | 4 | 106 | 12.4 | 7.43 | 183360.7617 |
| 0 | P51149 | Ras-related protein Rab-7a OS=Homo sapiens OX=9606 GN=RAB7A PE=1 SV=1 | 28 | 4 | 4 | 4 | 207 | 23.5 | 7.41 | 76777.22266 |
| 0 | P62820 | Ras-related protein Rab-1A OS=Homo sapiens OX=9606 GN=RAB1A PE=1 SV=3 | 13 | 2 | 2 | 1 | 205 | 22.7 | 7.39 | 21637.71484 |
| 0 | P20930 | Filaggrin OS=Homo sapiens OX=9606 GN=FLG PE=1 SV=3 | 1 | 2 | 2 | 2 | 4061 | 434.9 | 7.37 | 4568.587891 |
| 0 | Q53F19 | Nuclear cap-binding protein subunit 3 OS=Homo sapiens OX=9606 GN=NCBP3 PE=1 SV=2 | 6 | 3 | 3 | 3 | 620 | 70.5 | 7.36 | |
| 0 | P18124 | 60S ribosomal protein L7 OS=Homo sapiens OX=9606 GN=RPL7 PE=1 SV=1 | 12 | 2 | 2 | 2 | 248 | 29.2 | 7.3 | 57044.39844 |
| 0 | Q99959 | Plakophilin-2 OS=Homo sapiens OX=9606 GN=PKP2 PE=1 SV=2 | 4 | 2 | 2 | 2 | 881 | 97.4 | 7.18 | |
| 0 | P63208 | S-phase kinase-associated protein 1 OS=Homo sapiens OX=9606 GN=SKP1 PE=1 SV=2 | 23 | 2 | 3 | 2 | 163 | 18.6 | 7.11 | 195402.3203 |
| 0 | P22061 | Protein-L-isoaspartate(D-aspartate) O-methyltransferase OS=Homo sapiens OX=9606 GN=PCMT1 PE=1 SV=4 | 26 | 2 | 3 | 2 | 227 | 24.6 | 7.11 | 119434.6055 |
| 0 | P31151 | Protein S100-A7 OS=Homo sapiens OX=9606 GN=S100A7 PE=1 SV=4 | 36 | 3 | 3 | 3 | 101 | 11.5 | 7.08 | 71655.66797 |
| 0 | P06702 | Protein S100-A9 OS=Homo sapiens OX=9606 GN=S100A9 PE=1 SV=1 | 25 | 2 | 2 | 2 | 114 | 13.2 | 6.97 | 116164.6133 |
| 0 | Q96HQ2 | CDKN2AIP N-terminal-like protein OS=Homo sapiens OX=9606 GN=CDKN2AIPNL PE=1 SV=1 | 24 | 2 | 2 | 2 | 116 | 13.2 | 6.94 | 12395.20313 |
| 0 | P36578 | 60S ribosomal protein L4 OS=Homo sapiens OX=9606 GN=RPL4 PE=1 SV=5 | 7 | 2 | 2 | 2 | 427 | 47.7 | 6.88 | 35019.53906 |
| 0 | O60925 | Prefoldin subunit 1 OS=Homo sapiens OX=9606 GN=PFDN1 PE=1 SV=2 | 31 | 3 | 3 | 3 | 122 | 14.2 | 6.81 | 56855.92285 |
| 0 | Q8N1G0 | Zinc finger protein 687 OS=Homo sapiens OX=9606 GN=ZNF687 PE=1 SV=1 | 5 | 2 | 2 | 2 | 1237 | 129.4 | 6.77 | 12304.47168 |
| 0 | P51572 | B-cell receptor-associated protein 31 OS=Homo sapiens OX=9606 GN=BCAP31 PE=1 SV=3 | 9 | 2 | 2 | 2 | 246 | 28 | 6.76 | 122290.457 |
| 0 | P56270 | Myc-associated zinc finger protein OS=Homo sapiens OX=9606 GN=MAZ PE=1 SV=1 | 5 | 2 | 2 | 2 | 477 | 48.6 | 6.75 | 51503.35938 |
| 0 | P27635 | 60S ribosomal protein L10 OS=Homo sapiens OX=9606 GN=RPL10 PE=1 SV=4 | 18 | 2 | 2 | 2 | 214 | 24.6 | 6.7 | |
| 0 | P34932 | Heat shock 70 kDa protein 4 OS=Homo sapiens OX=9606 GN=HSPA4 PE=1 SV=4 | 4 | 2 | 2 | 2 | 840 | 94.3 | 6.67 | |
| 0 | P33992 | DNA replication licensing factor MCM5 OS=Homo sapiens OX=9606 GN=MCM5 PE=1 SV=5 | 4 | 2 | 2 | 2 | 734 | 82.2 | 6.63 | 47486.58789 |
| 0 | Q6UN15 | Pre-mRNA 3'-end-processing factor FIP1 OS=Homo sapiens OX=9606 GN=FIP1L1 PE=1 SV=1 | 6 | 2 | 2 | 2 | 594 | 66.5 | 6.6 | |
| 0 | P04080 | Cystatin-B OS=Homo sapiens OX=9606 GN=CSTB PE=1 SV=2 | 34 | 2 | 2 | 2 | 98 | 11.1 | 6.42 | 55836.47461 |
| 0 | Q9Y3X0 | Coiled-coil domain-containing protein 9 OS=Homo sapiens OX=9606 GN=CCDC9 PE=1 SV=1 | 5 | 2 | 2 | 2 | 531 | 59.7 | 6.37 | 10481.63574 |
| 0 | Q08170 | Serine/arginine-rich splicing factor 4 OS=Homo sapiens OX=9606 GN=SRSF4 PE=1 SV=2 | 5 | 3 | 3 | 1 | 494 | 56.6 | 6.37 | 36560.09375 |
| 0 | P04062 | Lysosomal acid glucosylceramidase OS=Homo sapiens OX=9606 GN=GBA PE=1 SV=3 | 4 | 1 | 2 | 1 | 536 | 59.7 | 6.35 | 25018.62695 |
| 0 | Q15369 | Elongin-C OS=Homo sapiens OX=9606 GN=ELOC PE=1 SV=1 | 38 | 2 | 2 | 2 | 112 | 12.5 | 6.33 | |
| 0 | O00479 | High mobility group nucleosome-binding domain-containing protein 4 OS=Homo sapiens OX=9606 GN=HMGN4 PE=1 SV=3 | 36 | 2 | 3 | 2 | 90 | 9.5 | 6.33 | 98692.7168 |
| 0 | O60832 | H/ACA ribonucleoprotein complex subunit DKC1 OS=Homo sapiens OX=9606 GN=DKC1 PE=1 SV=3 | 7 | 2 | 2 | 2 | 514 | 57.6 | 6.32 | 29881.98633 |
| 0 | O75494 | Serine/arginine-rich splicing factor 10 OS=Homo sapiens OX=9606 GN=SRSF10 PE=1 SV=1 | 13 | 2 | 2 | 2 | 262 | 31.3 | 6.27 | 95388.39453 |
| 0 | Q13557 | Calcium/calmodulin-dependent protein kinase type II subunit delta OS=Homo sapiens OX=9606 GN=CAMK2D PE=1 SV=3 | 6 | 3 | 3 | 3 | 499 | 56.3 | 6.27 | 82506.41406 |
| 0 | P61421 | V-type proton ATPase subunit d 1 OS=Homo sapiens OX=9606 GN=ATP6V0D1 PE=1 SV=1 | 8 | 2 | 3 | 2 | 351 | 40.3 | 6.26 | 27475.92188 |
| 0 | P60174 | Triosephosphate isomerase OS=Homo sapiens OX=9606 GN=TPI1 PE=1 SV=4 | 10 | 2 | 2 | 2 | 249 | 26.7 | 6.19 | |
| 0 | Q9Y3B4 | Splicing factor 3B subunit 6 OS=Homo sapiens OX=9606 GN=SF3B6 PE=1 SV=1 | 27 | 3 | 3 | 3 | 125 | 14.6 | 6.1 | 108817.4141 |
| 0 | Q01130 | Serine/arginine-rich splicing factor 2 OS=Homo sapiens OX=9606 GN=SRSF2 PE=1 SV=4 | 11 | 1 | 1 | 1 | 221 | 25.5 | 6.01 | 33669.28125 |
| 0 | P49368 | T-complex protein 1 subunit gamma OS=Homo sapiens OX=9606 GN=CCT3 PE=1 SV=4 | 8 | 2 | 2 | 2 | 545 | 60.5 | 5.96 | 40850.61426 |
| 0 | O60573 | Eukaryotic translation initiation factor 4E type 2 OS=Homo sapiens OX=9606 GN=EIF4E2 PE=1 SV=1 | 13 | 2 | 2 | 2 | 245 | 28.3 | 5.93 | 41476.71484 |
| 0 | P17096 | High mobility group protein HMG-I/HMG-Y OS=Homo sapiens OX=9606 GN=HMGA1 PE=1 SV=3 | 38 | 2 | 2 | 2 | 107 | 11.7 | 5.89 | 104392.0098 |
| 0 | Q6P2Q9 | Pre-mRNA-processing-splicing factor 8 OS=Homo sapiens OX=9606 GN=PRPF8 PE=1 SV=2 | 2 | 2 | 2 | 2 | 2335 | 273.4 | 5.85 | 35560.45703 |

| | | | | | | | | | | |
|---|---|---|---|---|---|---|---|---|---|---|
| 0 | Q9H814 | Phosphorylated adapter RNA export protein OS=Homo sapiens OX=9606 GN=PHAX PE=1 SV=1 | 4 | 1 | 2 | 1 | 394 | 44.4 | 5.84 | |
| 0 | P28799 | Progranulin OS=Homo sapiens OX=9606 GN=GRN PE=1 SV=2 | 3 | 1 | 2 | 1 | 593 | 63.5 | 5.81 | 19366.6543 |
| 0 | Q6PKG0 | La-related protein 1 OS=Homo sapiens OX=9606 GN=LARP1 PE=1 SV=2 | 4 | 2 | 2 | 2 | 1096 | 123.4 | 5.81 | |
| 0 | P62306 | Small nuclear ribonucleoprotein F OS=Homo sapiens OX=9606 GN=SNRPF PE=1 SV=1 | 41 | 2 | 2 | 2 | 86 | 9.7 | 5.76 | 115388.3867 |
| 0 | Q9P013 | Spliceosome-associated protein CWC15 homolog OS=Homo sapiens OX=9606 GN=CWC15 PE=1 SV=2 | 11 | 2 | 2 | 2 | 229 | 26.6 | 5.76 | 139686.9453 |
| 0 | P63000 | Ras-related C3 botulinum toxin substrate 1 OS=Homo sapiens OX=9606 GN=RAC1 PE=1 SV=1 | 13 | 2 | 2 | 2 | 192 | 21.4 | 5.69 | 50085.60156 |
| 0 | P32969 | 60S ribosomal protein L9 OS=Homo sapiens OX=9606 GN=RPL9 PE=1 SV=1 | 11 | 1 | 2 | 1 | 192 | 21.9 | 5.67 | |
| 0 | O15212 | Prefoldin subunit 6 OS=Homo sapiens OX=9606 GN=PFDN6 PE=1 SV=1 | 21 | 2 | 2 | 2 | 129 | 14.6 | 5.5 | 49680.67969 |
| 0 | Q9NW07 | Zinc finger protein 358 OS=Homo sapiens OX=9606 GN=ZNF358 PE=1 SV=2 | 4 | 2 | 2 | 2 | 568 | 59.3 | 5.49 | |
| 0 | P09234 | U1 small nuclear ribonucleoprotein C OS=Homo sapiens OX=9606 GN=SNRPC PE=1 SV=1 | 13 | 1 | 2 | 1 | 159 | 17.4 | 5.43 | 45356.49902 |
| 0 | P61163 | Alpha-centractin OS=Homo sapiens OX=9606 GN=ACTR1A PE=1 SV=1 | 6 | 2 | 2 | 2 | 376 | 42.6 | 5.43 | 37133.31445 |
| 0 | Q9BTL3 | RNA guanine-N7 methyltransferase activating subunit OS=Homo sapiens OX=9606 GN=RAMAC PE=1 SV=1 | 26 | 2 | 2 | 2 | 118 | 14.4 | 5.42 | 21934.0957 |
| 0 | Q92928 | Putative Ras-related protein Rab-1C OS=Homo sapiens OX=9606 GN=RAB1C PE=5 SV=2 | 14 | 2 | 2 | 1 | 201 | 22 | 5.4 | 7819.465332 |
| 0 | P62308 | Small nuclear ribonucleoprotein G OS=Homo sapiens OX=9606 GN=SNRPG PE=1 SV=1 | 26 | 2 | 2 | 2 | 76 | 8.5 | 5.37 | 218392.1797 |
| 0 | O43143 | Pre-mRNA-splicing factor ATP-dependent RNA helicase DHX15 OS=Homo sapiens OX=9606 GN=DHX15 PE=1 SV=2 | 1 | 1 | 2 | 1 | 795 | 90.9 | 5.35 | |
| 0 | Q9GZZ1 | N-alpha-acetyltransferase 50 OS=Homo sapiens OX=9606 GN=NAA50 PE=1 SV=1 | 17 | 2 | 2 | 2 | 169 | 19.4 | 5.33 | 34028.36133 |
| 0 | O75607 | Nucleoplasmin-3 OS=Homo sapiens OX=9606 GN=NPM3 PE=1 SV=3 | 10 | 1 | 1 | 1 | 178 | 19.3 | 5.21 | 29403.42188 |
| 0 | P10599 | Thioredoxin OS=Homo sapiens OX=9606 GN=TXN PE=1 SV=3 | 19 | 1 | 2 | 1 | 105 | 11.7 | 5.19 | 42059.36719 |
| 0 | Q8TE02 | Elongator complex protein 5 OS=Homo sapiens OX=9606 GN=ELP5 PE=1 SV=2 | 9 | 1 | 1 | 1 | 316 | 34.8 | 5.18 | 22593.82617 |
| 0 | P55769 | NHP2-like protein 1 OS=Homo sapiens OX=9606 GN=SNU13 PE=1 SV=3 | 19 | 2 | 2 | 2 | 128 | 14.2 | 5.17 | 76761.21094 |
| 0 | Q15427 | Splicing factor 3B subunit 4 OS=Homo sapiens OX=9606 GN=SF3B4 PE=1 SV=1 | 6 | 2 | 3 | 2 | 424 | 44.4 | 5.14 | 20469.19531 |
| 0 | Q9H3K6 | BolA-like protein 2 OS=Homo sapiens OX=9606 GN=BOLA2 PE=1 SV=1 | 19 | 1 | 1 | 1 | 86 | 10.1 | 5.11 | 30187.04102 |
| 0 | O43670 | BUB3-interacting and GLEBS motif-containing protein ZNF207 OS=Homo sapiens OX=9606 GN=ZNF207 PE=1 SV=1 | 3 | 1 | 2 | 1 | 478 | 50.7 | 5.1 | 22116.39844 |
| 0 | P26368 | Splicing factor U2AF 65 kDa subunit OS=Homo sapiens OX=9606 GN=U2AF2 PE=1 SV=4 | 8 | 3 | 3 | 3 | 475 | 53.5 | 5.09 | 60699.08398 |
| 0 | Q5EBL8 | PDZ domain-containing protein 11 OS=Homo sapiens OX=9606 GN=PDZD11 PE=1 SV=2 | 19 | 1 | 2 | 1 | 140 | 16.1 | 5.08 | 19484.82617 |
| 0 | P62633 | Cellular nucleic acid-binding protein OS=Homo sapiens OX=9606 GN=CNBP PE=1 SV=1 | 17 | 2 | 2 | 2 | 177 | 19.5 | 5.05 | 29000.60742 |
| 0 | Q9BT78 | COP9 signalosome complex subunit 4 OS=Homo sapiens OX=9606 GN=COPS4 PE=1 SV=1 | 5 | 2 | 2 | 2 | 406 | 46.2 | 4.99 | 24051.31641 |
| 0 | P55081 | Microfibrillar-associated protein 1 OS=Homo sapiens OX=9606 GN=MFAP1 PE=1 SV=2 | 6 | 1 | 1 | 1 | 439 | 51.9 | 4.98 | |
| 0 | Q96SI1 | BTB/POZ domain-containing protein KCTD15 OS=Homo sapiens OX=9606 GN=KCTD15 PE=1 SV=1 | 7 | 1 | 1 | 1 | 283 | 31.9 | 4.9 | 49532.73047 |
| 0 | P62266 | 40S ribosomal protein S23 OS=Homo sapiens OX=9606 GN=RPS23 PE=1 SV=3 | 15 | 2 | 2 | 2 | 143 | 15.8 | 4.86 | 96879.05273 |
| 0 | Q8TA86 | Retinitis pigmentosa 9 protein OS=Homo sapiens OX=9606 GN=RP9 PE=1 SV=2 | 11 | 2 | 3 | 2 | 221 | 26.1 | 4.84 | 36009.49219 |
| 0 | P49207 | 60S ribosomal protein L34 OS=Homo sapiens OX=9606 GN=RPL34 PE=1 SV=3 | 10 | 1 | 2 | 1 | 117 | 13.3 | 4.83 | 24062.36133 |
| 0 | Q9NVT9 | Armadillo repeat-containing protein 1 OS=Homo sapiens OX=9606 GN=ARMC1 PE=1 SV=1 | 7 | 1 | 1 | 1 | 282 | 31.3 | 4.79 | 20496.91992 |
| 0 | Q15370 | Elongin-B OS=Homo sapiens OX=9606 GN=ELOB PE=1 SV=1 | 43 | 2 | 2 | 2 | 118 | 13.1 | 4.77 | 49518.42188 |
| 0 | P62304 | Small nuclear ribonucleoprotein E OS=Homo sapiens OX=9606 GN=SNRPE PE=1 SV=1 | 29 | 2 | 2 | 2 | 92 | 10.8 | 4.73 | 199299.0313 |
| 0 | Q9UNZ5 | Leydig cell tumor 10 kDa protein homolog OS=Homo sapiens OX=9606 GN=C19orf53 PE=1 SV=1 | 22 | 2 | 3 | 2 | 99 | 10.6 | 4.72 | 32764.76855 |
| 0 | P62873 | Guanine nucleotide-binding protein G(I)/G(S)/G(T) subunit beta-1 OS=Homo sapiens OX=9606 GN=GNB1 PE=1 SV=3 | 6 | 1 | 1 | 1 | 340 | 37.4 | 4.66 | 15429.22754 |
| 0 | Q8N5F7 | NF-kappa-B-activating protein OS=Homo sapiens OX=9606 GN=NKAP PE=1 SV=1 | 7 | 1 | 1 | 1 | 415 | 47.1 | 4.62 | 8623.53125 |
| 0 | Q15393 | Splicing factor 3B subunit 3 OS=Homo sapiens OX=9606 GN=SF3B3 PE=1 SV=4 | 3 | 2 | 2 | 2 | 1217 | 135.5 | 4.59 | 25456.95117 |
| 0 | Q9Y608 | Leucine-rich repeat flightless-interacting protein 2 OS=Homo sapiens OX=9606 GN=LRRFIP2 PE=1 SV=1 | 3 | 2 | 2 | 2 | 721 | 82.1 | 4.58 | 30168.24023 |
| 0 | O00193 | Small acidic protein OS=Homo sapiens OX=9606 GN=SMAP PE=1 SV=1 | 13 | 1 | 1 | 1 | 183 | 20.3 | 4.54 | |
| 0 | P61158 | Actin-related protein 3 OS=Homo sapiens OX=9606 GN=ACTR3 PE=1 SV=3 | 4 | 1 | 1 | 1 | 418 | 47.3 | 4.42 | 36174.28516 |
| 0 | Q8NCA5 | Protein FAM98A OS=Homo sapiens OX=9606 GN=FAM98A PE=1 SV=2 | 2 | 1 | 2 | 1 | 518 | 55.2 | 4.34 | |
| 0 | Q8WXC6 | COP9 signalosome complex subunit 9 OS=Homo sapiens OX=9606 GN=COPS9 PE=1 SV=3 | 35 | 1 | 2 | 1 | 57 | 6.2 | 4.29 | |
| 0 | Q9BUT9 | MAPK regulated corepressor interacting protein 2 OS=Homo sapiens OX=9606 GN=MCRIP2 PE=1 SV=2 | 19 | 1 | 1 | 1 | 160 | 17.8 | 4.29 | |
| 0 | P23526 | Adenosylhomocysteinase OS=Homo sapiens OX=9606 GN=AHCY PE=1 SV=4 | 3 | 1 | 1 | 1 | 432 | 47.7 | 4.27 | |
| 0 | P62854 | 40S ribosomal protein S26 OS=Homo sapiens OX=9606 GN=RPS26 PE=1 SV=3 | 19 | 2 | 2 | 2 | 115 | 13 | 4.25 | 52221.59375 |
| 0 | Q14684 | Ribosomal RNA processing protein 1 homolog B OS=Homo sapiens OX=9606 GN=RRP1B PE=1 SV=3 | 2 | 1 | 2 | 1 | 758 | 84.4 | 4.25 | |
| 0 | Q04837 | Single-stranded DNA-binding protein, mitochondrial OS=Homo sapiens OX=9606 GN=SSBP1 PE=1 SV=1 | 10 | 1 | 1 | 1 | 148 | 17.2 | 4.23 | 23758.17773 |
| 0 | P42285 | Exosome RNA helicase MTR4 OS=Homo sapiens OX=9606 GN=MTREX PE=1 SV=3 | 2 | 1 | 1 | 1 | 1042 | 117.7 | 4.22 | 16628.85742 |
| 0 | Q8N9Q2 | Protein SREK1IP1 OS=Homo sapiens OX=9606 GN=SREK1IP1 PE=1 SV=1 | 16 | 2 | 2 | 2 | 155 | 18.2 | 4.19 | 236756.4063 |
| 0 | P61758 | Prefoldin subunit 3 OS=Homo sapiens OX=9606 GN=VBP1 PE=1 SV=4 | 13 | 1 | 1 | 1 | 197 | 22.6 | 4.17 | |
| 0 | Q10570 | Cleavage and polyadenylation specificity factor subunit 1 OS=Homo sapiens OX=9606 GN=CPSF1 PE=1 SV=2 | 1 | 1 | 1 | 1 | 1443 | 160.8 | 4.03 | 9354.52832 |
| 0 | P49589 | Cysteine--tRNA ligase, cytoplasmic OS=Homo sapiens OX=9606 GN=CARS1 PE=1 SV=3 | 2 | 1 | 1 | 1 | 748 | 85.4 | 4.03 | |
| 0 | Q9HB71 | Calcyclin-binding protein OS=Homo sapiens OX=9606 GN=CACYBP PE=1 SV=2 | 8 | 1 | 1 | 1 | 228 | 26.2 | 3.99 | 14774.41895 |
| 0 | P50990 | T-complex protein 1 subunit theta OS=Homo sapiens OX=9606 GN=CCT8 PE=1 SV=4 | 2 | 1 | 1 | 1 | 548 | 59.6 | 3.96 | |
| 0 | P05109 | Protein S100-A8 OS=Homo sapiens OX=9606 GN=S100A8 PE=1 SV=1 | 12 | 1 | 2 | 1 | 93 | 10.8 | 3.93 | 22236.06836 |
| 0 | P02792 | Ferritin light chain OS=Homo sapiens OX=9606 GN=FTL PE=1 SV=2 | 9 | 1 | 1 | 1 | 175 | 20 | 3.92 | 46035.55078 |
| 0 | P41091 | Eukaryotic translation initiation factor 2 subunit 3 OS=Homo sapiens OX=9606 GN=EIF2S3 PE=1 SV=3 | 4 | 1 | 1 | 1 | 472 | 51.1 | 3.89 | 30539.26367 |
| 0 | Q9HCS7 | Pre-mRNA-splicing factor SYF1 OS=Homo sapiens OX=9606 GN=XAB2 PE=1 SV=2 | 2 | 2 | 3 | 2 | 855 | 99.9 | 3.85 | 15889.81738 |
| 0 | Q9H8G2 | Caspase activity and apoptosis inhibitor 1 OS=Homo sapiens OX=9606 GN=CAAP1 PE=1 SV=2 | 4 | 1 | 1 | 1 | 361 | 38.3 | 3.76 | |
| 0 | O60828 | Polyglutamine-binding protein 1 OS=Homo sapiens OX=9606 GN=PQBP1 PE=1 SV=1 | 22 | 2 | 2 | 2 | 265 | 30.5 | 3.75 | 26897.84766 |
| 0 | Q96ND8 | Zinc finger protein 583 OS=Homo sapiens OX=9606 GN=ZNF583 PE=2 SV=2 | 3 | 1 | 1 | 1 | 569 | 66 | 3.75 | 35526.8125 |

| | | | | | | | | | | |
|---|---|---|---|---|---|---|---|---|---|---|
| 0 | P12273 | Prolactin-inducible protein OS=Homo sapiens OX=9606 GN=PIP PE=1 SV=1 | 11 | 1 | 1 | 1 | 146 | 16.6 | 3.75 | |
| 0 | Q9UK41 | Vacuolar protein sorting-associated protein 28 homolog OS=Homo sapiens OX=9606 GN=VPS28 PE=1 SV=1 | 13 | 1 | 1 | 1 | 221 | 25.4 | 3.74 | 34641.63281 |
| 0 | O43175 | D-3-phosphoglycerate dehydrogenase OS=Homo sapiens OX=9606 GN=PHGDH PE=1 SV=4 | 3 | 1 | 1 | 1 | 533 | 56.6 | 3.73 | 12101.91992 |
| 0 | P61289 | Proteasome activator complex subunit 3 OS=Homo sapiens OX=9606 GN=PSME3 PE=1 SV=1 | 5 | 1 | 1 | 1 | 254 | 29.5 | 3.73 | 38284.17188 |
| 0 | Q15424 | Scaffold attachment factor B1 OS=Homo sapiens OX=9606 GN=SAFB PE=1 SV=4 | 2 | 1 | 1 | 1 | 915 | 102.6 | 3.68 | |
| 0 | O95670 | V-type proton ATPase subunit G 2 OS=Homo sapiens OX=9606 GN=ATP6V1G2 PE=1 SV=1 | 14 | 1 | 1 | 1 | 118 | 13.6 | 3.6 | |
| 0 | Q15007 | Pre-mRNA-splicing regulator WTAP OS=Homo sapiens OX=9606 GN=WTAP PE=1 SV=2 | 4 | 1 | 1 | 1 | 396 | 44.2 | 3.59 | 20562.36914 |
| 0 | Q7L5D6 | Golgi to ER traffic protein 4 homolog OS=Homo sapiens OX=9606 GN=GET4 PE=1 SV=1 | 4 | 1 | 1 | 1 | 327 | 36.5 | 3.59 | 35565.66406 |
| 0 | Q5RKV6 | Exosome complex component MTR3 OS=Homo sapiens OX=9606 GN=EXOSC6 PE=1 SV=1 | 10 | 1 | 1 | 1 | 272 | 28.2 | 3.59 | |
| 0 | Q9NZL9 | Methionine adenosyltransferase 2 subunit beta OS=Homo sapiens OX=9606 GN=MAT2B PE=1 SV=1 | 4 | 1 | 1 | 1 | 334 | 37.5 | 3.58 | |
| 0 | P17987 | T-complex protein 1 subunit alpha OS=Homo sapiens OX=9606 GN=TCP1 PE=1 SV=1 | 2 | 1 | 1 | 1 | 556 | 60.3 | 3.56 | 14174.04199 |
| 0 | P61927 | 60S ribosomal protein L37 OS=Homo sapiens OX=9606 GN=RPL37 PE=1 SV=2 | 28 | 3 | 3 | 3 | 97 | 11.1 | 3.56 | 79068.02832 |
| 0 | O75190 | DnaJ homolog subfamily B member 6 OS=Homo sapiens OX=9606 GN=DNAJB6 PE=1 SV=2 | 4 | 1 | 1 | 1 | 326 | 36.1 | 3.52 | |
| 0 | Q53GL7 | Protein mono-ADP-ribosyltransferase PARP10 OS=Homo sapiens OX=9606 GN=PARP10 PE=1 SV=2 | 7 | 2 | 3 | 2 | 1025 | 109.9 | 3.52 | 191658.4609 |
| 0 | P08590 | Myosin light chain 3 OS=Homo sapiens OX=9606 GN=MYL3 PE=1 SV=3 | 8 | 1 | 1 | 1 | 195 | 21.9 | 3.49 | 40938.70703 |
| 0 | Q00610 | Clathrin heavy chain 1 OS=Homo sapiens OX=9606 GN=CLTC PE=1 SV=5 | 1 | 1 | 1 | 1 | 1675 | 191.5 | 3.48 | 9704.509766 |
| 0 | Q13765 | Nascent polypeptide-associated complex subunit alpha OS=Homo sapiens OX=9606 GN=NACA PE=1 SV=1 | 7 | 1 | 1 | 1 | 215 | 23.4 | 3.48 | 11909.77832 |
| 0 | P17480 | Nucleolar transcription factor 1 OS=Homo sapiens OX=9606 GN=UBTF PE=1 SV=1 | 2 | 1 | 1 | 1 | 764 | 89.4 | 3.42 | 22044.88867 |
| 0 | Q9Y4Z0 | U6 snRNA-associated Sm-like protein LSm4 OS=Homo sapiens OX=9606 GN=LSM4 PE=1 SV=1 | 12 | 1 | 1 | 1 | 139 | 15.3 | 3.4 | 11759.08398 |
| 0 | O96019 | Actin-like protein 6A OS=Homo sapiens OX=9606 GN=ACTL6A PE=1 SV=1 | 7 | 1 | 1 | 1 | 429 | 47.4 | 3.38 | |
| 0 | O15182 | Centrin-3 OS=Homo sapiens OX=9606 GN=CETN3 PE=1 SV=2 | 8 | 1 | 1 | 1 | 167 | 19.5 | 3.36 | 15833.94629 |
| 0 | Q9BRJ7 | Tudor-interacting repair regulator protein OS=Homo sapiens OX=9606 GN=NUDT16L1 PE=1 SV=1 | 8 | 1 | 1 | 1 | 211 | 23.3 | 3.34 | 36655.72656 |
| 0 | P30153 | Serine/threonine-protein phosphatase 2A 65 kDa regulatory subunit A alpha isoform OS=Homo sapiens OX=9606 GN=PPP2R1A PE=1 SV=4 | 2 | 1 | 1 | 1 | 589 | 65.3 | 3.28 | 21015.55273 |
| 0 | Q9NPE3 | H/ACA ribonucleoprotein complex subunit 3 OS=Homo sapiens OX=9606 GN=NOP10 PE=1 SV=1 | 20 | 1 | 1 | 1 | 64 | 7.7 | 3.24 | 17436.50586 |
| 0 | Q8NAV1 | Pre-mRNA-splicing factor 38A OS=Homo sapiens OX=9606 GN=PRPF38A PE=1 SV=1 | 5 | 1 | 1 | 1 | 312 | 37.5 | 3.23 | 19410.03125 |
| 0 | Q8WXF1 | Paraspeckle component 1 OS=Homo sapiens OX=9606 GN=PSPC1 PE=1 SV=1 | 4 | 1 | 1 | 1 | 523 | 58.7 | 3.22 | 11166.03809 |
| 0 | Q9NQC3 | Reticulon-4 OS=Homo sapiens OX=9606 GN=RTN4 PE=1 SV=2 | 3 | 1 | 1 | 1 | 1192 | 129.9 | 3.2 | 19827.11328 |
| 0 | Q08554 | Desmocollin-1 OS=Homo sapiens OX=9606 GN=DSC1 PE=1 SV=2 | 2 | 1 | 1 | 1 | 894 | 99.9 | 3.17 | |
| 0 | Q5D862 | Filaggrin-2 OS=Homo sapiens OX=9606 GN=FLG2 PE=1 SV=1 | 2 | 1 | 1 | 1 | 2391 | 247.9 | 3.16 | |
| 0 | P55072 | Transitional endoplasmic reticulum ATPase OS=Homo sapiens OX=9606 GN=VCP PE=1 SV=4 | 2 | 1 | 1 | 1 | 806 | 89.3 | 3.14 | |
| 0 | Q5TCS8 | Adenylate kinase 9 OS=Homo sapiens OX=9606 GN=AK9 PE=1 SV=2 | 1 | 1 | 1 | 1 | 1911 | 221.3 | 3.14 | 6614.452148 |
| 0 | P49642 | DNA primase small subunit OS=Homo sapiens OX=9606 GN=PRIM1 PE=1 SV=1 | 6 | 1 | 1 | 1 | 420 | 49.9 | 3.12 | |
| 0 | Q13015 | Protein AF1q OS=Homo sapiens OX=9606 GN=MLLT11 PE=1 SV=1 | 32 | 1 | 1 | 1 | 90 | 10.1 | 3.1 | 8488.383789 |
| 0 | Q8TF09 | Dynein light chain roadblock-type 2 OS=Homo sapiens OX=9606 GN=DYNLRB2 PE=1 SV=1 | 17 | 1 | 1 | 1 | 96 | 10.8 | 3.09 | 20492.33203 |
| 0 | Q15738 | Sterol-4-alpha-carboxylate 3-dehydrogenase, decarboxylating OS=Homo sapiens OX=9606 GN=NSDHL PE=1 SV=2 | 4 | 1 | 1 | 1 | 373 | 41.9 | 3.08 | 30615.13086 |
| 0 | P49006 | MARCKS-related protein OS=Homo sapiens OX=9606 GN=MARCKSL1 PE=1 SV=2 | 7 | 1 | 1 | 1 | 195 | 19.5 | 3.08 | 40196.12109 |
| 0 | P20645 | Cation-dependent mannose-6-phosphate receptor OS=Homo sapiens OX=9606 GN=M6PR PE=1 SV=1 | 5 | 1 | 1 | 1 | 277 | 31 | 3.06 | 19921.04297 |
| 0 | P17980 | 26S proteasome regulatory subunit 6A OS=Homo sapiens OX=9606 GN=PSMC3 PE=1 SV=3 | 3 | 1 | 1 | 1 | 439 | 49.2 | 3.06 | 20190.12695 |
| 0 | Q9BZQ6 | ER degradation-enhancing alpha-mannosidase-like protein 3 OS=Homo sapiens OX=9606 GN=EDEM3 PE=1 SV=2 | 1 | 1 | 1 | 1 | 932 | 104.6 | 3.06 | |
| 0 | P33993 | DNA replication licensing factor MCM7 OS=Homo sapiens OX=9606 GN=MCM7 PE=1 SV=4 | 2 | 1 | 1 | 1 | 719 | 81.3 | 3.05 | 15763.29199 |
| 0 | P62857 | 40S ribosomal protein S28 OS=Homo sapiens OX=9606 GN=RPS28 PE=1 SV=1 | 17 | 1 | 1 | 1 | 69 | 7.8 | 3.02 | |
| 0 | Q01081 | Splicing factor U2AF 35 kDa subunit OS=Homo sapiens OX=9606 GN=U2AF1 PE=1 SV=3 | 6 | 1 | 1 | 1 | 240 | 27.9 | 3.01 | 14696.34082 |
| 0 | O14579 | Coatomer subunit epsilon OS=Homo sapiens OX=9606 GN=COPE PE=1 SV=3 | 7 | 1 | 1 | 1 | 308 | 34.5 | 3.01 | 28923.08594 |
| 0 | Q13547 | Histone deacetylase 1 OS=Homo sapiens OX=9606 GN=HDAC1 PE=1 SV=1 | 2 | 1 | 1 | 1 | 482 | 55.1 | 2.99 | |
| 0 | O43852 | Calumenin OS=Homo sapiens OX=9606 GN=CALU PE=1 SV=2 | 4 | 1 | 1 | 1 | 315 | 37.1 | 2.97 | |
| 0 | Q9ULR0 | Pre-mRNA-splicing factor ISY1 homolog OS=Homo sapiens OX=9606 GN=ISY1 PE=1 SV=3 | 5 | 1 | 1 | 1 | 285 | 33 | 2.94 | |
| 0 | Q04917 | 14-3-3 protein eta OS=Homo sapiens OX=9606 GN=YWHAH PE=1 SV=4 | 7 | 1 | 1 | 1 | 246 | 28.2 | 2.9 | |
| 0 | Q15027 | Arf-GAP with coiled-coil, ANK repeat and PH domain-containing protein 1 OS=Homo sapiens OX=9606 GN=ACAP1 PE=1 SV=1 | 2 | 1 | 1 | 1 | 740 | 81.5 | 2.9 | 56762.43359 |
| 0 | Q96EB1 | Elongator complex protein 4 OS=Homo sapiens OX=9606 GN=ELP4 PE=1 SV=2 | 4 | 1 | 1 | 1 | 424 | 46.6 | 2.9 | |
| 0 | Q14974 | Importin subunit beta-1 OS=Homo sapiens OX=9606 GN=KPNB1 PE=1 SV=2 | 2 | 1 | 1 | 1 | 876 | 97.1 | 2.88 | 23737.55469 |
| 0 | Q9Y2W2 | WW domain-binding protein 11 OS=Homo sapiens OX=9606 GN=WBP11 PE=1 SV=1 | 4 | 1 | 1 | 1 | 641 | 70 | 2.79 | 10997.18945 |
| 0 | Q9Y6V0 | Protein piccolo OS=Homo sapiens OX=9606 GN=PCLO PE=1 SV=5 | 0 | 1 | 1 | 1 | 5142 | 560.4 | 2.78 | 79353.96875 |
| 0 | Q9NXG2 | THUMP domain-containing protein 1 OS=Homo sapiens OX=9606 GN=THUMPD1 PE=1 SV=2 | 5 | 1 | 1 | 1 | 353 | 39.3 | 2.75 | |
| 0 | P27348 | 14-3-3 protein theta OS=Homo sapiens OX=9606 GN=YWHAQ PE=1 SV=1 | 6 | 1 | 1 | 1 | 245 | 27.7 | 2.74 | 7664.299805 |
| 0 | P46060 | Ran GTPase-activating protein 1 OS=Homo sapiens OX=9606 GN=RANGAP1 PE=1 SV=1 | 3 | 1 | 1 | 1 | 587 | 63.5 | 2.72 | 13827.38574 |
| 0 | Q9Y3F4 | Serine-threonine kinase receptor-associated protein OS=Homo sapiens OX=9606 GN=STRAP PE=1 SV=1 | 4 | 1 | 1 | 1 | 350 | 38.4 | 2.72 | |
| 0 | Q13148 | TAR DNA-binding protein 43 OS=Homo sapiens OX=9606 GN=TARDBP PE=1 SV=1 | 4 | 1 | 1 | 1 | 414 | 44.7 | 2.71 | 17985.56641 |
| 0 | Q9ULA0 | Aspartyl aminopeptidase OS=Homo sapiens OX=9606 GN=DNPEP PE=1 SV=2 | 2 | 1 | 1 | 1 | 485 | 53.4 | 2.7 | 17374.82617 |
| 0 | Q8NBS9 | Thioredoxin domain-containing protein 5 OS=Homo sapiens OX=9606 GN=TXNDC5 PE=1 SV=2 | 4 | 1 | 1 | 1 | 432 | 47.6 | 2.69 | 21167.71484 |
| 0 | Q15291 | Retinoblastoma-binding protein 5 OS=Homo sapiens OX=9606 GN=RBBP5 PE=1 SV=2 | 5 | 1 | 1 | 1 | 538 | 59.1 | 2.67 | 25404.55664 |
| 0 | Q9H773 | dCTP pyrophosphatase 1 OS=Homo sapiens OX=9606 GN=DCTPP1 PE=1 SV=1 | 13 | 1 | 1 | 1 | 170 | 18.7 | 2.67 | |
| 0 | Q13616 | Cullin-1 OS=Homo sapiens OX=9606 GN=CUL1 PE=1 SV=2 | 2 | 1 | 1 | 1 | 776 | 89.6 | 2.65 | 36376.59375 |

| | | | | | | | | | | |
|---|---|---|---|---|---|---|---|---|---|---|
| 0 | Q14444 | Caprin-1 OS=Homo sapiens OX=9606 GN=CAPRIN1 PE=1 SV=2 | 2 | 1 | 1 | 1 | 709 | 78.3 | 2.64 | 24348.92578 |
| 0 | P31025 | Lipocalin-1 OS=Homo sapiens OX=9606 GN=LCN1 PE=1 SV=1 | 6 | 1 | 1 | 1 | 176 | 19.2 | 2.62 | 12880.84082 |
| 0 | Q99832 | T-complex protein 1 subunit eta OS=Homo sapiens OX=9606 GN=CCT7 PE=1 SV=2 | 5 | 1 | 1 | 1 | 543 | 59.3 | 2.61 | 5243.024902 |
| 0 | O43148 | mRNA cap guanine-N7 methyltransferase OS=Homo sapiens OX=9606 GN=RNMT PE=1 SV=1 | 4 | 1 | 1 | 1 | 476 | 54.8 | 2.6 | |
| 0 | Q9BQA1 | Methylosome protein 50 OS=Homo sapiens OX=9606 GN=WDR77 PE=1 SV=1 | 5 | 1 | 1 | 1 | 342 | 36.7 | 2.59 | 13391.67188 |
| 0 | Q01469 | Fatty acid-binding protein 5 OS=Homo sapiens OX=9606 GN=FABP5 PE=1 SV=3 | 7 | 1 | 1 | 1 | 135 | 15.2 | 2.56 | 21253.89063 |
| 0 | Q7Z3C6 | Autophagy-related protein 9A OS=Homo sapiens OX=9606 GN=ATG9A PE=1 SV=3 | 5 | 2 | 2 | 2 | 839 | 94.4 | 2.55 | 45683.30078 |
| 0 | P04843 | Dolichyl-diphosphooligosaccharide--protein glycosyltransferase subunit 1 OS=Homo sapiens OX=9606 GN=RPN1 PE=1 SV=1 | 3 | 1 | 1 | 1 | 607 | 68.5 | 2.55 | 11145.87793 |
| 0 | O14964 | Hepatocyte growth factor-regulated tyrosine kinase substrate OS=Homo sapiens OX=9606 GN=HGS PE=1 SV=1 | 1 | 1 | 1 | 1 | 777 | 86.1 | 2.54 | 86253.57813 |
| 0 | P61160 | Actin-related protein 2 OS=Homo sapiens OX=9606 GN=ACTR2 PE=1 SV=1 | 3 | 1 | 1 | 1 | 394 | 44.7 | 2.53 | 14528.49609 |
| 0 | P49756 | RNA-binding protein 25 OS=Homo sapiens OX=9606 GN=RBM25 PE=1 SV=3 | 2 | 1 | 1 | 1 | 843 | 100.1 | 2.52 | |
| 0 | Q15651 | High mobility group nucleosome-binding domain-containing protein 3 OS=Homo sapiens OX=9606 GN=HMGN3 PE=1 SV=2 | 15 | 1 | 1 | 1 | 99 | 10.7 | 2.51 | 33767.71875 |
| 0 | P62195 | 26S proteasome regulatory subunit 8 OS=Homo sapiens OX=9606 GN=PSMC5 PE=1 SV=1 | 3 | 1 | 1 | 1 | 406 | 45.6 | 2.49 | |
| 0 | D6REC4 | Cilia- and flagella-associated protein 99 OS=Homo sapiens OX=9606 GN=CFAP99 PE=3 SV=1 | 2 | 1 | 1 | 1 | 459 | 52.3 | 2.43 | 61723.68359 |
| 0 | Q96AB3 | Isochorismatase domain-containing protein 2 OS=Homo sapiens OX=9606 GN=ISOC2 PE=1 SV=1 | 16 | 1 | 1 | 1 | 205 | 22.3 | 2.42 | |
| 0 | P15924 | Desmoplakin OS=Homo sapiens OX=9606 GN=DSP PE=1 SV=3 | 0 | 1 | 1 | 1 | 2871 | 331.6 | 2.42 | |
| 0 | P26641 | Elongation factor 1-gamma OS=Homo sapiens OX=9606 GN=EEF1G PE=1 SV=3 | 2 | 1 | 1 | 1 | 437 | 50.1 | 2.42 | 41741.83203 |
| 0 | Q96DI7 | U5 small nuclear ribonucleoprotein 40 kDa protein OS=Homo sapiens OX=9606 GN=SNRNP40 PE=1 SV=1 | 3 | 1 | 1 | 1 | 357 | 39.3 | 2.41 | 31852.09766 |
| 0 | P41223 | Protein BUD31 homolog OS=Homo sapiens OX=9606 GN=BUD31 PE=1 SV=2 | 7 | 1 | 1 | 1 | 144 | 17 | 2.38 | 26624.95117 |
| 0 | Q9BZJ0 | Crooked neck-like protein 1 OS=Homo sapiens OX=9606 GN=CRNKL1 PE=1 SV=4 | 2 | 1 | 1 | 1 | 848 | 100.4 | 2.37 | |
| 0 | O43166 | Signal-induced proliferation-associated 1-like protein 1 OS=Homo sapiens OX=9606 GN=SIPA1L1 PE=1 SV=4 | 2 | 1 | 1 | 1 | 1804 | 199.9 | 2.36 | |
| 0 | P52565 | Rho GDP-dissociation inhibitor 1 OS=Homo sapiens OX=9606 GN=ARHGDIA PE=1 SV=3 | 16 | 1 | 1 | 1 | 204 | 23.2 | 2.36 | 15034.0752 |
| 0 | P49773 | Histidine triad nucleotide-binding protein 1 OS=Homo sapiens OX=9606 GN=HINT1 PE=1 SV=2 | 11 | 1 | 1 | 1 | 126 | 13.8 | 2.35 | 45425.73047 |
| 0 | Q8IZP0 | Abl interactor 1 OS=Homo sapiens OX=9606 GN=ABI1 PE=1 SV=4 | 2 | 1 | 1 | 1 | 508 | 55 | 2.35 | 12393.6377 |
| 0 | P62891 | 60S ribosomal protein L39 OS=Homo sapiens OX=9606 GN=RPL39 PE=1 SV=2 | 25 | 1 | 3 | 1 | 51 | 6.4 | 2.33 | 93591.2168 |
| 0 | P62995 | Transformer-2 protein homolog beta OS=Homo sapiens OX=9606 GN=TRA2B PE=1 SV=1 | 3 | 1 | 1 | 1 | 288 | 33.6 | 2.3 | 165168.6094 |
| 0 | Q86SS6 | Synaptotagmin-9 OS=Homo sapiens OX=9606 GN=SYT9 PE=1 SV=1 | 4 | 1 | 1 | 1 | 491 | 56.2 | 2.29 | |
| 0 | Q9UGM3 | Deleted in malignant brain tumors 1 protein OS=Homo sapiens OX=9606 GN=DMBT1 PE=1 SV=2 | 7 | 1 | 1 | 1 | 2413 | 260.6 | 2.27 | |
| 0 | O60906 | Sphingomyelin phosphodiesterase 2 OS=Homo sapiens OX=9606 GN=SMPD2 PE=1 SV=2 | 3 | 1 | 1 | 1 | 423 | 47.6 | 2.27 | 34612.92969 |
| 0 | Q86WX3 | Active regulator of SIRT1 OS=Homo sapiens OX=9606 GN=RPS19BP1 PE=1 SV=1 | 15 | 1 | 1 | 1 | 136 | 15.4 | 2.25 | |
| 0 | Q00765 | Receptor expression-enhancing protein 5 OS=Homo sapiens OX=9606 GN=REEP5 PE=1 SV=3 | 6 | 1 | 1 | 1 | 189 | 21.5 | 2.24 | 13575.86035 |
| 0 | Q9Y3C1 | Nucleolar protein 16 OS=Homo sapiens OX=9606 GN=NOP16 PE=1 SV=2 | 4 | 1 | 1 | 1 | 178 | 21.2 | 2.22 | |
| 0 | P30041 | Peroxiredoxin-6 OS=Homo sapiens OX=9606 GN=PRDX6 PE=1 SV=3 | 5 | 1 | 1 | 1 | 224 | 25 | 2.18 | 25143.65234 |
| 0 | Q5VZK9 | F-actin-uncapping protein LRRC16A OS=Homo sapiens OX=9606 GN=CARMIL1 PE=1 SV=1 | 1 | 1 | 1 | 1 | 1371 | 151.5 | 2.17 | |
| 0 | Q16378 | Proline-rich protein 4 OS=Homo sapiens OX=9606 GN=PRR4 PE=1 SV=3 | 10 | 1 | 1 | 1 | 134 | 15.1 | 2.17 | 19136.00195 |
| 0 | Q9UN86 | Ras GTPase-activating protein-binding protein 2 OS=Homo sapiens OX=9606 GN=G3BP2 PE=1 SV=2 | 3 | 1 | 1 | 1 | 482 | 54.1 | 2.16 | 34716.70313 |
| 0 | O94864 | STAGA complex 65 subunit gamma OS=Homo sapiens OX=9606 GN=SUPT7L PE=1 SV=1 | 3 | 1 | 1 | 1 | 414 | 46.2 | 2.16 | 13596.44434 |
| 0 | O75762 | Transient receptor potential cation channel subfamily A member 1 OS=Homo sapiens OX=9606 GN=TRPA1 PE=1 SV=3 | 2 | 1 | 1 | 1 | 1119 | 127.4 | 2.15 | |
| 0 | A5D8V6 | Vacuolar protein sorting-associated protein 37C OS=Homo sapiens OX=9606 GN=VPS37C PE=1 SV=2 | 3 | 1 | 1 | 1 | 355 | 38.6 | 2.15 | 17690.69141 |
| 0 | Q9NWB6 | Arginine and glutamate-rich protein 1 OS=Homo sapiens OX=9606 GN=ARGLU1 PE=1 SV=1 | 3 | 1 | 1 | 1 | 273 | 33.2 | 2.12 | 23746.22461 |
| 0 | Q99961 | Endophilin-A2 OS=Homo sapiens OX=9606 GN=SH3GL1 PE=1 SV=1 | 3 | 1 | 1 | 1 | 368 | 41.5 | 2.11 | 31399.4043 |
| 0 | P04075 | Fructose-bisphosphate aldolase A OS=Homo sapiens OX=9606 GN=ALDOA PE=1 SV=2 | 4 | 1 | 1 | 1 | 364 | 39.4 | 2.1 | |
| 0 | Q6UXN9 | WD repeat-containing protein 82 OS=Homo sapiens OX=9606 GN=WDR82 PE=1 SV=1 | 4 | 1 | 1 | 1 | 313 | 35.1 | 2.1 | |
| 0 | P78406 | mRNA export factor OS=Homo sapiens OX=9606 GN=RAE1 PE=1 SV=1 | 2 | 1 | 1 | 1 | 368 | 40.9 | 2.1 | 19368.78906 |
| 0 | P22087 | rRNA 2'-O-methyltransferase fibrillarin OS=Homo sapiens OX=9606 GN=FBL PE=1 SV=2 | 4 | 1 | 1 | 1 | 321 | 33.8 | 2.09 | 20962.12305 |
| 0 | P11802 | Cyclin-dependent kinase 4 OS=Homo sapiens OX=9606 GN=CDK4 PE=1 SV=2 | 5 | 1 | 1 | 1 | 303 | 33.7 | 2.09 | |
| 0 | O94906 | Pre-mRNA-processing factor 6 OS=Homo sapiens OX=9606 GN=PRPF6 PE=1 SV=1 | 1 | 1 | 1 | 1 | 941 | 106.9 | 2.06 | |
| 0 | P07477 | Trypsin-1 OS=Homo sapiens OX=9606 GN=PRSS1 PE=1 SV=1 | 2 | 1 | 1 | 1 | 247 | 26.5 | 2.05 | 66945.52344 |
| 0 | Q9NW64 | Pre-mRNA-splicing factor RBM22 OS=Homo sapiens OX=9606 GN=RBM22 PE=1 SV=1 | 7 | 2 | 2 | 2 | 420 | 46.9 | 2.05 | 40823.77148 |
| 0 | O00471 | Exocyst complex component 5 OS=Homo sapiens OX=9606 GN=EXOC5 PE=1 SV=1 | 3 | 1 | 1 | 1 | 708 | 81.8 | 2.05 | 53661.98828 |
| 0 | Q16181 | Septin-7 OS=Homo sapiens OX=9606 GN=SEPTIN7 PE=1 SV=2 | 2 | 1 | 1 | 1 | 437 | 50.6 | 2.05 | 9046.702148 |
| 0 | Q8WXW3 | Progesterone-induced-blocking factor 1 OS=Homo sapiens OX=9606 GN=PIBF1 PE=1 SV=2 | 2 | 1 | 1 | 1 | 757 | 89.8 | 2.04 | 17856.55469 |
| 0 | P62310 | U6 snRNA-associated Sm-like protein LSm3 OS=Homo sapiens OX=9606 GN=LSM3 PE=1 SV=2 | 14 | 1 | 1 | 1 | 102 | 11.8 | 2.04 | 35343.89453 |
| 0 | O43660 | Pleiotropic regulator 1 OS=Homo sapiens OX=9606 GN=PLRG1 PE=1 SV=1 | 2 | 1 | 1 | 1 | 514 | 57.2 | 2.04 | 17444.5293 |
| 0 | P06493 | Cyclin-dependent kinase 1 OS=Homo sapiens OX=9606 GN=CDK1 PE=1 SV=3 | 3 | 1 | 1 | 1 | 297 | 34.1 | 2.03 | 21109.83398 |
| 0 | P30825 | High affinity cationic amino acid transporter 1 OS=Homo sapiens OX=9606 GN=SLC7A1 PE=1 SV=1 | 2 | 1 | 1 | 1 | 629 | 67.6 | 2.02 | 13138.04102 |
| 0 | Q96S66 | Chloride channel CLIC-like protein 1 OS=Homo sapiens OX=9606 GN=CLCC1 PE=1 SV=1 | 2 | 1 | 1 | 1 | 551 | 62 | 2.02 | 47608.03125 |
| 0 | P05114 | Non-histone chromosomal protein HMG-14 OS=Homo sapiens OX=9606 GN=HMGN1 PE=1 SV=3 | 14 | 1 | 2 | 1 | 100 | 10.7 | 1.98 | 33316.70703 |
| 0 | Q7Z478 | ATP-dependent RNA helicase DHX29 OS=Homo sapiens OX=9606 GN=DHX29 PE=1 SV=2 | 0 | 1 | 1 | 1 | 1369 | 155.1 | 1.96 | |
| 0 | Q5JTH9 | RRP12-like protein OS=Homo sapiens OX=9606 GN=RRP12 PE=1 SV=2 | 1 | 1 | 1 | 1 | 1297 | 143.6 | 1.94 | 16688 |
| 0 | Q66PJ3 | ADP-ribosylation factor-like protein 6-interacting protein 4 OS=Homo sapiens OX=9606 GN=ARL6IP4 PE=1 SV=2 | 4 | 1 | 1 | 1 | 421 | 44.9 | 1.94 | 16783.28125 |
| 0 | Q9UNE7 | E3 ubiquitin-protein ligase CHIP OS=Homo sapiens OX=9606 GN=STUB1 PE=1 SV=2 | 3 | 1 | 1 | 1 | 303 | 34.8 | 1.94 | |

| | | | | | | | | | | |
|---|---|---|---|---|---|---|---|---|---|---|
| 0 | P78371 | T-complex protein 1 subunit beta OS=Homo sapiens OX=9606 GN=CCT2 PE=1 SV=4 | 2 | 1 | 1 | 1 | 535 | 57.5 | 1.93 | |
| 0 | Q13123 | Protein Red OS=Homo sapiens OX=9606 GN=IK PE=1 SV=3 | 1 | 1 | 1 | 1 | 557 | 65.6 | 1.92 | 465679.125 |
| 0 | Q9BQG0 | Myb-binding protein 1A OS=Homo sapiens OX=9606 GN=MYBBP1A PE=1 SV=2 | 1 | 2 | 2 | 2 | 1328 | 148.8 | 1.92 | 36762.69043 |
| 0 | P35568 | Insulin receptor substrate 1 OS=Homo sapiens OX=9606 GN=IRS1 PE=1 SV=1 | 1 | 1 | 1 | 1 | 1242 | 131.5 | 1.9 | 13436.08301 |
| 0 | Q9HB58 | Sp110 nuclear body protein OS=Homo sapiens OX=9606 GN=SP110 PE=1 SV=5 | 1 | 1 | 1 | 1 | 689 | 78.3 | 1.86 | 46269.33203 |
| 0 | P62491 | Ras-related protein Rab-11A OS=Homo sapiens OX=9606 GN=RAB11A PE=1 SV=3 | 5 | 1 | 1 | 1 | 216 | 24.4 | 1.86 | |
| 0 | O43172 | U4/U6 small nuclear ribonucleoprotein Prp4 OS=Homo sapiens OX=9606 GN=PRPF4 PE=1 SV=2 | 2 | 1 | 1 | 1 | 522 | 58.4 | 1.83 | |
| 0 | Q96I25 | Splicing factor 45 OS=Homo sapiens OX=9606 GN=RBM17 PE=1 SV=1 | 2 | 1 | 1 | 1 | 401 | 44.9 | 1.79 | |
| 0 | P05141 | ADP/ATP translocase 2 OS=Homo sapiens OX=9606 GN=SLC25A5 PE=1 SV=7 | 8 | 1 | 1 | 1 | 298 | 32.8 | 1.79 | 9017.486328 |
| 0 | Q2TAY7 | WD40 repeat-containing protein SMU1 OS=Homo sapiens OX=9606 GN=SMU1 PE=1 SV=2 | 2 | 1 | 1 | 1 | 513 | 57.5 | 1.76 | 55711.37891 |
| 0 | P37837 | Transaldolase OS=Homo sapiens OX=9606 GN=TALDO1 PE=1 SV=2 | 2 | 1 | 1 | 1 | 337 | 37.5 | 1.73 | 36237.05859 |
| 0 | P78345 | Ribonuclease P protein subunit p38 OS=Homo sapiens OX=9606 GN=RPP38 PE=1 SV=2 | 4 | 1 | 1 | 1 | 283 | 31.8 | 1.72 | |
| 0 | Q9Y314 | Nitric oxide synthase-interacting protein OS=Homo sapiens OX=9606 GN=NOSIP PE=1 SV=1 | 3 | 1 | 1 | 1 | 301 | 33.2 | 1.7 | |
| 0 | Q96IK0 | Transmembrane protein 101 OS=Homo sapiens OX=9606 GN=TMEM101 PE=1 SV=1 | 4 | 1 | 1 | 1 | 257 | 28.8 | 1.69 | 28125.02148 |
| 0 | Q13601 | KRR1 small subunit processome component homolog OS=Homo sapiens OX=9606 GN=KRR1 PE=1 SV=4 | 4 | 1 | 1 | 1 | 381 | 43.6 | 1.68 | |
| 0 | P84098 | 60S ribosomal protein L19 OS=Homo sapiens OX=9606 GN=RPL19 PE=1 SV=1 | 5 | 1 | 1 | 1 | 196 | 23.5 | 1.66 | |
| 0 | Q99848 | Probable rRNA-processing protein EBP2 OS=Homo sapiens OX=9606 GN=EBNA1BP2 PE=1 SV=2 | 3 | 1 | 1 | 1 | 306 | 34.8 | 1.6 | |
| 0 | O75396 | Vesicle-trafficking protein SEC22b OS=Homo sapiens OX=9606 GN=SEC22B PE=1 SV=4 | 10 | 1 | 1 | 1 | 215 | 24.6 | 0 | |
| 0 | Q8N5L8 | Ribonuclease P protein subunit p25-like protein OS=Homo sapiens OX=9606 GN=RPP25L PE=1 SV=1 | 6 | 1 | 1 | 1 | 163 | 17.6 | 0 | |
| 0 | Q9ULI4 | Kinesin-like protein KIF26A OS=Homo sapiens OX=9606 GN=KIF26A PE=1 SV=3 | 2 | 1 | 1 | 1 | 1882 | 194.5 | 0 | 7714.280762 |
| 0 | Q6XYB7 | Transcription factor LBX2 OS=Homo sapiens OX=9606 GN=LBX2 PE=1 SV=1 | 8 | 1 | 1 | 1 | 198 | 21.5 | 0 | |
| 0 | Q9UPR0 | Inactive phospholipase C-like protein 2 OS=Homo sapiens OX=9606 GN=PLCL2 PE=1 SV=2 | 3 | 1 | 1 | 1 | 1127 | 125.8 | 0 | 74536.40625 |
| 0 | Q9P219 | Protein Daple OS=Homo sapiens OX=9606 GN=CCDC88C PE=1 SV=3 | 0 | 1 | 1 | 1 | 2028 | 228.1 | 0 | 22210.19727 |
| 0 | Q6XD76 | Achaete-scute homolog 4 OS=Homo sapiens OX=9606 GN=ASCL4 PE=1 SV=1 | 7 | 1 | 1 | 1 | 172 | 19.2 | 0 | 10043.80273 |
| 0 | P21281 | V-type proton ATPase subunit B, brain isoform OS=Homo sapiens OX=9606 GN=ATP6V1B2 PE=1 SV=3 | 5 | 1 | 1 | 1 | 511 | 56.5 | 0 | 10965.56738 |
| 0 | Q9UBN4 | Short transient receptor potential channel 4 OS=Homo sapiens OX=9606 GN=TRPC4 PE=1 SV=1 | 1 | 1 | 1 | 1 | 977 | 112 | 0 | 85405.21094 |
| 0 | Q9NWU2 | Glucose-induced degradation protein 8 homolog OS=Homo sapiens OX=9606 GN=GID8 PE=1 SV=1 | 11 | 1 | 1 | 1 | 228 | 26.7 | 0 | 14357.53418 |
| 0 | Q9UNS2 | COP9 signalosome complex subunit 3 OS=Homo sapiens OX=9606 GN=COPS3 PE=1 SV=3 | 4 | 1 | 1 | 1 | 423 | 47.8 | 0 | |
| 0 | Q8WWM7 | Ataxin-2-like protein OS=Homo sapiens OX=9606 GN=ATXN2L PE=1 SV=2 | 2 | 1 | 1 | 1 | 1075 | 113.3 | 0 | |
| 0 | P08195 | 4F2 cell-surface antigen heavy chain OS=Homo sapiens OX=9606 GN=SLC3A2 PE=1 SV=3 | 2 | 1 | 1 | 1 | 630 | 68 | 0 | |
| 0 | Q96L91 | E1A-binding protein p400 OS=Homo sapiens OX=9606 GN=EP400 PE=1 SV=4 | 1 | 1 | 1 | 1 | 3159 | 343.3 | 0 | |
